# Site-resolved energetic information from HX/MS experiments

**DOI:** 10.1101/2024.08.04.606547

**Authors:** Chenlin Lu, Malcolm L. Wells, Andrew Reckers, Anum Glasgow

**Author notes:** These authors made equal contributions (Chenlin Lu, Malcolm L. Wells).

## Abstract

High-resolution energetic information about protein conformational ensembles is essential for understanding protein function, yet remains challenging to obtain. We present PIGEON-FEATHER, a method for calculating ensemble free energies of opening (ΔG_op_) at single- or near-single-amino acid resolution for proteins of all sizes from hydrogen exchange/mass spectrometry (HX/MS) data. PIGEON-FEATHER disambiguates and reconstructs all experimentally measured HX/MS isotopic mass envelopes using a Bayesian Monte Carlo sampling approach. We applied PIGEON-FEATHER to reveal how *E. coli* and human dihydrofolate reductases (ecDHFR, hDHFR) have evolved distinct ensembles. We show how two competitive inhibitors bind these orthologs differently, solving the longstanding mystery of why both therapeutic molecules inhibit hDHFR, but only one inhibits ecDHFR. Extending PIGEON-FEATHER to a large protein-DNA complex, we mapped ligand-induced ensemble reweighting in the *E. coli lac* repressor to describe the functional switching mechanism crucial for transcriptional regulation.

---

Proteins perform biological functions as conformational ensembles: sets of interconverting conformational states with different populations and energies.^1–3^ More energetically favorable conformational states are more probable, so shifts in local energetic relationships drive shifts in population within the ensemble. Molecular perturbations like ligand binding cause proteins to reweight their ensembles, regulating their functions. Disease-causing mutations disrupt ensembles. Because of this key relationship between a protein’s ensemble and function, mapping the conformational ensembles of proteins is crucial for accurately describing and predicting their behavior.

However, biophysical techniques are limited in their capacity to measure ensemble energies, especially at high resolution. X-ray crystallography and cryo-electron microscopy (cryo-EM) capture only the lowest-energy conformational states of proteins. Nuclear magnetic resonance (NMR) spectroscopy monitors shifts among conformational states but is limited by experimental throughput and protein size. Molecular dynamics (MD) simulations can reveal the details of the earliest stages in a conformational transition, but are constrained by simulation timescales, inaccuracies in force fields, and the availability of structural data.

We set out to measure amino acid residue-level ensemble energies using hydrogen exchange with mass spectrometry (HX/MS). With the advent of integrated robotics^4,5^ and accurate protein structure prediction methods^6,7^, HX/MS presents a powerful strategy for probing protein ensembles in solution. HX/MS measures the rate of exchange of labile hydrogen atoms with deuterium in a protein’s backbone amide groups to report changes in its ensemble at the peptide level (Fig. 1a).^8^ For hydrogen exchange to occur in a folded protein at equilibrium (EX2 conditions, see section 1 of the SI), the protein must transiently expose its backbone amide hydrogens to solvent, which incurs an energetic cost for each exchangeable site: the free energy of opening, or ΔG_op_ (Fig. 1b).^9^ The ΔG_op_ is an ensemble energy derived from the hydrogen exchange rate.

**Figure 1.**
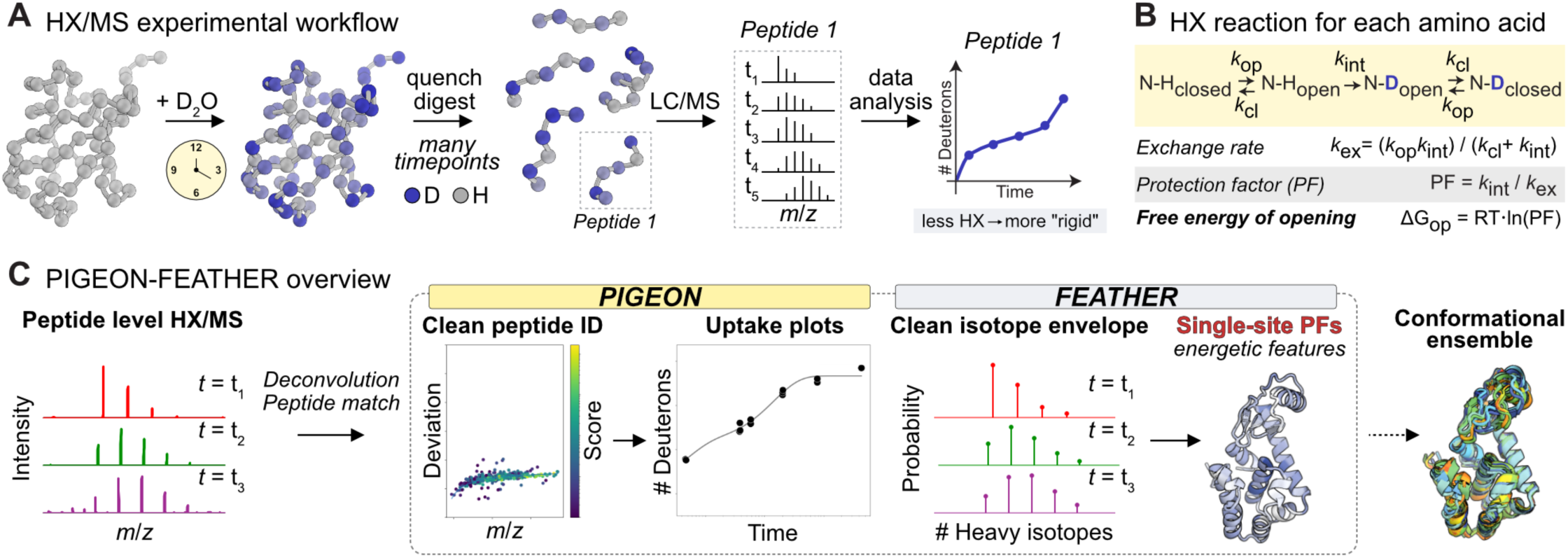
PIGEON-FEATHER overview. **A.** HX/MS workflow. Increased H-D exchange over the timecourse for a peptide in one functional state compared to another state indicates less involvement in backbone amide hydrogen bond interactions. **B.** Derivation of free energies of opening (ΔG_op_) for protein ensembles from protection factors (PFs), calculated from H-D exchange rates. **C.** Summary of the PIGEON-FEATHER method to disambiguate peptides from HX/MS data and calculate single-site ΔG_op_ values as features of the protein ensemble.

Because of a lack of methods to determine exchange rates at single residue resolution, HX/MS data are typically interpreted as relative differences in total hydrogen exchange between states (*e.g.*, apo vs. holo proteins) for selected peptides. This approach is flawed for three reasons. First, total hydrogen exchange is biased by the experimental timecourse, because exchange half-lives can span years but the experiment is performed within hours.^10^ Second, the effects of protein sequence on exchange rates are ignored. The ΔG_op_ depends on the intrinsic rate of exchange for each residue, or how it exchanges in an unstructured polypeptide (Fig. 1b), which depends on its own and its neighbors’ identities^11^; but the peptide resolution of HX/MS data precludes assignments of residue-specific rates. Third, the manner of curating, combining, and visualizing the peptide-level total hydrogen exchange data influences how we understand protein ensembles.^12,13^

We therefore developed PIGEON-FEATHER *(Peptide ID Generation for Exchange Of Nuclei - Free Energy Assignment Through Hydrogen Exchange Rates)*, an HX/MS analysis method to assign ΔG_op_ for almost all the residues in a protein (Fig. 1c). PIGEON-FEATHER disambiguates peptides based on tandem MS fragment coverage and fits hydrogen exchange rates via Bayesian sampling to reconstruct full isotopic mass envelopes, capitalizing on information-rich raw HX/MS data for ΔG_op_ determination. Benchmarking shows that PIGEON-FEATHER calculates exchange rates with near-perfect accuracy, consistently outperforming existing methods. We applied PIGEON-FEATHER to bacterial and human enzyme orthologs to quantify how evolutionary divergence in ΔG_op_ enables catalysis in different organisms. We measured how two competitive inhibitor therapeutics differentially impact the bacterial and human enzyme orthologs to discern why only one of them is efficacious in the human enzyme, yet both inhibit the bacterial enzyme. Finally, we used PIGEON-FEATHER to connect ligand-induced allosteric changes in the regulatory domain of a large transcription factor with local unfolding in its DNA-binding domain.

## Results

### Peptide disambiguation using PIGEON

In hydrogen-deuterium (H-D) exchange/MS experiments, proteins are nonspecifically proteolyzed over an exchange timecourse and measured for D uptake. We first perform a tandem MS experiment on the undeuterated protein and match the monoisotopic masses and *y*- and *b*- ion fragments to theoretical masses based on the protein sequence. The observed mass, charge, and chromatographic retention time are used to identify the corresponding peaks in the deuterated sample spectra. However, this is often inadequate to uniquely specify one peptide (Extended Data Fig. 1a-d).^14–16^

PIGEON enables accurate peptide identification (Extended Data Fig. 1e). First, PIGEON corrects systematic calibration errors (Fig. S1a-c). Next, PIGEON chooses a consensus match for each peptide, using the match with the best fragment ion coverage when there are multiple matches (Fig. S1d). Finally, PIGEON identifies remaining duplicated matches by monoisotopic *m*/*z* and retention time, and discards those without fragment support (Extended Data Fig. 1e, iii, g). Co-eluting peptides where both matches have unique fragment support meeting the significance threshold may be retained or discarded per user-selectable ‘keep’ and ‘drop’ modes (Extended Data Fig. 1e, iv, 1h). The remaining assignments form a peptide pool for HX analysis using FEATHER (Extended Data Fig. 1e, v, 1h).

We evaluated how PIGEON impacts peptide assignments for HX/MS on *E. coli* dihydrofolate reductase (ecDHFR). There are 2,107 possible peptides of length 3-15 amino acids in the ecDHFR sequence, many of which are degenerate in mass, especially when considering instrument sensitivity limitations (Fig. 2a, b). PIGEON uses a 0.1 Da/e threshold for identifying duplicates to account for 10-100-fold lower MS sensitivity. We assessed the impact of each filtering step on the number of peptides (Extended Data Fig. 1h, Table S1). In one case, we discarded 44 of 276 initial peptide assignments as duplicates.

**Figure 2.**
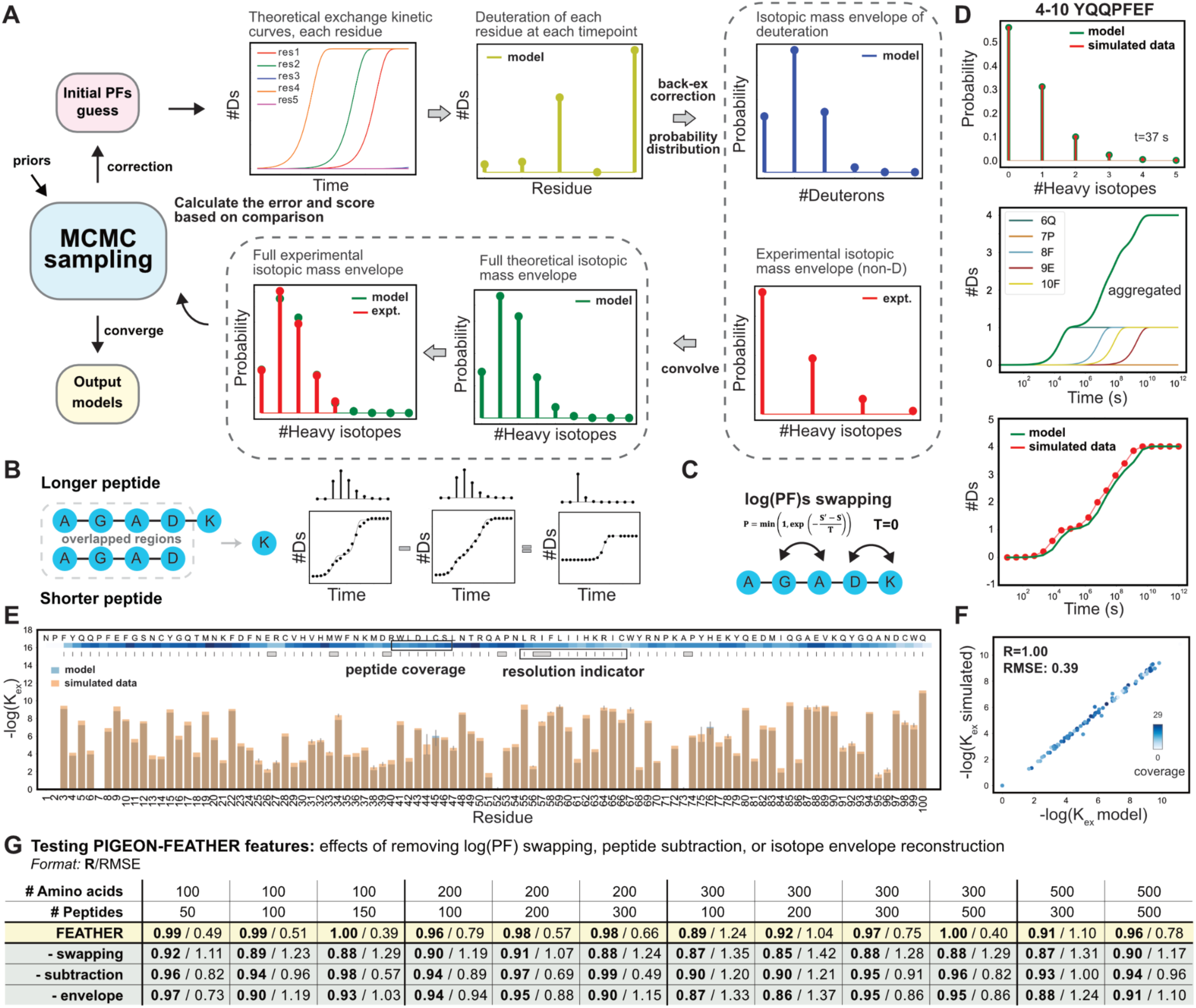
MS envelope fitting with FEATHER. **A.** Isotopic mass envelope reconstruction incorporates a Markov Chain Monte Carlo (MCMC) sampling framework. **B.** Peptide subtraction to generate data at a higher resolution. **C.** During MCMC sampling, each residue has a chance to swap its log(PF) with a neighboring residue. **D.** Representative fitting results on a simulated dataset: the reconstructed envelope vs. the simulated counterpart, alongside FEATHER-calculated D uptake kinetic curves for individual residues. These are aggregated to produce a peptide uptake plot and juxtaposed with centroid-level data. **E.** Overlay of the model-derived exchange rates with simulated data. The standard deviation of exchange rates within a cluster aggregated across multiple bootstrapping runs is represented as an error bar (see Methods). **F.** Correlation between the modeled and simulated kinetic exchange rates. Points represent individual residues colored by peptide coverage. **G.** FEATHER benchmarks on synthetic data.

### ΔG_op_ assignment using FEATHER

Following peptide disambiguation from the tandem MS experiment using PIGEON, HX/MS data can be analyzed using FEATHER. FEATHER is a Bayesian method to assign ΔG_op_ (Fig. 2). While there are several methods to calculate centroid values from isotopic mass envelopes to fit ΔG_op_^17–19^, the degenerate nature of MS centroids causes considerable information loss, as many envelopes can yield identical centroid values.^16^ By contrast, direct envelope fitting introduces additional constraints, thereby minimizing calculation uncertainties and enhancing accuracy.^20^ Building on a centroid-based Bayesian method^17^, we implemented FEATHER to include sampling improvements and isotopic mass envelope fitting (Fig. 2a).^20^

First, FEATHER initiates full isotopic mass envelope reconstruction for each peptide at each timepoint with initial ΔG_op_ guesses for each residue. The ΔG_op_ values are refined through iterative Markov Chain Monte Carlo (MCMC) sampling. ΔG_op_ are explored on discrete grids by the corresponding protection factors (PFs) on logarithmic scale within a range of 0 to 14 (Fig. 1b). The full theoretical kinetic exchange curves for each residue are built from single exponential functions given the ΔG_op_ value. D-to-H back-exchange, an effect of the chromatography steps in HX/MS, is corrected at each timepoint, followed by a binomial probability distribution calculation to determine the isotopic envelope of deuteration. The isotopic envelope of deuteration and experimental isotopic envelope are then convolved to obtain a theoretical envelope. Then, the likelihood function is calculated based on the comparison between the theoretical and observed MS data (details in Methods). FEATHER continues MCMC sampling until the model converges on a posterior distribution for ΔG_op_ across all residues (Fig. 2a).

In addition to MS reconstruction, three features improve FEATHER performance: peptide subtraction, swap-sampling, and structural priors. First, peptides with N/C terminal overlaps are subtracted from one another at the mass envelope level to produce mini-peptides at an enhanced resolution as a data augmentation for ΔG_op_ fitting (Fig. 2b). We also implemented an explicit log(PF)s swapping step to enhance the sampling (Fig. 2c). During each swapping step, a subset of sampled residues may swap their log(PF) values with adjacent amino acids. The final ΔG_op_ values are derived by combining results from multiple bootstrap replicates. The standard deviation of ΔG_op_ within a cluster aggregated across multiple bootstrapping runs is used as a measurement of the uncertainty. We also implemented two optional Bayesian priors: a *structural prior*, which leverages a protein structure to bias sampling, and an *uptake prior*, which compares calculated exchange rates with centroid-level uptake (see Methods).

### Simulated data benchmarks

To assess FEATHER accuracy, we prepared simulated datasets by assigning exchange rates to all residues in synthetic proteins of varied sequences and lengths (Tables S2, S3). The proteins were decomposed into peptides with sequence coverage, noise, and overlap reminiscent of true HX/MS data (Extended Data Fig. 2a, Methods). We used FEATHER to calculate the exchange rates and ΔG_op_ values for each residue in these proteins (Fig. 2d, e). FEATHER performance was measured as the correlation coefficient of the true vs. calculated log(PF) (R) and root-mean-square error (RMSE) (Fig. 2f). FEATHER achieved R values near 1 for proteins of all sizes (Fig. 2g).

We found that peptide subtraction, swap-sampling, and MS reconstruction contribute synergistically to FEATHER performance (Fig. 2g, Table S4); the benefit of including all three features exceeds the sum of their individual contributions (Table S4). Notably, FEATHER benefits more from increased data volume when using mass envelope data rather than centroids, highlighting further advantages of envelope fitting. Further, removing half of the timepoints, removing timepoints beyond 10^6^ seconds, and noising the MS peaks did not substantially reduce accuracy (Extended Data Fig. 2b, Tables S3, S5).

To compare FEATHER to other methods^17–21^, we also analyzed our optimal synthetic dataset using BayesianHDX^17^, PyHDX^18^, ExPfact^19^, and HDsite^20^ (Extended Data Fig. 2c, Table S6). FEATHER considerably outperforms each method. Although FEATHER builds on its sampling method, BayesianHDX fits centroids and samples less efficiently, leading to diminished accuracy. HDsite was the first method to reconstruct full mass envelopes, but its iterative fitting approach and scoring function reduces its performance. Due to its incompatibility with dense data and weeks-long compute time, we could only test HDsite on one small dataset. Overall, FEATHER combines the best features of prior methods with sampling and architecture improvements for unprecedentedly accurate ΔG_op_ calculations.

### PIGEON-FEATHER determines single-site ΔG_op_ values for ecDHFR

As an essential enzyme in nucleotide synthesis, ecDHFR is a clinical target for treating infections.^22^ After confirming its activity (Fig. S2), we tested the performance of PIGEON-FEATHER on ecDHFR (Fig. 3a-d). Comparing the fitted model with the experimental mass spectra and centroid-level D uptake demonstrates that FEATHER can accurately fit real HX/MS data (Fig. 3a) with low error (Fig. 3b). All D uptake plots and fitted models are in the SI files.

**Figure 3.**
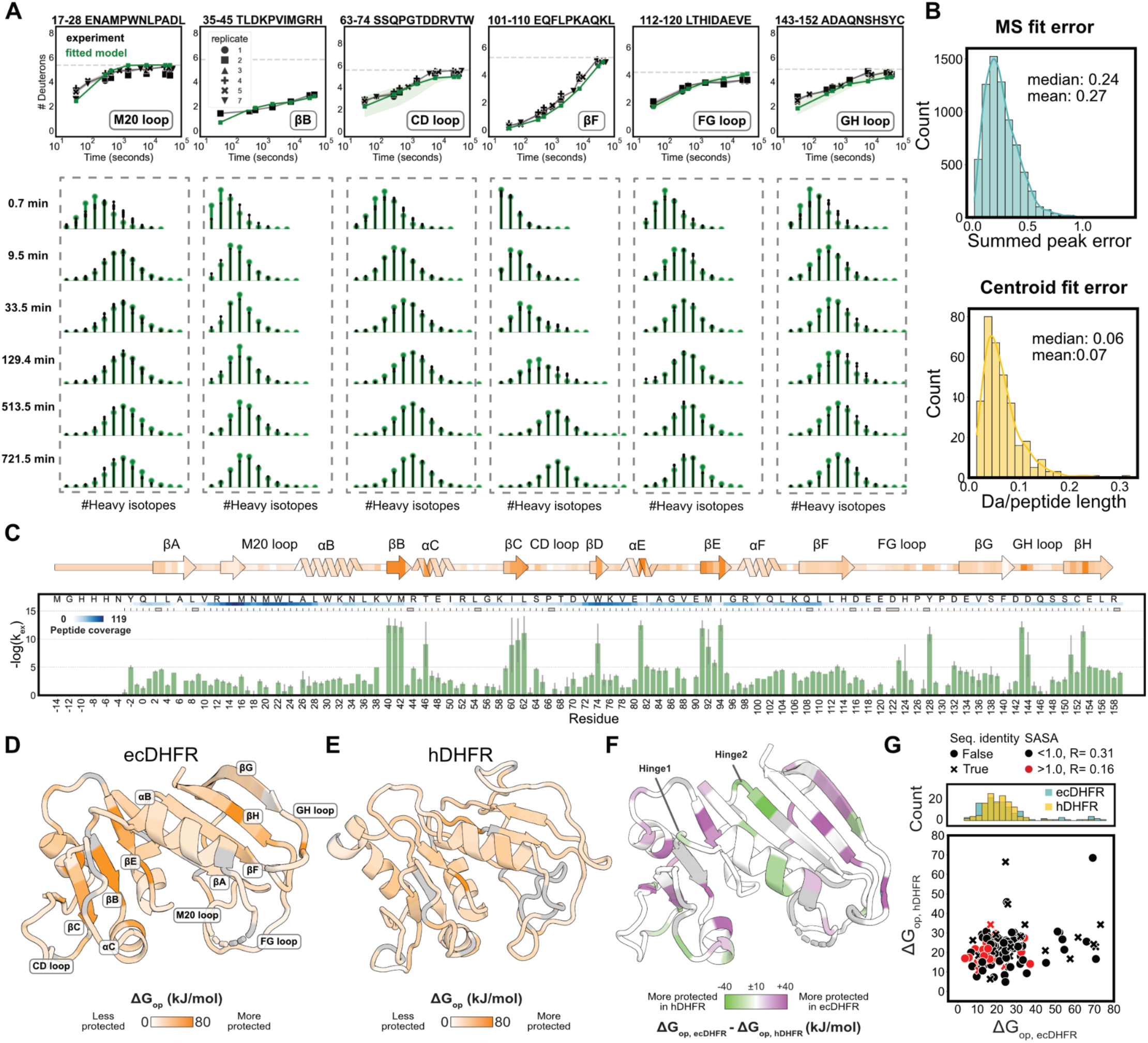
Site-resolved local energetics of *E. coli* and human DHFR orthologs. **A.** Representative fitted centroid uptake plots with corresponding isotopic mass envelope fitting. Dotted line represents experimental full D level. The fitted models are shown in green, and the shadows on the uptake plots are calculated based on the standard deviation of PFs within the cluster. Peptide positions on the protein are annotated in the lower right corner. **B.** Distribution of mass spectrum fit errors and centroid fit errors. **C.** FEATHER-derived exchange rates, formatted similarly to Fig. 2E. **D.** Absolute ΔG_op_ of ecDHFR projected onto the crystal structure in the APO state (PDB: 5DFR^76^) and the secondary structure of ecDHFR using SSDraw (C, top).^77^ Darker hues indicate amino acids with backbone amide groups that are more protected from HX. **E.** Absolute ΔG_op_ of hDHFR projected onto the crystal structure in the APO state (PDB: 4M6J). **F.** Difference between hDHFR and ecDHFR ΔG_op_ values projected onto the crystal structure of ecDHFR based on structural alignment with USalign.^78^ Proline and residues with insufficient or noisy data coverage are colored gray in panels D-F. **G.** Correlation of ΔG_op_ between ecDHFR and hDHFR. ecDHFR contains several comparatively highly protected residues (-log(k_ex_)>8 in panel C). The correlation is not higher among identical residues in the two orthologs.

From 341 PIGEON-disambiguated peptides, we resolved 86% of positions at single-site resolution and 6% as dipeptides, totaling 92% of the protein sequence (Figs. 3c, S3, Tables S7, S8). The coverage and resolution exceeds previous HX/MS experiments for ecDHFR^23–25^ due to using several acid proteases (Fig. S4). The projection of ΔG_op_ on the structure highlights βB, βC, and βE as the regions most protected from exchange (Fig. 3c, d). Critically, PIGEON-FEATHER can resolve single-site ΔG_op_ in ecDHFR’s catalytically important Met20, FG, and GH loops. The Met20 loop residues exchange quickly (Fig. 3a, c, d) as expected given diffuse electron density in APO ecDHFR X-ray crystal structures.^26,27^ However, the GH loop contains well-protected residues, like A143 and D144 (ΔG_op_=71, 45 kJ/mol).

The two PIGEON modes produce consistent FEATHER results with an R value of 0.90 (Fig. S5). Correlating ΔG_op_ from singlicate experiments against the pooled dataset (n=6) gives an average R value of 0.79, demonstrating the method’s reproducibility (Figs. S6, S7a-g). In the absence of long timepoints (>4 hours), we observed a drop of 0.05 in R, similar to the synthetic data benchmark (Figs. S7h, S8). We found that three bootstrapping replicates are sufficient (Fig. S9), and that FEATHER converges to consistent results even without the structural and uptake priors (Fig. S10). FEATHER-derived PFs show a 0.48 correlation with a widely used phenomenological model that defines the PF as a function of the numbers of amide hydrogen bonds and local contacts (Fig. S11).^28^ FEATHER-derived PFs also correlate with computationally calculated hydrogen bonds and solvent accessible surface area (SASA) using the macromolecular modeling suite Rosetta, suggesting a potential use of the method for structural modeling (Fig. S11).^29,30^

### Single-site *ΔG_op_* values for ecDHFR and hDHFR enable comparison

Like protein sequences and structures, ΔG_op_ values may be evolutionarily conserved as features of the conformational ensemble.^31^ Although the sequence identity between ecDHFR and hDHFR is only 26%^32^, the X-ray crystal structures are almost identical (all-atom RMSD=1 Å, PDB IDs: 4M6J, 5DFR).^33,34^ While peptide-level H-D exchange cannot be quantitatively compared for proteins with different sequences, absolute ΔG_op_ for structurally aligned regions of related proteins may be compared. We therefore applied HX/MS-PIGEON-FEATHER to determine ΔG_op_ values for hDHFR (Figs. 3e, S12, S13, Tables S9, S10). Despite their modest sequence identity, the residue-level ΔG_op_ values for the two DHFRs are moderately correlated (Fig. 4f, g).

**Figure 4.**
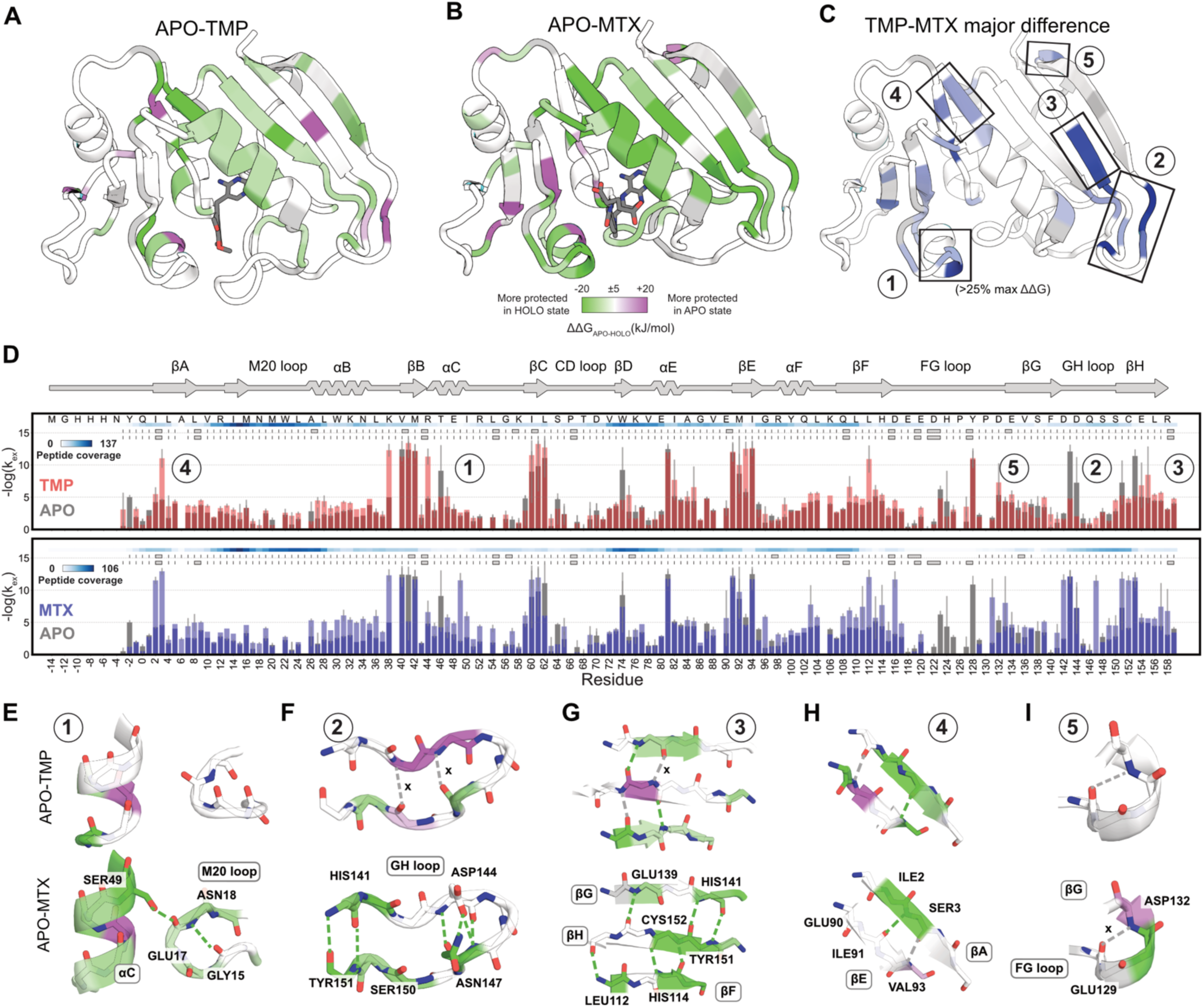
Two competitive inhibitors reweight the ecDHFR ensemble differently. **A.** ΔΔG_op (TMP-APO)_ mapped onto the crystal structure of TMP-ecDHFR (PDB: 6XG5^79^). **B.** ΔΔG_op (MTX-APO)_ mapped onto the crystal structure of MTX-ecDHFR (PDB: 1RG7^41^). **C.** Main difference between ΔG_op(MTX)_ and ΔG_op(TMP)_. **D.** Overlay of FEATHER-derived exchange rates for TMP and MTX with the APO state, formatted similarly to Fig. 3e. **E-I.** Close view of the Met20 loop, GH loop, central strands and small helix highlighted in C, showing molecular details of hydrogen bonds that differentiate the ΔΔG_op (TMP-APO)_ and ΔΔG_op (MTX-APO)._

Although the two orthologs are highly structurally similar, previous ^15^N relaxation dispersion experiments showed that the ms-scale motions that dominate ecDHFR Met20 loop are absent in hDHFR, and mutations in ecDHFR that disrupt these motions slow NADP^+^ dissociation.^35–37^ It was hypothesized that lower intracellular NADP^+^ and THF concentrations in vertebrates allow efficient cofactor exchange in hDHFR, enabling the maintenance of the structure of the Met20 loop, while promoting a hinge-like open/close conformational change in the central β sheet to control the ligand flux to the substrate binding pocket. Bacteria, which have ∼1:1 NADPH:NADP^+^, may instead have evolved a loop-mediated exchange mechanism to avoid end product inhibition.^38^

In support of this hypothesis, PIGEON-FEATHER yields the otherwise non-intuitive result that several positions in the central β sheet, a universally conserved DHFR motif, have much higher ΔG_op_ in ecDHFR than in hDHFR (Fig. 3g). Additionally, we find that several residues in the FG loop and the nucleotide binding domain are stabilized by 11-16 kJ/mol in ecDHFR vs. hDHFR (Fig. 3f, g, S13). In contrast, several peripheral residues in βF are stabilized by 21–42 kJ/mol in hDHFR compared to ecDHFR, illustrating the energetic cost associated with these global effects. We hypothesize that this stabilization compensates for the larger loop dynamics in ecDHFR and facilitates the hinge-like conformational change in hDHFR.^33^ In line with this hypothesis, the hinge loops, which mediate the open-close conformational change in hDHFR and are much shorter in ecDHFR, have higher ΔG_op_ in hDHFR than in ecDHFR, thereby acting as stabilizing pivot points for the coordinated motions of the two domains. PIGEON-FEATHER thus introduces a strategy to classify the ligand flux and catalysis mechanisms for many other DHFR orthologs with varied-length functional loops and hinges towards developing pathogen-specific antibiotics.^33,39^

### Two competitive inhibitors differentially impact ecDHFR and hDHFR ensembles

Trimethoprim (TMP) and methotrexate (MTX) are competitive inhibitors that bind ecDHFR with sub-nM affinities, but because TMP inhibits bacterial DHFRs at least four orders of magnitude more than vertebrate orthologs, it used as an antibiotic.^32,39,40^ X-ray crystal structures for TMP- and MTX-ecDHFR have <0.5 Å all-atom RMSD (Fig. 4a, b).^41,42^ NMR spectroscopy revealed near-identical µs-to-ms motions in the two inhibitor-bound states.^43^ Both states favor a conformation of the Met20 loop that blocks the active site.^41–44^ As mounting drug resistance mutations^45^ reduce inhibitor efficacy and motivate the search for new antibiotics, we revisited the question of how TMP and MTX impact the ecDHFR ensemble.

PIGEON-FEATHER-derived changes in ΔG_op_ (ΔΔG_op_) between the APO and inhibitor states revealed that TMP and MTX have distinct effects on ecDHFR (Fig. 4a-d, Figs. S13-19). Although both inhibitors stabilize the β sheet and binding pocket helices, they each also have unique energetic effects (Figs. 4, S13, S16-18), particularly at helix αC (Figs. 4e, S17), the Met20 loop (Figs. 4e, S17), the GH loop (Figs. 4f, S17), and the N-terminal region of αB (Fig. S17). Compared to the APO enzyme, αC is stabilized in MTX-ecDHFR via interactions with the Met20 loop, while TMP binding has no effect (Fig. 4e). In the αC-proximal loop, R57 forms bidentate hydrogen bonds with MTX that are not possible with TMP (Fig. S18). In αB, the difference is most pronounced in A26 (ΔΔG_op,MTX-TMP_=14 kJ/mol). Interestingly, allosteric TMP-specific changes in ΔG_op_ are pervasive in the β sheet where TMP resistance mutations are enriched, suggesting that these ensemble effects are integral to TMP binding in ecDHFR (Figs. 4g, h, S19). We highlight these changes in ΔG_op_ to demonstrate the resolution and quantification afforded by PIGEON-FEATHER compared to conventional HX/MS analysis methods, and to emphasize results that are not obvious from structures.

Altogether, PIGEON-FEATHER shows how each inhibitor reshapes the ecDHFR ensemble. MTX interacts with the M20 loop and introduces a backbone-sidechain hydrogen bond between E17 in the Met20 loop and S49 in αC (Fig. 4e), further stabilizing the backbone-backbone hydrogen bond between N18 and G15, while the GH, FG, and Met20 loops at the edge of the sheet form a strong intramolecular hydrogen bond network (Fig. 4b, f-i). The backbone hydrogen bonds in the central beta sheet (Fig. 4g, h) are stabilized in MTX-ecDHFR, while TMP binding triggers a stabilization/destabilization switch in this region. In the MTX-bound state, the backbone hydrogen bonds in βG are disrupted by the hydrogen bond rearrangement in the small helix that connects it with the FG loop (Fig. 4i). Because TMP binding does not bias the Met20 loop to the same conformation as MTX, the nucleotide-binding domain, FG loop, and βG strand remain relatively unperturbed (Figs. 4a, S13, S17-19).

These results combined with X-ray crystallography suggest that TMP’s selective inhibition of bacterial DHFRs may be linked to its binding conformation and to allosteric effects on the βG and βH strands. Generally, the amino acid content and local structure is similar in the active site among bacterial and vertebrate orthologs (Fig. S20).^39,46^ In ecDHFR and hDHFR complexes, the diaminopyrimidine ring of TMP is positioned similarly, but the trimethoxyphenyl group is positioned differently, with a 90° rotation in the dihedral angle for the pyrimidine-to-methylene bond (Fig. S21a).^42,47^ We hypothesize that this rotameric change may destabilize αB to reduce TMP-hDHFR affinity. Further, PIGEON-FEATHER shows that TMP binding also relies heavily on the rearrangement of hydrogen bonds in the β-sheet, especially the βG strand, which may have a higher energetic cost in vertebrate DHFR orthologs where the βG strand forms an extended loop with the adjoining FG loop (Figs. 3e, S21c). While previous studies demonstrated the unexpected impacts of allosteric mutations in the βG and βH strands on DHFR activity and TMP inhibition^48–53^, e.g. insertion of this extended loop from hDHFR into ecDHFR leads to its destabilization and deficiency in catalysis^48^, PIGEON-FEATHER offers molecular-level insights to contextualize these effects by reporting how ligand-induced conformational reweighting in this region contributes to ligand binding. TMP resistance mutations^54–56^ in the β sheet of ecDHFR may limit TMP efficacy by similarly disrupting the wild-type pattern of ligand stabilization in this region (Fig. S19). By contrast, we hypothesize that the energetic requirement for MTX binding in the β sheet is met by both ortholog ensembles and likely offset through MTX interactions with αC and three functional loops, resulting in the inhibition of both ecDHFR and hDHFR.

We confirmed this hypothesis via HX/MS-PIGEON-FEATHER to compare the effects of TMP and MTX on the hDHFR ensemble. As in ecDHFR, both inhibitors globally stabilize hDHFR (Figs. S22, S23, Tables S9, S10). However, the most significant differences in how TMP and MTX bind hDHFR are in the binding pocket due to MTX-specific interactions, a direct effect that is also observed in ecDHFR, and in the βG and βH strands, an indirect effect from ligand binding that is unique in hDHFR as we hypothesized (Fig. 5a-c). In summary, PIGEON-FEATHER revealed that TMP binding to the DHFR fold requires specific energetic contributions from the βG and βH strands (Fig. 5b), which is achieved by water-mediated interactions in the extended loop in βG in hDHFR (Fig. 5c, 3). In contrast, in ecDHFR, this stabilization is accomplished by direct backbone hydrogen bonds (Fig. 5d, 3). This supports a higher TMP binding affinity for ecDHFR.

**Figure 5.**
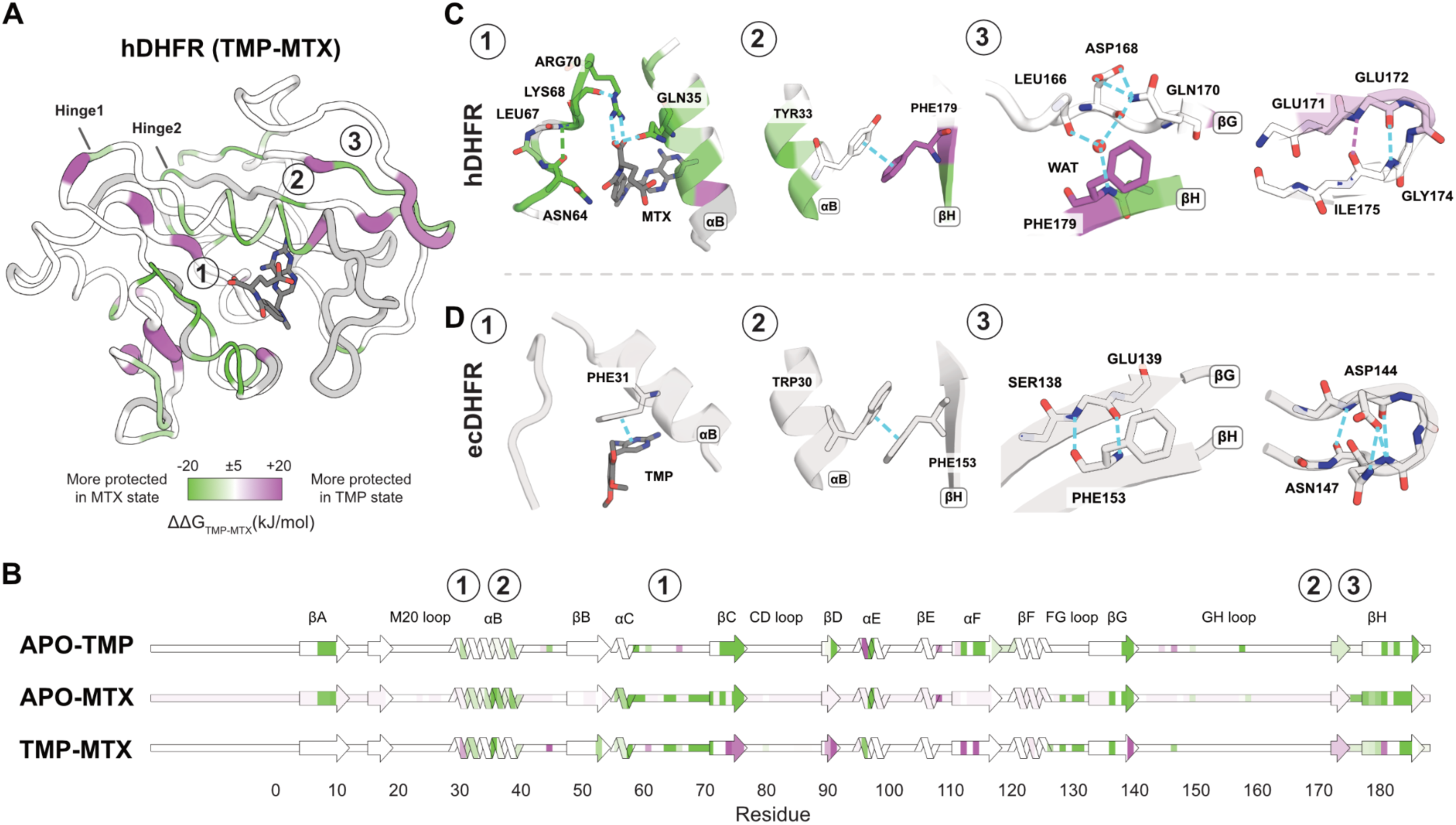
Comparing the effects of TMP and MTX binding on the hDHFR ensemble. **A.** ΔΔG_op (TMP-MTX)_ mapped onto the crystal structure of hDHFR (PDB: 1U72). **B.** Each binary comparison of FEATHER-derived ΔG_op_ between functional states with boxed regions from A labeled on the protein sequence. **C.** Close view of the boxed regions in A: 1) active site, 2) αB-βH region, and 3) βG-βH hydrogen bonds. **D.** For comparison to each panel in B, the same views of the crystal structure of ecDHFR (PDB: 6XG5).

### PIGEON-FEATHER reveals the transcriptional activation mechanism for the *lac* repressor (LacI)

A key advantage of coupling HX experiments to MS rather than NMR spectroscopy is that MS is higher-throughput and not limited by protein size. Thus, HX/MS-PIGEON-FEATHER is suitable for large and complex samples in various functional states, for example proteins that interact with both nucleic acids and small molecule regulators. We therefore applied this approach to tackle a classic problem in protein biophysics with unprecedented quantification, completeness, and resolution: how ligand binding to a large, dimeric transcription factor, LacI, regulates gene expression.

LacI is a model system from *E. coli*. Isopropyl β-D-1-thiogalactopyranoside (IPTG) binds the regulatory ligand-binding domain (LBD) of LacI to cause an order-disorder transition in the DNA-binding domain (DBD) more than 40 Å away, but X-ray crystal structures show <1.5 Å RMSD among the apo, DNA-bound, and IPTG-bound LBDs.^57^ We previously applied HX/MS to learn how IPTG and other ligands modulate the rigidity of secondary structures in the LBD.^58^ The results were well-correlated with functional data from mutational studies^59,60^ and were validated by subsequent NMR experiments^61^ that confirmed the roles of the same regions in stabilized and truncated mutants. However, these studies did not show how the energetic changes in the LBD upon IPTG binding actually remodel the DBD ensemble. We applied PIGEON-FEATHER with fully replicated HX/MS experiments to probe the allostery of this large protein at the residue level, including the DBD (Fig. 6). The peptide-level trends were consistent with our previous study (Figs. 6a, b, S24-26), but PIGEON-FEATHER finally revealed crucial biological insights about the IPTG to DNA ensemble transition.

**Figure 6.**
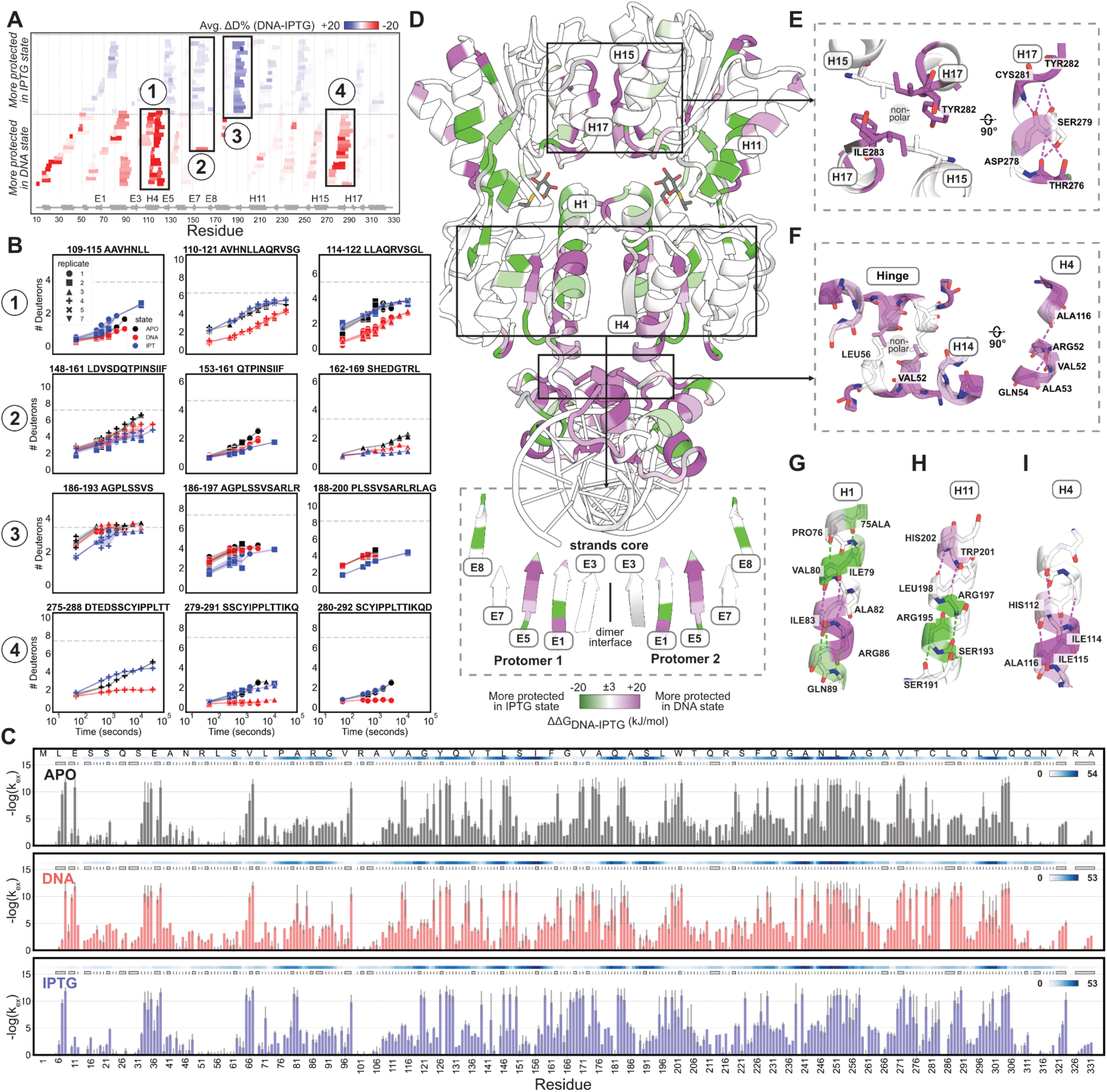
IPTG binding shifts the LacI ensemble to the transcriptionally active state. **A.** Woods plot showing the D uptake difference between the DNA and IPTG states. **B.** Representative uptake plots of peptides in the regions that differentiate the DNA and IPTG states, boxed in panel A. **C.** FEATHER-derived high-resolution exchange rates for LacI functional states. **D.** ΔΔG_op (IPTG-DNA)_ mapped onto the crystal structure of ecDHFR (PDB: 1EFA^80^), with IPTG from PDB: 2P9H^57^ aligned to show the binding pocket. **E-I.** Close-up views of the dimer interface at helix H15/H17, hinge helix and helix H4, helix H1, H11, and H4, showing molecular details of hydrogen bonds with ΔΔG_op (IPTG-DNA)._

We found that IPTG binding differentially reweights specific sets of interactions at the dimer interface across the N- and C-terminal lobes of the LBD over a 70 Å distance to disfavor LacI-DNA interactions and activate transcription (Fig. 6c, d). The C-terminal LBD subdomain is virtually identical by X-ray crystallography in the DNA- and IPTG-bound states.^57^ Here, PIGEON-FEATHER shows that helices H17 and H15 are more protected in the DNA state due to better hydrophobic interactions among I283, Y282, and L251, while further stabilizing the D278-C281 and D278-Y282 backbone hydrogen bonds (ΔΔG_op, DNA-IPTG_ =26, 25 kJ/mol) (Fig. 6e). Despite the 55 Å distance, this region is coupled to the DBD, where in the DNA state, a cross-interface hydrophobic interaction between L56 and V52 stabilizes the hinge helix and protects the R51-A116 hydrogen bond, further stabilizing helix H4 (Fig. 6f). This bond is a true molecular switch: in IPTG-LacI, the probability of this interaction is dramatically decreased to bias the ensemble to its active state.

Helix H1 in the N-terminal subdomain of the LBD is one of the most elusive helices in LacI. We previously observed polymorphism in its exchange behavior: the N-terminal half is more protected from H-D exchange in the IPTG state, while the C-terminal half is less protected (Fig. S25). Using PIGEON-FEATHER, we determined that the N-terminal protection at A75 and I79 originates from IPTG binding (alongside S193 and S197 in the pocket periphery), while the C-terminal deprotection is due to the weakening of the hydrogen bonds R86-A82, S85-A81, K84-V80, I83-I79 (ΔΔG_op, DNA-IPTG_ =17, 17, 22, 9 kJ/mol), caused by the LBD hinge motion (Fig. 6g, h). The IPTG-triggered disruption to this region culminates in destabilization of the C-terminal region of helix H4 and the hinge helix (Fig. 6f, i). The interactions of the hinge helices with each other and with the minor groove of the DNA operator are therefore reduced^62^, which linearizes the DNA and releases the helix-turn-helix motif of the DBD from the major groove via increased local unfolding (Fig. 6d).

## Discussion

PIGEON-FEATHER determines high-resolution ΔG_op_ values as energetic features of protein ensembles from conventional HX/MS measurements. Detailed benchmarks show that it surpasses other methods and scales with protein size for nearly perfect accuracy, improving the utility and interpretability of HX/MS such that it may be seamlessly combined with cryo-EM and MD simulations. PIGEON-FEATHER includes several features that contribute to its high accuracy. After disambiguating peptides using tandem MS fragment identification, it fits their full isotopic mass envelopes, which circumvents the degeneracy of centroid fitting inherent in other methods by introducing additional constraints and minimizing calculation uncertainties. Inspired by the physics of the hydrogen exchange phenomenon, PIGEON-FEATHER’s sampling mechanism dramatically improves sampling efficiency on the rugged ΔG_op_ space in the MCMC simulations. The peptide subtraction feature further improves the method’s accuracy.

We used PIGEON-FEATHER to learn three secrets of protein behavior. First, we determined how slow-timescale, functionally important conformational ensemble equilibria are conserved and diverged in DHFR orthologs to learn how two organisms adapted the same ancient protein fold to perform and regulate catalysis in different cellular environments. Then we discovered the molecular basis for bacterial DHFR inhibition by the antibiotic TMP and contrasted its binding mechanism with the broad-spectrum chemotherapeutic agent MTX to demonstrate how two competitive inhibitors bind DHFR orthologs in different ways, leading to different species selectivity and informing future drug design efforts. Finally, we extended PIGEON-FEATHER to the large LacI-DNA complex to reveal how IPTG binding reshapes the LacI ensemble to activate transcription.

The power of PIGEON-FEATHER lies in its ability to reveal how proteins traverse free energy landscapes in response to various perturbations, including ligand binding, mutations, and environmental conditions (pH, temperature, pressure, solvent, and denaturant). The inclusion of ensemble energetic information adds an extra dimension to the sequence-structure-function paradigm for proteins. Integrated with recent advances in protein structure determination^63^, structure prediction^7,64^, structure search^65^, and bioinformatic analysis^66,67^, PIGEON-FEATHER enables a deeper understanding of proteins and protein families. Because the method permits the comparison of ΔG_op_ within fold families, biologists may directly observe changes in ensemble energies over the course of evolution. PIGEON-FEATHER reveals allosteric protein mechanisms at high resolution, which can aid the identification and design of new therapeutics.

PIGEON-FEATHER can also be integrated into protein engineering and computational structural modeling methods, for example to build residue-level scoring functions for enhanced docking and virtual epitope screening, integrating with structural modeling^68^ and prediction^6,69^. With PIGEON-FEATHER, HX/MS datasets can be used to train machine learning models aimed at designing proteins with defined free energy landscapes. In principle, PIGEON-FEATHER can be extended to support multimodal distributions and HX-electron capture dissociation (ECD)/electron transfer dissociation (ETD)/MS. We will incorporate these capabilities in future developments.

## Methods

### PIGEON: detailed methods

#### Initial peptide identification and removal of systematic bias

PIGEON requires a set of MS2 datasets, as well as a sequence for each protein of interest. PIGEON can disambiguate multiple proteins in complexes or batched samples. We recommend including all proteases used in the analysis to rule out matches to self-cleavage peptides. Compatible compound lists in .mgf format can be produced from common MS data formats by exporting directly from commercially available MS software or with the open source MSConvert software.^70^ We collect 2-4 MS2 datasets per HX/MS experiment and produce a compound list in .mgf format with Bruker Compass DataAnalysis. We match MS1 and MS2 peaks to theoretical peptide and fragment *m/z* values^70,71^ and assign a score to each match (MS compound + theoretical peptide):

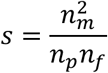

Where n_m_ is the number of fragment ion matches, n_p_ is the number of MS2 peaks in the compound, and n_f_ is the number of theoretical fragment ions for the peptide. To account for different protease specificities, heterogeneity in protease activities, and in-source fragmentation, our database includes every possible proteolytic fragment for each protein in the system up to a maximum length of 15 residues (Extended Data Fig. 1b). To account for any possible systematic *m/z* errors from the MS calibration, we apply an initial high 30 ppm tolerance for *m*/*z* error (Extended Data Fig.1e-f, Figure S1a). The 30 ppm tolerance introduces many false identifications, so we impose a lower tolerance threshold after correcting for calibration errors using PIGEON (Extended Data Fig.1e, ii). To do this, PIGEON first pools all identified peaks across MS2 datasets and identifies a subset of high-confidence peaks (with scores >0.05) from this pool (Fig. S1b), then fits a heuristic trend line to the subset using a fractional exponent function or polynomial function. We re-threshold the data to ±7 ppm to include only points around this trendline (Fig. S1c). Up to half of the peaks are discarded at this stage, with up to 100% increase in mean score.

#### Choice of consensus match for each peptide

In HX/MS experiments, one to three charge states are typical per peptide. Each charge state for each peptide is typically matched to multiple MS compounds. These correspond to the same MS1 peak, but differ by small amounts in retention time, *m*/*z*, and MS2 spectra. In cases of mass degeneracy, peptides may also be matched to additional compounds corresponding to other MS1 peaks with different retention times. PIGEON chooses the match with the highest fragment score for each peptide. In the approximately 30% of cases with equal fragment score among matches, PIGEON chooses the match with higher MS1 peak intensity. This leaves 100-1,000 peaks, with a 2-fold to 4-fold increase in mean score (Extended Data Fig. 1e, ii, Fig. S1d). At this stage, the user may also impose a minimum score for matches, for use with higher quality datasets.

#### Removal of false duplicates

The most problematic case for HX/MS analysis is *m*/*z* degeneracy, where multiple peptides are redundantly assigned to a single peak in the MS because their theoretical *m*/*z* or masses are very close or identical (Extended Data Fig. 1a-c). For each consensus peak assignment, PIGEON searches all other peak assignments in the dataset for mass duplicates. PIGEON collects all matches for which the monoisotopic *m*/*z* and the retention time are within a threshold (default: 0.1 Da, 0.5 minutes) of the consensus match in question, and makes two lists. List (1) contains all matches within the threshold corresponding to the same peptide, and list (2) contains all matches within the threshold corresponding to other peptides (these matches are degenerate). If list (2) is empty, the peptide is retained. Otherwise, PIGEON performs fragment-based disambiguation. For each list, all fragment ion matches that also correspond to a fragment ion match in the other list are removed. If list (2) is not empty and there are no fragment matches remaining in list (1), the peptide is removed (Extended Data Fig. 1e, iii, g). Otherwise, PIGEON calculates a p-value for every match in each list based on the remaining (unique) fragment ion matches, according to a binomial test:

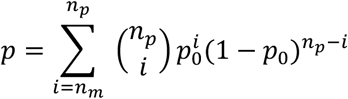

Where n_p_ is the number of peaks in the MS2 scan, n_m_ is the number of observed fragment matches, and p_0_ is the probability of each peak being matched to some fragment of the matched peptide under the null hypothesis that MS2 peaks are randomly distributed, given by:

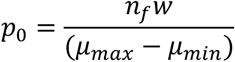

Where n_f_ is the number of theoretical fragment ions for the matched peptide, w is the width in *m/z* of the tolerance interval for fragment matches, and μ_max_ and μ_min_ are the maximum and minimum *m/z* values across all MS2 peaks and theoretical fragments.

This p is the likelihood of obtaining at least the observed number of fragment matches given the number of possible fragment ions and the number of MS2 peaks, assuming MS2 peaks are random noise not corresponding to the matched peptide. PIGEON then uses Fisher’s method for combining p-values to calculate joint p-values for sets of peptides.

The joint p is the likelihood of obtaining at least as many fragment matches as observed, treating every matched scan as an independent experiment, under the null hypothesis that none of the matched peptides are present and therefore p-values for individual experiments are uniformly distributed. Joint p-values p_1_ and p_2_ are calculated for lists (1) and (2) of matches. If p_1_ is not significant (default: p>0.05), the peptide in question is discarded. If p_2_ is significant, this indicates that other peptides co-elute and are present in an unknown ratio.

Because the optimal treatment in this case depends on the details of the analysis (*e.g.*, data coverage, resolution, and manual refinement) the user may either retain or remove the peptide of concern using the ‘keep’ or ‘drop’ PIGEON modes, respectively (Extended Data Fig. 1h).

### FEATHER: detailed methods

The FEATHER method uses a Bayesian framework to reconstruct full isotopic mass envelopes to fit PFs given three inputs from the user: the protein sequence, a peptide list from PIGEON, and the HX/MS dataset.

#### Peptide subtraction

The experimental peptides from the HX/MS dataset for each functional state of the protein are categorized according to their overlaps. Subsequently, exhaustive combinations of peptides within each category are subjected to subtraction (Fig. 2b). Peptides are subtracted iteratively until no further new peptides can be produced. There is also an option to only perform one round of subtraction. We use a Monte Carlo (MC) simulation to subtract the isotopic mass envelope, aiming to minimize the error recalculated between the convolution of the two short peptides and the parent peptide. The initial guess for this process is obtained through the deconvolution of the parent peptides using a Fourier transform.

#### Isotopic mass envelope reconstruction

Following peptide subtraction, initial ΔG_op_ guesses for each individual amino acid are updated using Markov Chain Monte Carlo (MCMC) sampling, by comparing the theoretical and measured mass envelopes along with the centroid value to estimate the posterior distributions of ΔG_op_ until convergence is achieved (Fig. 2a). The theoretical envelopes are derived by convolving the natural isotopic mass envelope from the MS2 experiment with the theoretical deuteration isotopic distribution at a specific time point. The deuteration isotopic distribution is calculated based on the single exponential kinetic curves of all residues in a peptide given the ΔG_op_ and the peptide’s back exchange level, determined from a fully deuterated control experiment. If the control experiment was not performed, FEATHER includes an option to use the theoretical value for the maximum peptide back exchange. The peptide exchange does not take into account the first two residues and prolines. The forward model (f_mod_), which calculates the isotopic mass envelope E_f,t_^fmod^ of peptide *f* at exchange time *t*, is defined as:

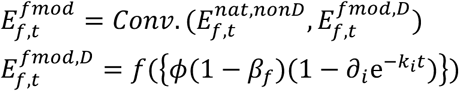

where Conv. is the convolution function, f(x) is the discrete probability density function of the binomial distribution, calculating the probability of obtaining 0 to n deuterated residues for a peptide of length n, saturation level *ϕ* is the deuterium percent of the D_2_O buffer, ∂_i_ is a delta function if residue *i* has an observable amide and zero otherwise, β_f_ is the back-exchange level of peptide f, and E_f,t_^nat,non-D^ is the natural isotopic mass envelope.

Working from the BayesianHDX^17^ score function, we replaced the original noise model with a Gaussian function of the summed squared error for individual isotopic mass envelope peaks, E_f, t, n, p_^exp^, for peptide *f* at time *t*, replicate *n,* and peak *p*. P_E_ represents the probability of observing the envelope E.

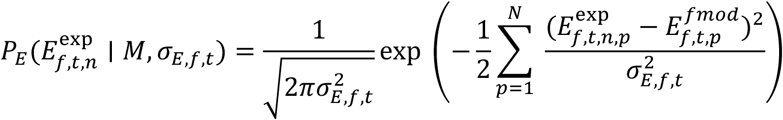

Here, the model M = ({k_i_}), is of the set of residue-resolved exchange rates, {k_i_}. N is the total number of peptides, σ_E, f,t_ is the average mass envelope peak error estimate for peptide *f* at time *t*.

We also incorporate the agreement of the fits with the centroid-level observations. The probability P_C_ of experimentally observed centroid value of the mass spectrum C^exp^ for peptide *f* at time *t*, replicate *n,* is given by:

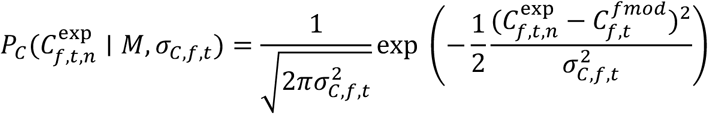

σ_C,f,t_ is the average centroid value error estimate for peptide *f* at time *t*.

The joint likelihood of isotopic mass envelope and centroid value of the mass spectrum is:

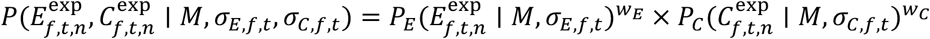

Where w_C_ and w_E_ are the weights of the centroid and envelope contributions, respectively. Their default values are both set to 1.

The likelihood function is the joint likelihood of all the peptides, timepoints and replicates.

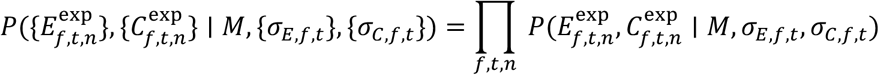

The ΔG_op_ are derived by combining results from multiple posterior analyses, each of which employs bootstrapping techniques where 90% of the peptides are randomly resampled from the entire dataset. In a typical bootstrapping replicate fitting process, simulated annealing is applied with the temperature decreasing from 50 to 0.1. At temperatures of 2 and 0.01, 1500 models are saved. This process is repeated five times, resulting in the collection of 7500 models. Finally, the top models (n=200 by default) from each bootstrap replicate are collected and pooled. K-means clustering is then applied to obtain the ΔG_op_ within each resolution segment, where the number of clusters corresponds to the size of the resolution segment (1 for a single resolved residue). The standard deviation of ΔG_op_ within the cluster, when combined across multiple bootstrapping runs, is used as an uncertainty measure.

#### Log(PF) swapping

During each sampling step, 20% of residues are randomly selected. Each of the residues in this subset may swap its log(PF) value with that of an adjacent residue (Fig. 2c). The probability that this swap is accepted is contingent upon the Metropolis criterion based on the score function at a temperature (T) of 0.

#### Priors

We also implemented two optional Bayesian priors to enhance FEATHER performance. The *structural prior* leverages computationally predicted or experimentally solved protein structures to bias the sampling based on probable hydrogen bonding patterns within the protein ensemble. This prior is justified by seminal work that demonstrated how backbone amide groups involved in hydrogen bonds with other protein atoms, ligands, or ordered water molecules limit H-D exchange.^10,72^ The *uptake prior* compares the FEATHER-determined exchange rates for each amino acid with the D uptake plots to ensure consistency with empirical observations in the calculated exchange rates. During the sampling process, two broad Gaussian biases are applied to the score function for positions with low and high PFs, based on their categorization according to D uptake.

In the structural and uptake priors, observable residues are categorized into three groups: high PF, low PF, and regular PF residues. This classification is based on an analysis of backbone hydrogen bonds in the protein structure and peptide uptake plots. For the structural prior, a residue is considered a high PF residue if it forms hydrogen bonds with other protein atoms, binding partners, or water molecules. Residues that lack backbone hydrogen bonds are classified as low PF residues. For the uptake prior, residues covered by slow-exchange peptides, where the maximum deuterium uptake is less than 20%, are considered high PF residues. Conversely, residues covered by fast-exchange peptides, where the minimum deuterium uptake is more than 100% (exceeding 100% due to full D sample error), are classified as low PF residues. Residues covered by both slow and fast exchange peptides are categorized as regular PF residues. To account for these categorizations in the sampling process, two soft biases are applied to the score function. For high PF residues, a prior Gaussian norm(6, 4.0) is added, and for low PF residues, a bias norm(1.5, 2.0) is included based on seminal NMR studies.^10,72^ The scales of the structural and uptake priors are set to 0.7 and 0.5, respectively. The PFs of regular residues are sampled evenly.

### Simulated data creation

An amino acid sequence and a corresponding array of ΔG_op_ of identical length and within a specified range (*e.g.*, 2–10) were randomly generated using the python built-in *random* module. The intrinsic exchange rates (k_int_) were calculated using the HDXrate library^73^ given the protein sequence and the pH and temperature of the experiment (pH 7, 293.0 K)^11,74^. Subsequently, the deuterium uptake kinetics of each residue *i* were determined using a single exponential function that incorporates PF and k_int_.

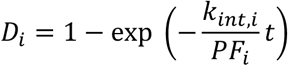

Peptides of lengths ranging from 5 to 12 residues were then randomly and evenly generated along the sequence range. The theoretical isotopic mass of a peptide was obtained using the forward model described above. We employed pyOpenMS^75^ to calculate the natural isotopic mass envelope of a simulated peptide. The peptide deuterium incorporation is a sum of deuterium uptake of all the residues within it except for the first two fast exchanging ones. The simulated data generation is optimized to achieve coverage comparable to that of the experimental dataset by ensuring a similar number of subtracted peptides after one round of subtraction.

## Data availability

All data needed to reproduce this work are available at the PRIDE database^81^ (dataset identifier PXD057539).

## Code availability

The source code for the software is available at https://github.com/glasgowlab/PIGEON-FEATHER, along with the synthetic datasets used for benchmarking, an example HX/MS dataset, and a tutorial.

## Acknowledgements

We gratefully acknowledge Glasgow Lab members for requesting, adding, discussing, testing, and troubleshooting various features of the software. PIGEON-FEATHER also benefited from testing by Dr. Danielle Swingle at the Advanced Science Research Center (ASRC) at the City University of New York (CUNY). We thank Dr. Andrea Piserchio, Dr. Kevin Gardner, and Dr. Rinat Abzalimov from the ASRC, as well as Dr. Shawn Costello, Sophie Shoemaker, and Dr. Naomi Latorraca for feedback on the method. We also thank Dr. Abzalimov in his role as the Mass Spectrometry Facility Manager at the ASRC, where the HX/MS experiments were performed. We are grateful to Dr. Supriya Pratihar, Dr. Hashim Al-Hashimi, and Dr. Art Palmer at Columbia University Medical Center for valuable discussions, and to Dr. Costello, Dr. Al-Hashimi, Dr. Abzalimov, Dr. Roksana Azad, Dr. Jeff Glasgow, Dr. Tanja Kortemme, and Dr. Stavros Lomvardas for critical feedback on this manuscript.

---

**Extended Data Fig 1.**
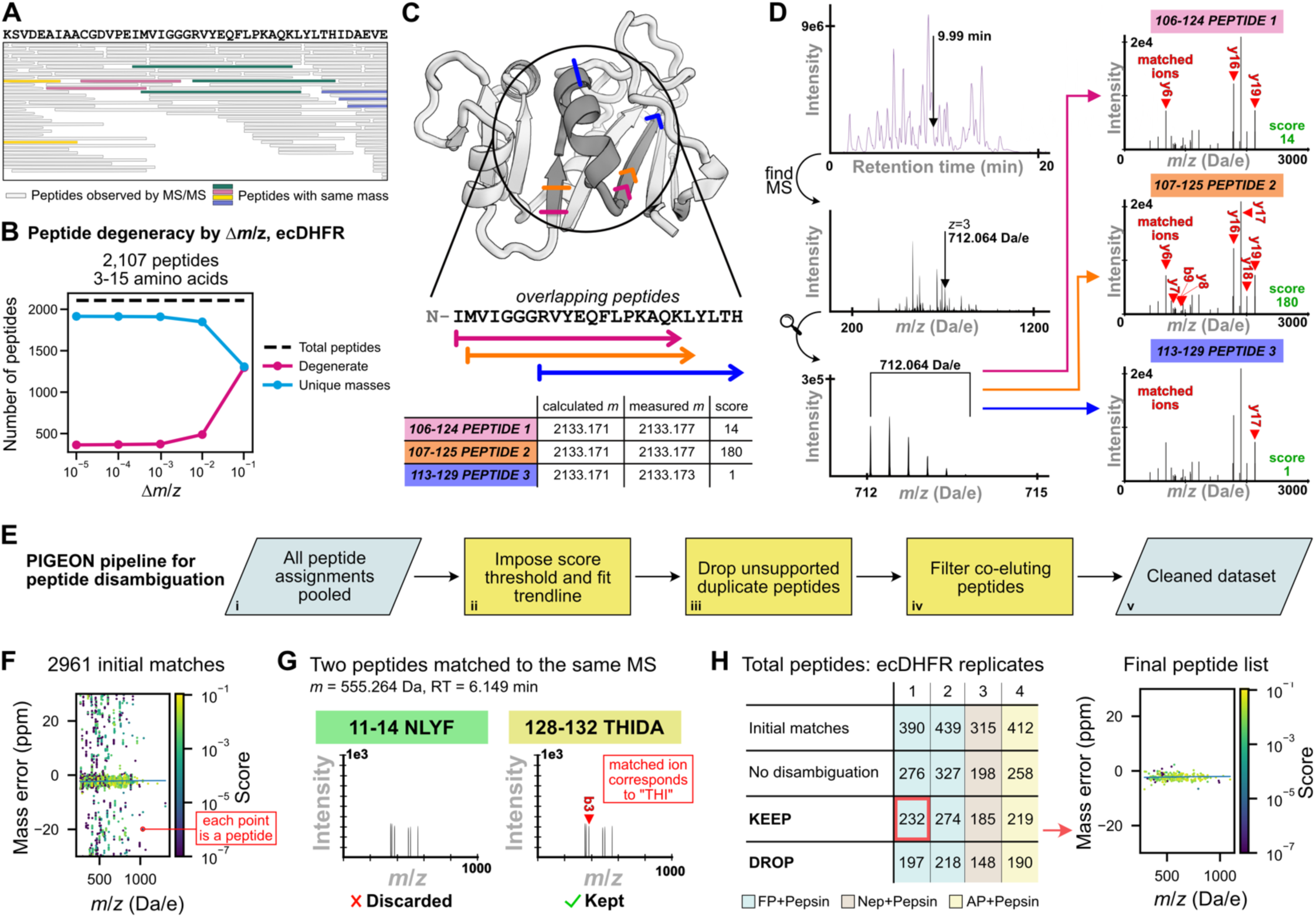
Peptide disambiguation with PIGEON. **A.** HX/MS peptides are frequently degenerate in mass. **B.** The number of degenerate peptides in *E. coli* DHFR (ecDHFR) given measurement uncertainty (Δ*m*/*z* ∼0.001 within experiments, ∼0.01 between experiments). **C.** ecDHFR peptides are duplicatively matched to the same peak. **D.** Disambiguation of peptides in panel C using tandem MS fragment information. The natural isotopic distribution corresponding to a single MS compound is shown, as extracted from the marked portion of the base peak intensity chromatogram. MS2 ions from this compound are compared to theoretical ions for the matched peptides, and the matches scored based on the degree of fragment ion support. **E. i.** MS2 data are pooled for all replicates and matched to theoretical peptides for each protein in the dataset. **ii.** Systematic mass errors are corrected by fitting a trendline to the highest scoring matches and re-thresholding around the trendline. All but the best peak for each peptide are discarded. **iii.** Each peptide for which there exists an identical match and which is not adequately supported by MS fragments is dropped. **iv.** Each charge state for which there exists an identical match (as in iii.) but which has fragment support is either kept (‘KEEP’), or discarded (‘DROP’). These peptides co-elute and both contribute to the signal. **v.** The final cleaned dataset contains peptides with scores distributed over the full range. **F.** Initial score and *m*/*z* error distribution for a representative pooled dataset (as in E, i). **G.** A duplicate match where the fragment ions support one peptide but not the other. **H.** Impact of each PIGEON step (systematic error correction and disambiguation) and peptide selection mode for four ecDHFR HX/MS datasets (Tables S1, S7). The use of multiple proteases increases peptide coverage: fungal protease (FP), pepsin, nepenthesin (NP), and alanyl aminopeptidase (AP). The *m*/*z* and score distribution for one resulting peptide pool (red box) are shown.

**Extended Data Fig 2.**
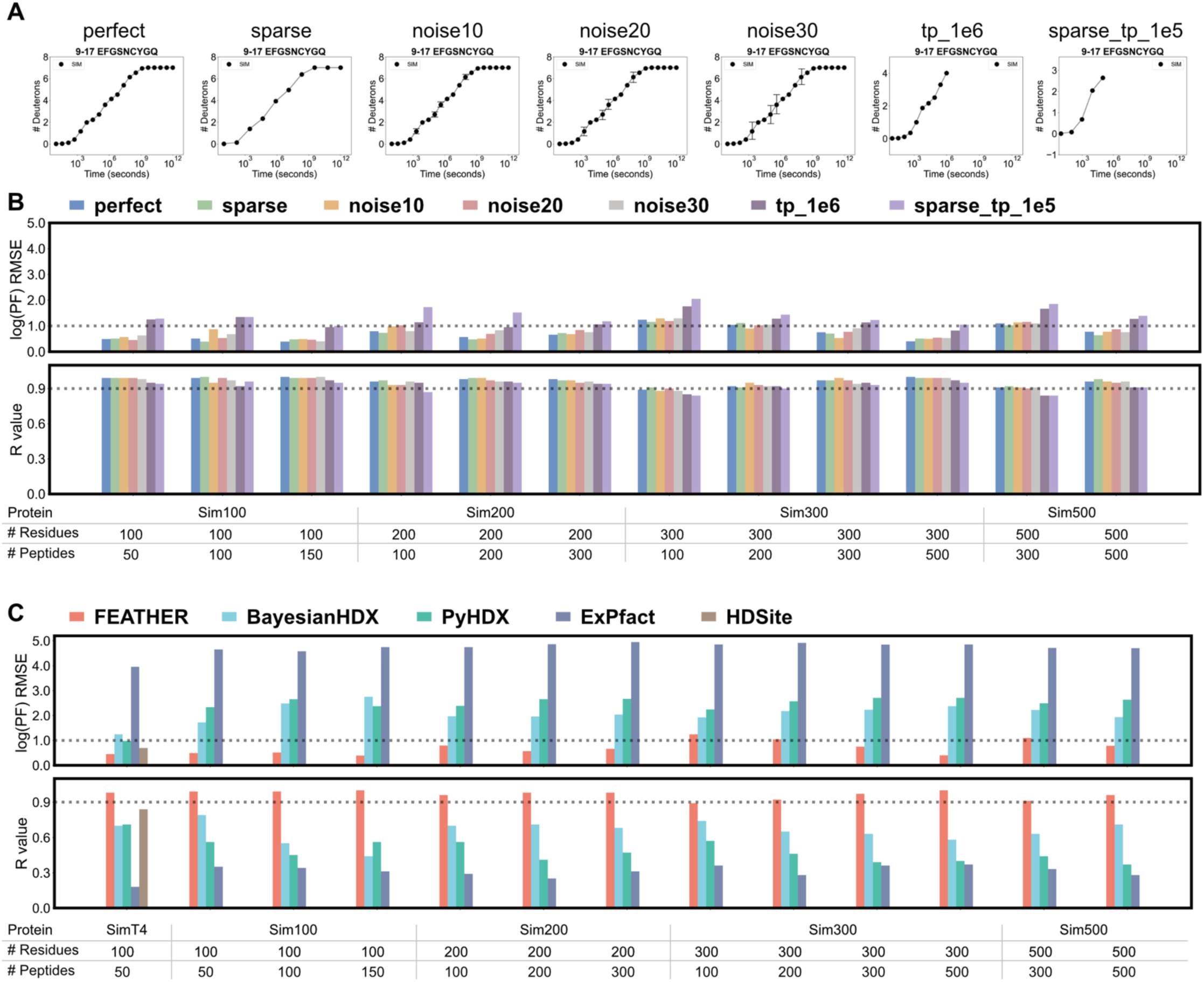
FEATHER benchmarks. **A.** Simulated datasets with varying data quality. **B.** Benchmarks on the simulated datasets. **C.** Comparison to other methods using the perfect simulated dataset. The raw data are available in Tables S5 and S6.

## References

1. Boehr, D. D., Nussinov, R. & Wright, P. E. The role of dynamic conformational ensembles in biomolecular recognition. Nat. Chem. Biol. 5, 789–796 (2009).

2. Cooper, A. Thermodynamic fluctuations in protein molecules. Proc. Natl. Acad. Sci. 73, 2740–2741 (1976).

3. Hilser, V. J. An Ensemble View of Allostery. Science 327, 653–654 (2010).

4. Kish, M. et al. Online Fully Automated System for Hydrogen/Deuterium-Exchange Mass Spectrometry with Millisecond Time Resolution. Anal. Chem. 95, 5000–5008 (2023).

5. Espada, A. et al. A Decoupled Automation Platform for Hydrogen/Deuterium Exchange Mass Spectrometry Experiments. J. Am. Soc. Mass Spectrom. 30, 2580–2583 (2019).

6. Krishna, R. et al. Generalized biomolecular modeling and design with RoseTTAFold All-Atom. Science 384, eadl2528 (2024).

7. Abramson, J. et al. Accurate structure prediction of biomolecular interactions with AlphaFold 3. Nature 630, 493–500 (2024).

8. Konermann, L., Pan, J. & Liu, Y.-H. Hydrogen exchange mass spectrometry for studying protein structure and dynamics. Chem. Soc. Rev. 40, 1224–1234 (2011).

9. Bai, Y., Sosnick, T. R., Mayne, L. & Englander, S. W. Protein Folding Intermediates: Native-State Hydrogen Exchange. Science 269, 192–197 (1995).

10. Skinner, J. J., Lim, W. K., Bédard, S., Black, B. E. & Englander, S. W. Protein hydrogen exchange: Testing current models. Protein Sci. 21, 987–995 (2012).

11. Bai, Y., Milne, J. S., Mayne, L. & Englander, S. W. Primary structure effects on peptide group hydrogen exchange. Proteins Struct. Funct. Bioinforma. 17, 75–86 (1993).

12. Masson, G. R. et al. Recommendations for performing, interpreting and reporting hydrogen deuterium exchange mass spectrometry (HDX-MS) experiments. Nat. Methods 16, 595–602 (2019).

13. Weis, D. D. Recommendations for the Propagation of Uncertainty in Hydrogen Exchange-Mass Spectrometric Measurements. J. Am. Soc. Mass Spectrom. 32, 1610–1617 (2021).

14. Eggertson, M. J., Fadgen, K., Engen, J. R. & Wales, T. E. Considerations in the Analysis of Hydrogen Exchange Mass Spectrometry Data. in Mass Spectrometry Data Analysis in Proteomics (ed. Matthiesen, R.) 407–435 (Springer, New York, NY, 2020). doi:10.1007/978-1-4939-9744-2_18.

15. Kan, Z., Ye, X., Skinner, J. J., Mayne, L. & Englander, S. W. ExMS2: An Integrated Solution for Hydrogen– Deuterium Exchange Mass Spectrometry Data Analysis. Anal. Chem. 91, 7474–7481 (2019).

16. Guttman, M., Weis, D. D., Engen, J. R. & Lee, K. K. Analysis of Overlapped and Noisy Hydrogen/Deuterium Exchange Mass Spectra. J. Am. Soc. Mass Spectrom. 24, 1906–1912 (2013).

17. Saltzberg, D. J. et al. A Residue-Resolved Bayesian Approach to Quantitative Interpretation of Hydrogen– Deuterium Exchange from Mass Spectrometry: Application to Characterizing Protein–Ligand Interactions. J. Phys. Chem. B 121, 3493–3501 (2017).

18. Smit, J. H. et al. Probing Universal Protein Dynamics Using Hydrogen–Deuterium Exchange Mass Spectrometry-Derived Residue-Level Gibbs Free Energy. Anal. Chem. 93, 12840–12847 (2021).

19. Stofella, M., Skinner, S. P., Sobott, F., Houwing-Duistermaat, J. & Paci, E. High-Resolution Hydrogen– Deuterium Protection Factors from Sparse Mass Spectrometry Data Validated by Nuclear Magnetic Resonance Measurements. J. Am. Soc. Mass Spectrom. 33, 813–822 (2022).

20. Kan, Z.-Y., Walters, B. T., Mayne, L. & Englander, S. W. Protein hydrogen exchange at residue resolution by proteolytic fragmentation mass spectrometry analysis. Proc. Natl. Acad. Sci. 110, 16438–16443 (2013).

21. Zhang, Z., Zhang, A. & Xiao, G. Improved Protein Hydrogen/Deuterium Exchange Mass Spectrometry Platform with Fully Automated Data Processing. Anal. Chem. 84, 4942–4949 (2012).

22. Huennekens, F. M. The methotrexate story: A paradigm for development of cancer chemotherapeutic agents. Adv. Enzyme Regul. 34, 397–419 (1994).

23. Oyeyemi, O. A. et al. Comparative Hydrogen–Deuterium Exchange for a Mesophilic vs Thermophilic Dihydrofolate Reductase at 25 °C: Identification of a Single Active Site Region with Enhanced Flexibility in the Mesophilic Protein. Biochemistry 50, 8251–8260 (2011).

24. Yamamoto, T. Mass Spectrometry on Segment-Specific Hydrogen Exchange of Dihydrofolate Reductase. J. Biochem. (Tokyo) 135, 17–24 (2004).

25. Wales, T. E. et al. Resolving chaperone-assisted protein folding on the ribosome at the peptide level. Nat. Struct. Mol. Biol. 1–10 (2024) doi:10.1038/s41594-024-01355-x.

26. Schnell, J. R., Dyson, H. J. & Wright, P. E. Structure, Dynamics, and Catalytic Function of Dihydrofolate Reductase. Annu. Rev. Biophys. 33, 119–140 (2004).

27. Wan, Q. et al. Toward resolving the catalytic mechanism of dihydrofolate reductase using neutron and ultrahigh-resolution X-ray crystallography. Proc. Natl. Acad. Sci. 111, 18225–18230 (2014).

28. Vendruscolo, M., Paci, E., Dobson, C. M. & Karplus, M. Rare Fluctuations of Native Proteins Sampled by Equilibrium Hydrogen Exchange. J. Am. Chem. Soc. 125, 15686–15687 (2003).

29. Alford, R. F. et al. The Rosetta All-Atom Energy Function for Macromolecular Modeling and Design. J. Chem. Theory Comput. 13, 3031–3048 (2017).

30. Nguyen, T. T., Marzolf, D. R., Seffernick, J. T., Heinze, S. & Lindert, S. Protein structure prediction using residue-resolved protection factors from hydrogen-deuterium exchange NMR. Structure 30, 313–320.e3 (2022).

31. Hollien, J. & Marqusee, S. Structural distribution of stability in a thermophilic enzyme. Proc. Natl. Acad. Sci. 96, 13674–13678 (1999).

32. Liu, C. T. et al. Functional significance of evolving protein sequence in dihydrofolate reductase from bacteria to humans. Proc. Natl. Acad. Sci. 110, 10159–10164 (2013).

33. Bhabha, G. et al. Divergent evolution of protein conformational dynamics in dihydrofolate reductase. Nat. Struct. Mol. Biol. 20, 1243–1249 (2013).

34. Bystroff, C. & Kraut, J. Crystal structure of unliganded Escherichia coli dihydrofolate reductase. Ligand-induced conformational changes and cooperativity in binding. Biochemistry 30, 2227–2239 (1991).

35. Bhabha, G. et al. A Dynamic Knockout Reveals That Conformational Fluctuations Influence the Chemical Step of Enzyme Catalysis. Science 332, 234–238 (2011).

36. Cameron, C. E. & Benkovic, S. J. Evidence for a functional role of the dynamics of glycine-121 of Escherichia coli dihydrofolate reductase obtained from kinetic analysis of a site-directed mutant. Biochemistry 36, 15792–15800 (1997).

37. Miller, G. P. & Benkovic, S. J. Strength of an Interloop Hydrogen Bond Determines the Kinetic Pathway in Catalysis by Escherichia coli Dihydrofolate Reductase. Biochemistry 37, 6336–6342 (1998).

38. Appleman, J. R. et al. Atypical transient state kinetics of recombinant human dihydrofolate reductase produced by hysteretic behavior. Comparison with dihydrofolate reductases from other sources. J. Biol. Chem. 264, 2625–2633 (1989).

39. Blakley, R. L. Eukaryotic Dihydrofolate Reductase. in Advances in Enzymology and Related Areas of Molecular Biology 23–102 (John Wiley & Sons, Ltd, 1995). doi:10.1002/9780470123164.ch2.

40. Carroll, M. J. et al. Evidence for dynamics in proteins as a mechanism for ligand dissociation. Nat. Chem. Biol. 8, 246–252 (2012).

41. Sawaya, M. R. & Kraut, J. Loop and Subdomain Movements in the Mechanism of Escherichia coli Dihydrofolate Reductase: Crystallographic Evidence,. Biochemistry 36, 586–603 (1997).

42. Manna, M. S. et al. A trimethoprim derivative impedes antibiotic resistance evolution. Nat. Commun. 12, 2949 (2021).

43. Mauldin, R. V., Carroll, M. J. & Lee, A. L. Dynamic Dysfunction in Dihydrofolate Reductase Results from Antifolate Drug Binding: Modulation of Dynamics within a Structural State. Structure 17, 386–394 (2009).

44. Chen, J., Dima, R. I. & Thirumalai, D. Allosteric communication in dihydrofolate reductase: signaling network and pathways for closed to occluded transition and back. J. Mol. Biol. 374, 250–266 (2007).

45. Krucinska, J. et al. Structure-guided functional studies of plasmid-encoded dihydrofolate reductases reveal a common mechanism of trimethoprim resistance in Gram-negative pathogens. Commun. Biol. 5, 1–14 (2022).

46. Rodriguez, D. C. P., Weber, K. C., Sundberg, B. & Glasgow, A. MAGPIE: An interactive tool for visualizing and analyzing protein–ligand interactions. Protein Sci. 33, e5027 (2024).

47. Bank, R. P. D. RCSB PDB - 2W3A: HUMAN DIHYDROFOLATE REDUCTASE COMPLEXED WITH NADPH AND TRIMETHOPRIM. https://www.rcsb.org/structure/2W3A.

48. Dion-Schultz, A. & Howell, E. E. Effects of insertions and deletions in a beta-bulge region of Escherichia coli dihydrofolate reductase. Protein Eng. Des. Sel. 10, 263–272 (1997).

49. Bullerjahn, A. M. & Freisheim, J. H. Site-directed deletion mutants of a carboxyl-terminal region of human dihydrofolate reductase. J. Biol. Chem. 267, 864–870 (1992).

50. Ratnam, M., Tan, X., Prendergast, N. J., Smith, P. L. & Freisheim, J. H. Ligand-induced structural constraints in human dihydrofolate reductase revealed by peptide-specific antibodies. Biochemistry 27, 4800–4804 (1988).

51. Howell, E. E., Booth, C., Farnum, M., Kraut, J. & Warren, M. S. A second-site mutation at phenylalanine-137 that increases catalytic efficiency in the mutant aspartate-27 .fwdarw. serine Escherichia coli dihydrofolate reductase. Biochemistry 29, 8561–8569 (1990).

52. Brown, K. A., Howell, E. E. & Kraut, J. Long-range structural effects in a second-site revertant of a mutant dihydrofolate reductase. Proc. Natl. Acad. Sci. 90, 11753–11756 (1993).

53. Dion, A., Linn, C. E., Bradrick, T. D., Georghiou, S. & Howell, E. E. How do mutations at phenylalanine-153 and isoleucine-155 partially suppress the effects of the aspartate-27 .fwdarw. serine mutation in Escherichia coli dihydrofolate reductase? Biochemistry 32, 3479–3487 (1993).

54. Manna, M. S. et al. A trimethoprim derivative impedes antibiotic resistance evolution. Nat. Commun. 12, 2949 (2021).

55. Watson, M., Liu, J.-W. & Ollis, D. Directed evolution of trimethoprim resistance in Escherichia coli. FEBS J. 274, 2661–2671 (2007).

56. Toprak, E. Evolutionary paths to strong antibiotic resistance under dynamically sustained drug stress. Nat. Genet. (2011).

57. Daber, R., Stayrook, S., Rosenberg, A. & Lewis, M. Structural analysis of lac repressor bound to allosteric effectors. J. Mol. Biol. 370, 609–619 (2007).

58. Glasgow, A. et al. Ligand-specific changes in conformational flexibility mediate long-range allostery in the lac repressor. Nat. Commun. 14, 1179 (2023).

59. Markiewicz, P., Kleina, L. G., Cruz, C., Ehret, S. & Miller, J. H. Genetic studies of the lac repressor. XIV. Analysis of 4000 altered Escherichia coli lac repressors reveals essential and non-essential residues, as well as ‘spacers’ which do not require a specific sequence. J. Mol. Biol. 240, 421–433 (1994).

60. Suckow, J. et al. Genetic Studies of the Lac Repressor XV: 4000 Single Amino Acid Substitutions and Analysis of the Resulting Phenotypes on the Basis of the Protein Structure. J. Mol. Biol. 261, 509–523 (1996).

61. Romanuka, J. et al. Genetic switching by the Lac repressor is based on two-state Monod–Wyman– Changeux allostery. Proc. Natl. Acad. Sci. 120, e2311240120 (2023).

62. Kalodimos, C. G., Folkers, G. E., Boelens, R. & Kaptein, R. Strong DNA binding by covalently linked dimeric Lac headpiece: Evidence for the crucial role of the hinge helices. Proc. Natl. Acad. Sci. 98, 6039– 6044 (2001).

63. Zhong, E. D., Bepler, T., Berger, B. & Davis, J. H. CryoDRGN: reconstruction of heterogeneous cryo-EM structures using neural networks. Nat. Methods 18, 176–185 (2021).

64. Lin, Z. et al. Evolutionary-scale prediction of atomic-level protein structure with a language model. Science 379, 1123–1130 (2023).

65. van Kempen, M. et al. Fast and accurate protein structure search with Foldseek. Nat. Biotechnol. 42, 243– 246 (2024).

66. Halabi, N., Rivoire, O., Leibler, S. & Ranganathan, R. Protein Sectors: Evolutionary Units of Three-Dimensional Structure. Cell 138, 774–786 (2009).

67. Steinegger, M. & Söding, J. MMseqs2 enables sensitive protein sequence searching for the analysis of massive data sets. Nat. Biotechnol. 35, 1026–1028 (2017).

68. Leaver-Fay, A. et al. ROSETTA3: an object-oriented software suite for the simulation and design of macromolecules. in Methods in enzymology vol. 487 545–574 (Elsevier, 2011).

69. Jumper, J. et al. Highly accurate protein structure prediction with AlphaFold. Nature 596, 583–589 (2021).

70. Goloborodko, A. A., Levitsky, L. I., Ivanov, M. V. & Gorshkov, M. V. Pyteomics—a Python Framework for Exploratory Data Analysis and Rapid Software Prototyping in Proteomics. J. Am. Soc. Mass Spectrom. 24, 301–304 (2013).

71. Levitsky, L. I., Klein, J. A., Ivanov, M. V. & Gorshkov, M. V. Pyteomics 4.0: Five Years of Development of a Python Proteomics Framework. J. Proteome Res. 18, 709–714 (2019).

72. Skinner, J. J., Lim, W. K., Bédard, S., Black, B. E. & Englander, S. W. Protein dynamics viewed by hydrogen exchange. Protein Sci. 21, 996–1005 (2012).

73. Smit, J. & roy. Jhsmit/HDXrate: HDXrate version 0.2.2. Zenodo 10.5281/zenodo.10022160 (2023).

74. Nguyen, D., Mayne, L., Phillips, M. C. & Walter Englander, S. Reference Parameters for Protein Hydrogen Exchange Rates. J. Am. Soc. Mass Spectrom. 29, 1936–1939 (2018).

75. Röst, H. L., Schmitt, U., Aebersold, R. & Malmström, L. pyOpenMS: A Python-based interface to the OpenMS mass-spectrometry algorithm library. PROTEOMICS 14, 74–77 (2014).

76. Bystroff, C. & Kraut, J. Crystal structure of unliganded Escherichia coli dihydrofolate reductase. Ligand-induced conformational changes and cooperativity in binding. Biochemistry 30, 2227–2239 (1991).

77. Bell, C. E. & Lewis, M. A closer view of the conformation of the Lac repressor bound to operator. Nat. Struct. Biol. 7, 209–214 (2000).

78. Zhang, C., Shine, M., Pyle, A. M. & Zhang, Y. US-align: universal structure alignments of proteins, nucleic acids, and macromolecular complexes. Nat. Methods 19, 1109–1115 (2022).

79. Manna, M. S. et al. A trimethoprim derivative impedes antibiotic resistance evolution. Nat. Commun. 12, 2949 (2021).

80. Bell, C. E. & Lewis, M. A closer view of the conformation of the Lac repressor bound to operator. Nat. Struct. Biol. 7, 209–214 (2000).

81. Perez-Riverol, Y. et al. The PRIDE database resources in 2022: a hub for mass spectrometry-based proteomics evidences. Nucleic Acids Res. 50, D543–D552 (2022).

