## Supplementary figures and images for "Site-resolved energetic information from HX/MS experiments"

### DHFR_uptake_errors_MTX_0.pdf

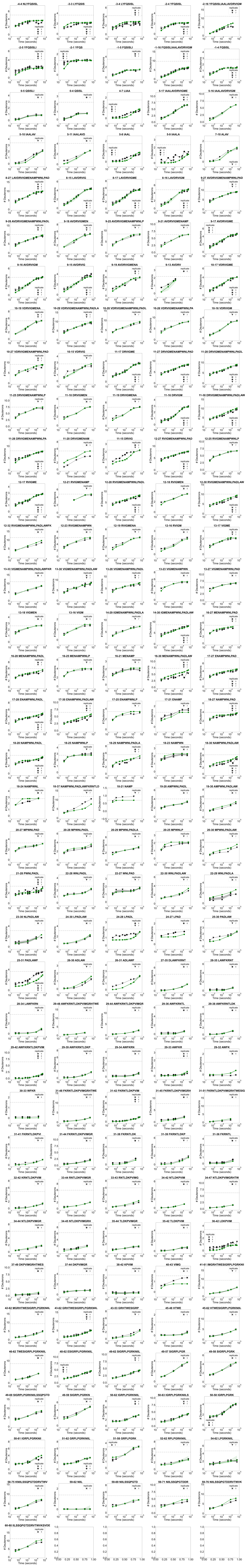

### DHFR_uptake_errors_MTX_1.pdf

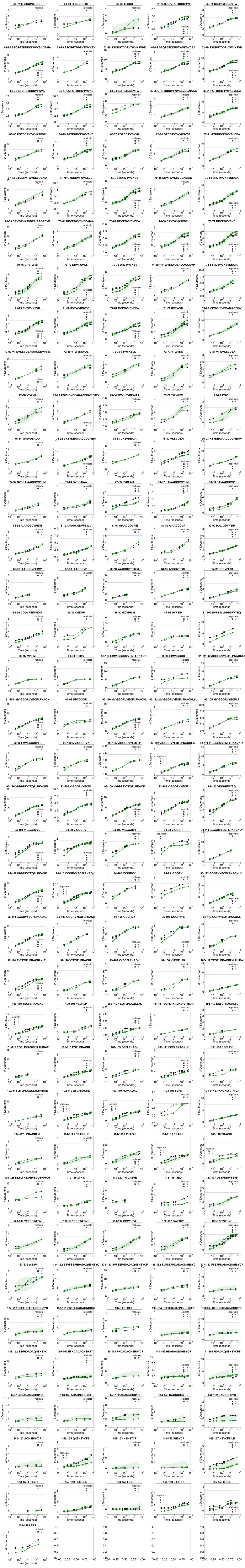

### DHFR_uptake_errors_TMP_0.pdf

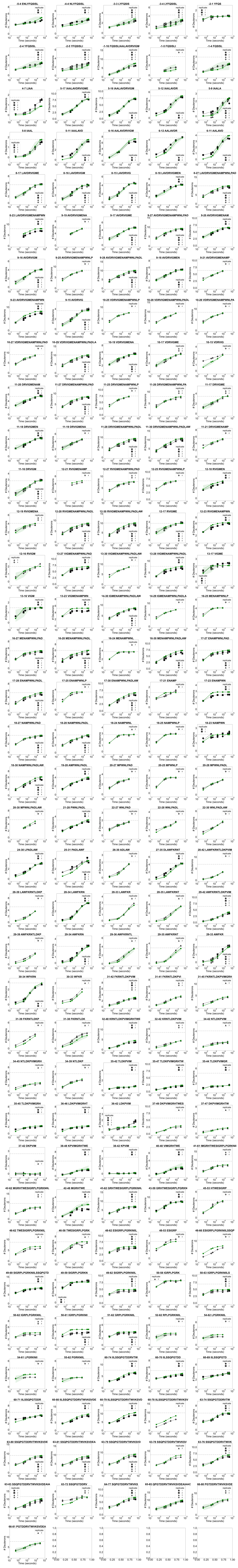

### DHFR_uptake_errors_TMP_1.pdf

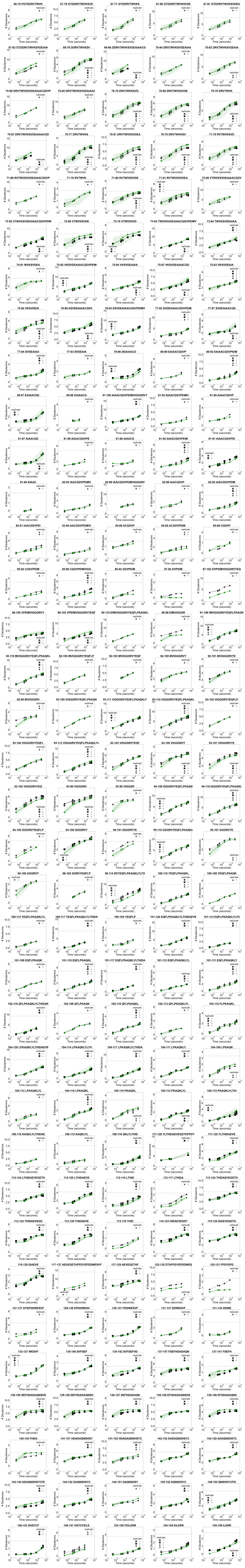

### hDHFR_exp_uptake_1.pdf

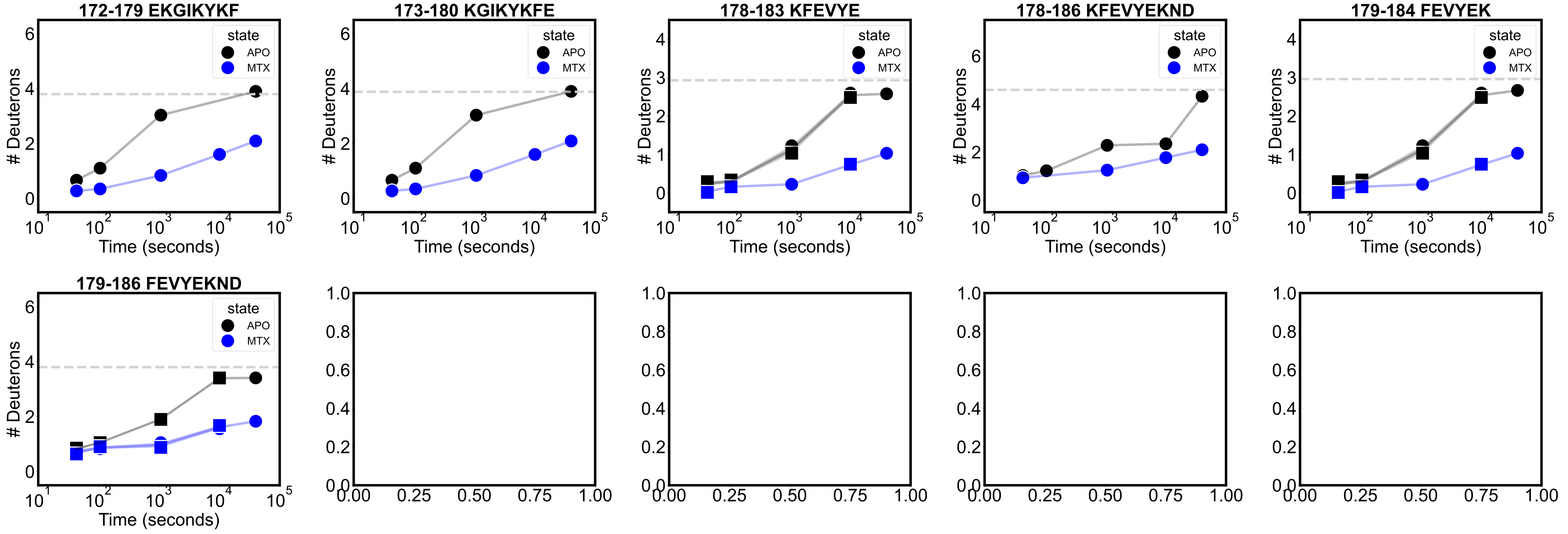

### LacI_exp_uptake_1.pdf

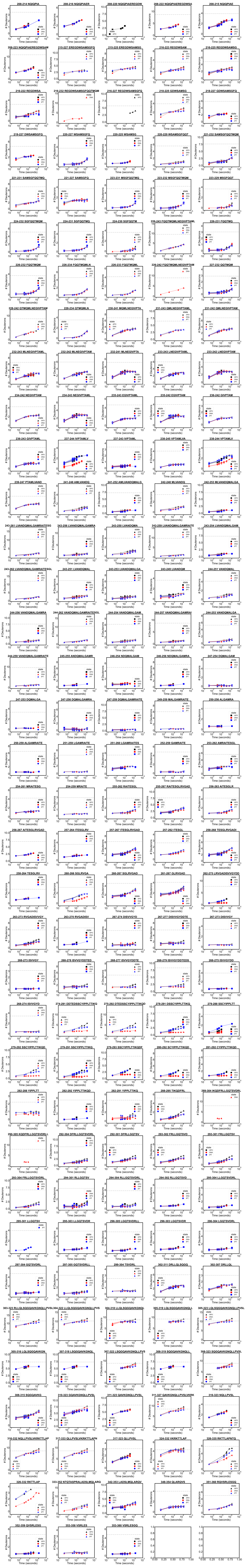
