## Supplemental information for "Site-resolved energetic information from HX/MS experiments"

[1] Department of Biochemistry and Molecular Biophysics, Columbia University Irving Medical Center, New York, NY, United States
* These authors made equal contributions.
† Corresponding author

### 1. Review of hydrogen exchange theory

In the HX/MS experiments described in this work, we measured the rate of hydrogen (H) exchange for deuterium (D) in the backbone amide groups of proteins at equilibrium. The H-D exchange reaction occurs when transient opening of the protein backbone exposes the amide group to solvent.

$$\mathrm{NH(closed)} \underset{k_{cl}}{\overset{k_{op}}{\rightleftharpoons}} \mathrm{NH(open)} \overset{k_{int}}{\rightleftharpoons} \mathrm{exchanged}$$

All canonical amino acids except for prolines are involved in backbone amide H bonds in the protein structure and can therefore be measured in the experiment.

Under the steady-state approximation:

$k_{ex} = k_{int}k_{op}/(k_{int}+k_{cl}+k_{op}) \approx k_{int}k_{op}/(k_{int}+k_{cl})$

In this reaction, $K_{op}$, defined as $(k_{op}/k_{cl})$, is the opening equilibrium constant. $k_{int}$ is the intrinsic or chemical exchange rate for the amino acid in the context of its neighbors in an unstructured polypeptide. The $k_{int}$ rates were previously experimentally determined.[1]

In folded proteins at equilibrium, the hydrogen exchange reaction occurs as the protein rapidly unfolds and refolds or experiences local fluctuations in the conformational ensemble, allowing each hydrogen to exchange individually. In this EX2 regime, $k_{cl} >> k_{int}$ because the fold prevents immediate rapid hydrogen exchange.

$K_{op}$ and $k_{int}$ are related by the H-D exchange rate, $k_{ex}$, that is observed in the experiment at the EX2 regime (where $K_{op} << 1$, favoring the closed state).

In the EX2 regime, therefore,

$k_{ex} = K_{op}k_{int}$.

With knowledge of these terms, we can determine the free energy of opening, $\Delta G_{op}$, for each amino acid.

$\Delta G_{op} = -RT\ln(K_{op}) = RT\ln(k_{int}/k_{ex})$.

(The ratio $k_{int}/k_{ex}$ is also known as the protection factor, PF).

$\Delta G_{op}$ can be used to determine the Gibbs free energy for the whole molecule unfolding, or $\Delta G_{unf}$. $\Delta G_{unf}$ contributes to $\Delta G_{op}$ together with the free energy of local fluctuations that are not denaturant-dependent, $\Delta G_{fl}$.

The dependence of $\Delta G_{unf}$ on denaturant concentration, $\Delta G_{unf}(den)$, is assumed to be linear.[2,3] The slowest-exchanging hydrogens are typically involved in major unfolding transitions, and therefore have strong denaturant dependence. For these hydrogens, $\Delta G_{op}$ is well-correlated with the equilibrium global unfolding constant $K_u$ and linearly correlated with the denaturant concentration [den].

$\Delta G_{unf}(den) = \Delta G_{unf}(0) - m[den]$

At low denaturant concentrations, however, the exchange of many hydrogens depends more on local structural fluctuations, leading to low *m* values near zero.

The observed H-D exchange rate depends on both local fluctuations and global unfolding.
$k_{ex} = [K_{op}(local) + K_{op}(global)]k_{int}$

Therefore,

$\Delta G_{op} = -RT\ln(K_{unf}+K_{fl})$

where

$K_{unf} = \exp(-\Delta G_{unf} - m[den]/RT)$
$K_{fl} = \exp(-\Delta G_{fl}/RT)$.

By repeating HX/MS with PIGEON-FEATHER at several denaturant concentrations, one can determine the contribution of the $\Delta G_{unf}$ and $\Delta G_{fl}$ terms to $\Delta G_{op}$.

Summarized from:
1. Bai *et al.*, 1995[4]
2. Chamberlain *et al.*, 1996[5]
3. Hollien and Marqusee, 1999[6]

### 2. Recommendations for PIGEON-FEATHER usage

The benchmarks and case studies in this work inform guidelines for HX/MS-PIGEON-FEATHER usage and data interpretation. Including more data with longer timepoints and deeper, broader sequence coverage improved resolution and accuracy for all proteins. Accordingly, we suggest that technical replicates should be pooled rather than processed individually and averaged.

We recommend PIGEON-FEATHER for single-site $\Delta G_{op}$ calculations rather than peptide-level analysis for protein domains if:
(1) peptide-level sequence coverage >90%;
(2) single-site resolution >50%;
(3) >50% peptides have exchange rates that fall within the timescale of the experiment;
(4) there are at least half as many peptides as amino acids in the protein; and
(5) the protein region of interest is in the EX2 rather than the EX1 or EXX regime.

Otherwise, the peptide-level analysis and visualization tools implemented in PIGEON-FEATHER provide a more appropriate option.

HX/MS experimental design can also be optimized for PIGEON-FEATHER. Before performing HX/MS, the user can optimize sample preparation conditions, the sample concentration, buffers, and instrument settings until the best PIGEON-generated peptide list is achieved, in terms of the number of unique peptides and the peptide-level sequence coverage. We find that performing replicates with multiple proteases can improve both metrics. To accurately capture the exchange dynamics, if possible, one should perform HX/MS such that at least half of the peptides are fully exchanged by the last timepoint, as confirmed by a fully deuterated control experiment.

### 3. Supplemental figures

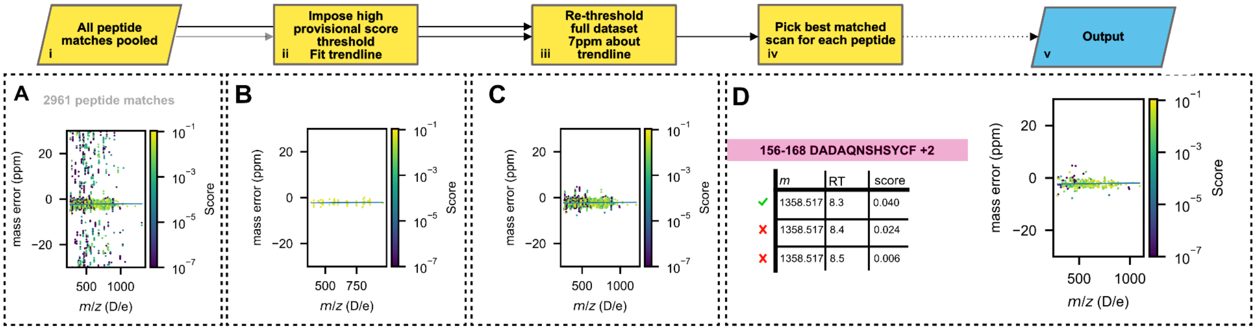

**Figure S1. PIGEON corrects systematic *m/z* error.**

**A.** Example input dataset (Extended Data Fig. 1f). **B.** Heuristic inverse square root fit to a restricted dataset with high confidence peptides (score >0.05). **C.** Impose cut on the full dataset (default: 7 ppm) about heuristic trendline. **D.** Pick the consensus match for each peptide by choosing the peak with highest fragment score (ties broken by choosing highest peak intensity). The output of this calculation is passed to the peptide disambiguation step (Extended Data Fig. 1e, iii).

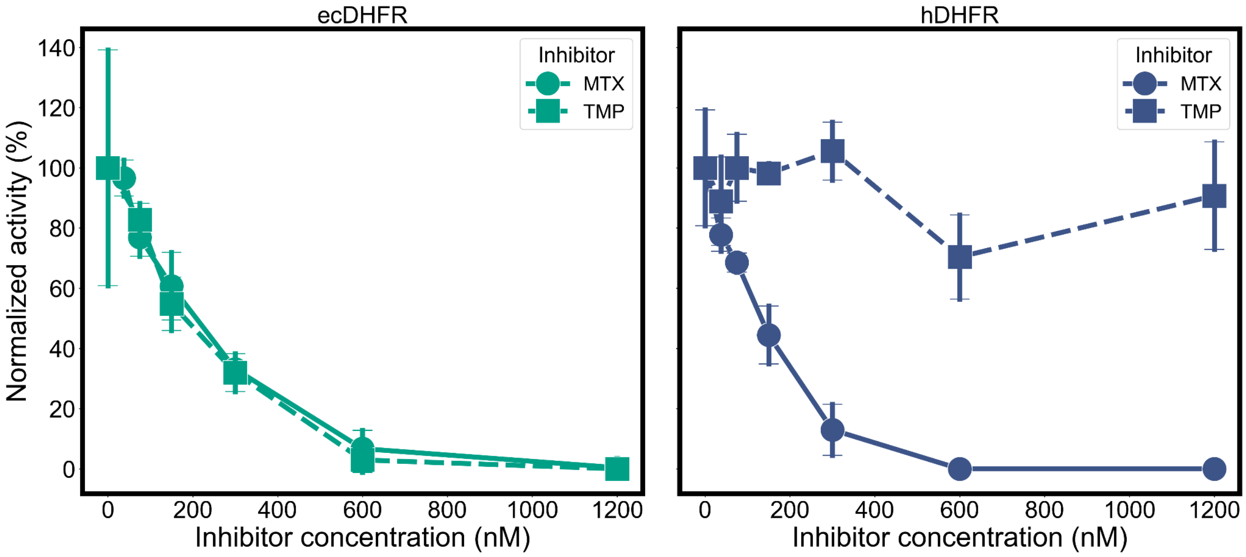

**Figure S2. DHFR activity assay.**

The reaction was conducted in a buffer containing 50 mM HEPES, 150 mM NaCl, 1 mM TCEP, and 7.5% glycerol, 100 nM ecDHFR, 100 μM NADPH, 32 μM DHF, pH 7.0, and a specific amount of the inhibitor (37.5-1200 nM, details in Supplemental methods).

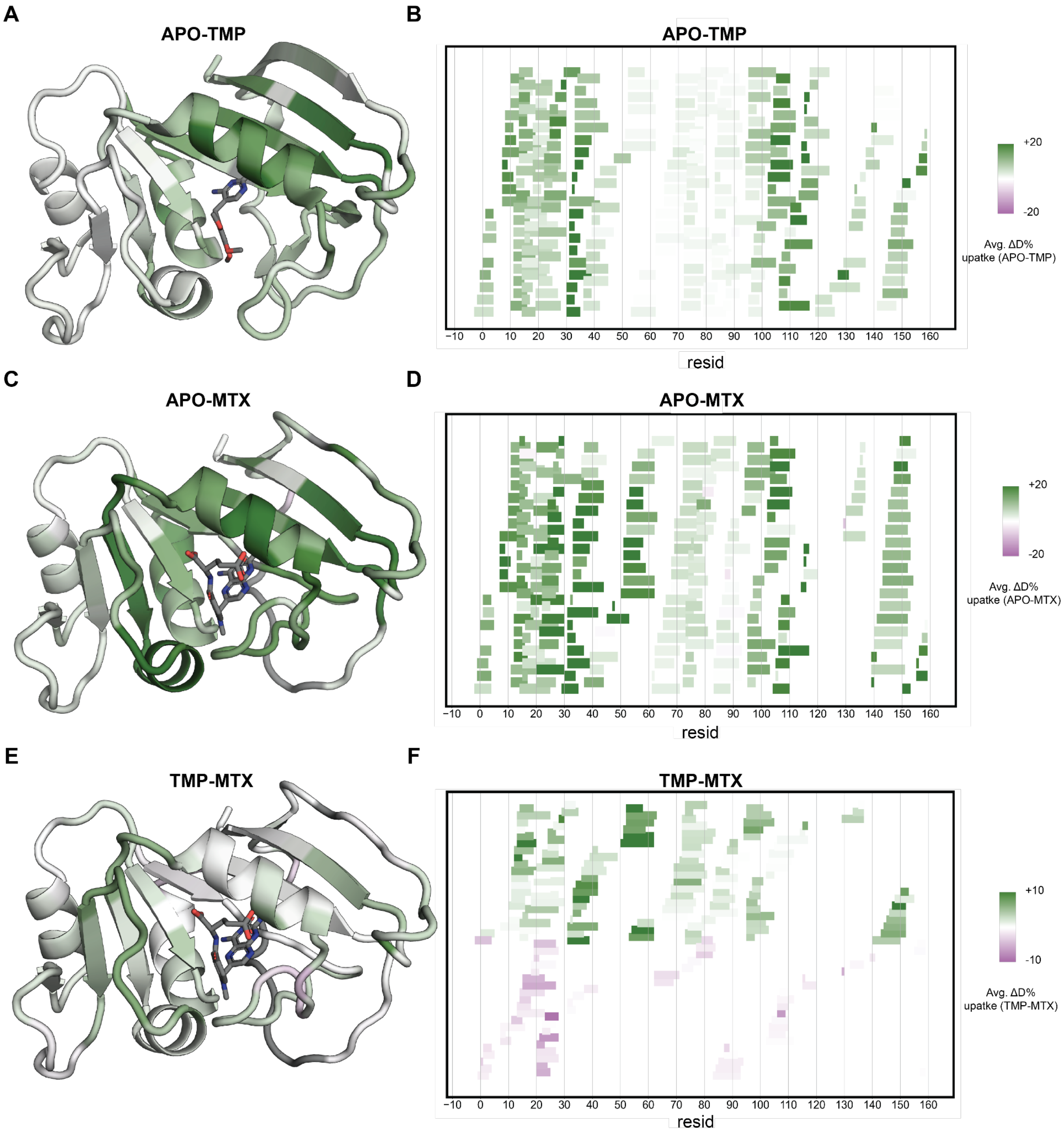

**Figure S3. Peptide level analysis of ecDHFR datasets.**

**A, C, E.** Average peptide deuteration uptake percentage difference (ΔD%) between APO and TMP state, APO and MTX state, TMP state and MTX state, respectively. The ΔD% was calculated for each peptide by averaging the experimental deuterium uptake at each timepoint and dividing by the total number of exchangeable protons in the peptide. The green color of most sites indicates that the first state undergoes more H-D exchange, while the second state is more protected from exchange. **B, D, F.** Woods plots showing the ΔD% of all the peptides.

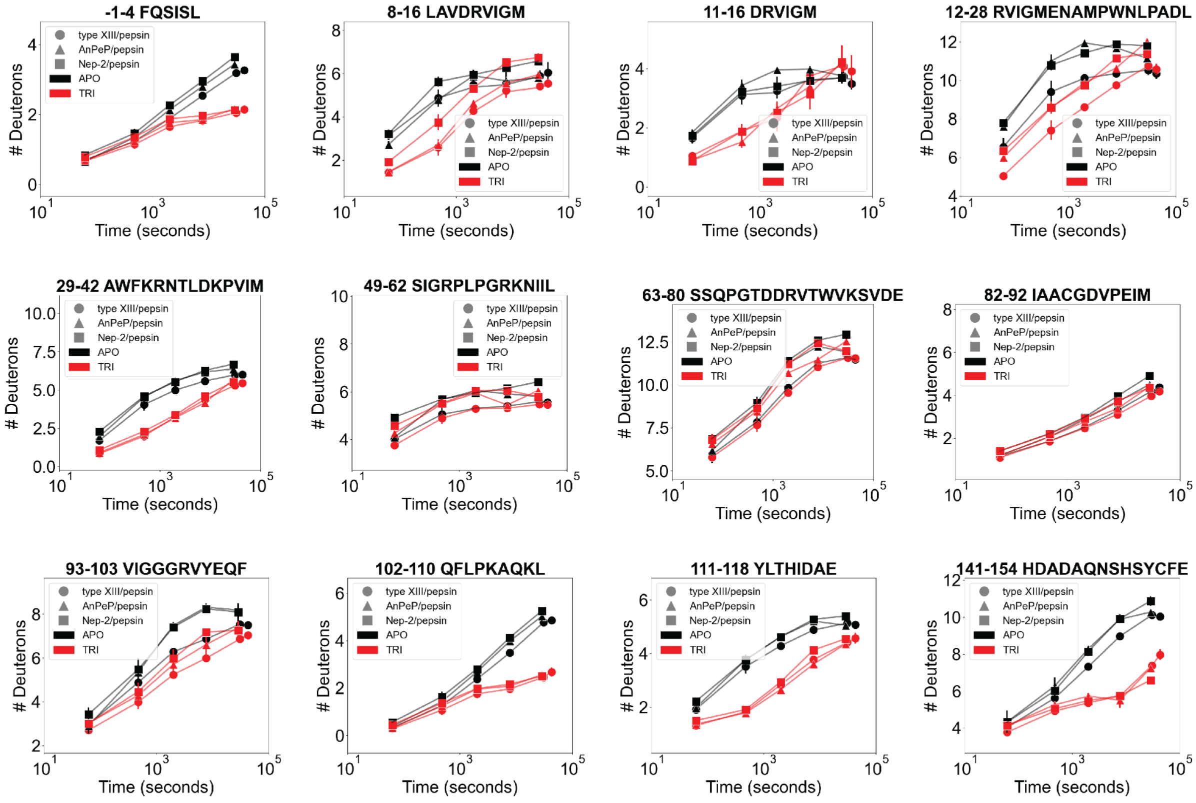

**Figure S4. Protease reproducibility.**

Examples of D uptake plots of the same ecDHFR peptides collected using three different protease digest columns. TRI, trimethoprim-bound ecDHFR.

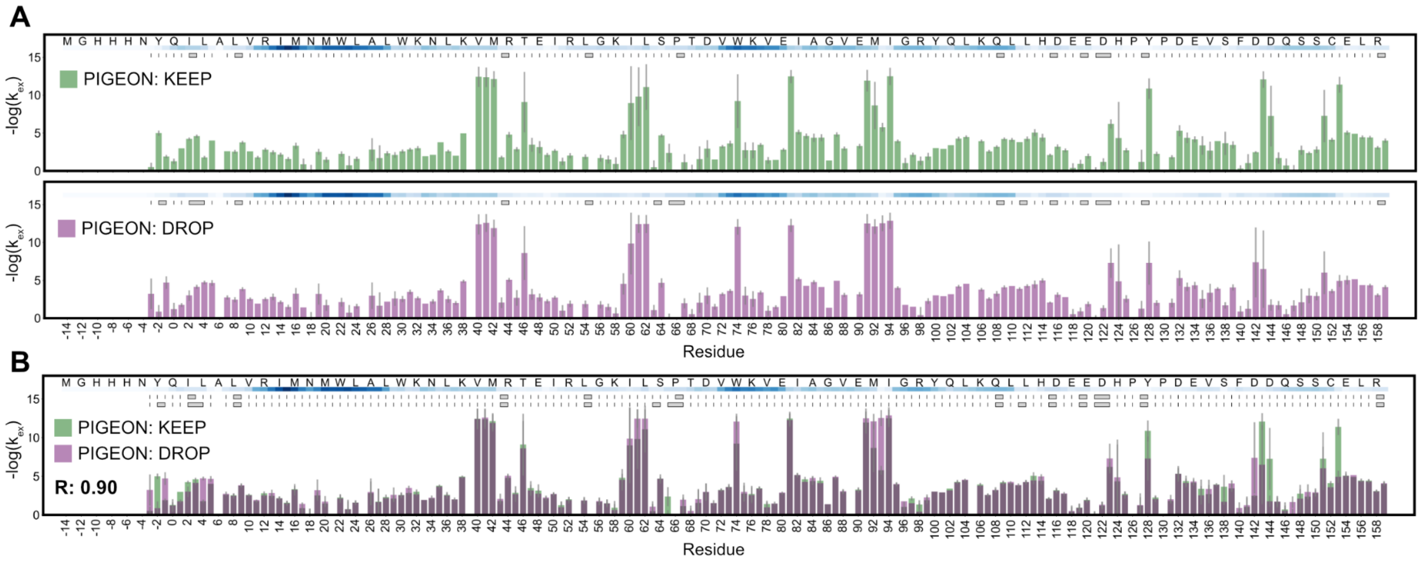

**Figure S5. The effect of PIGEON modes on FEATHER results.**

**A.** Comparison of FEATHER results with different PIGEON options for ecDHFR datasets in the APO state. **B.** Pairwise comparison of results in panel A. R values of the correlation are annotated below the legend.

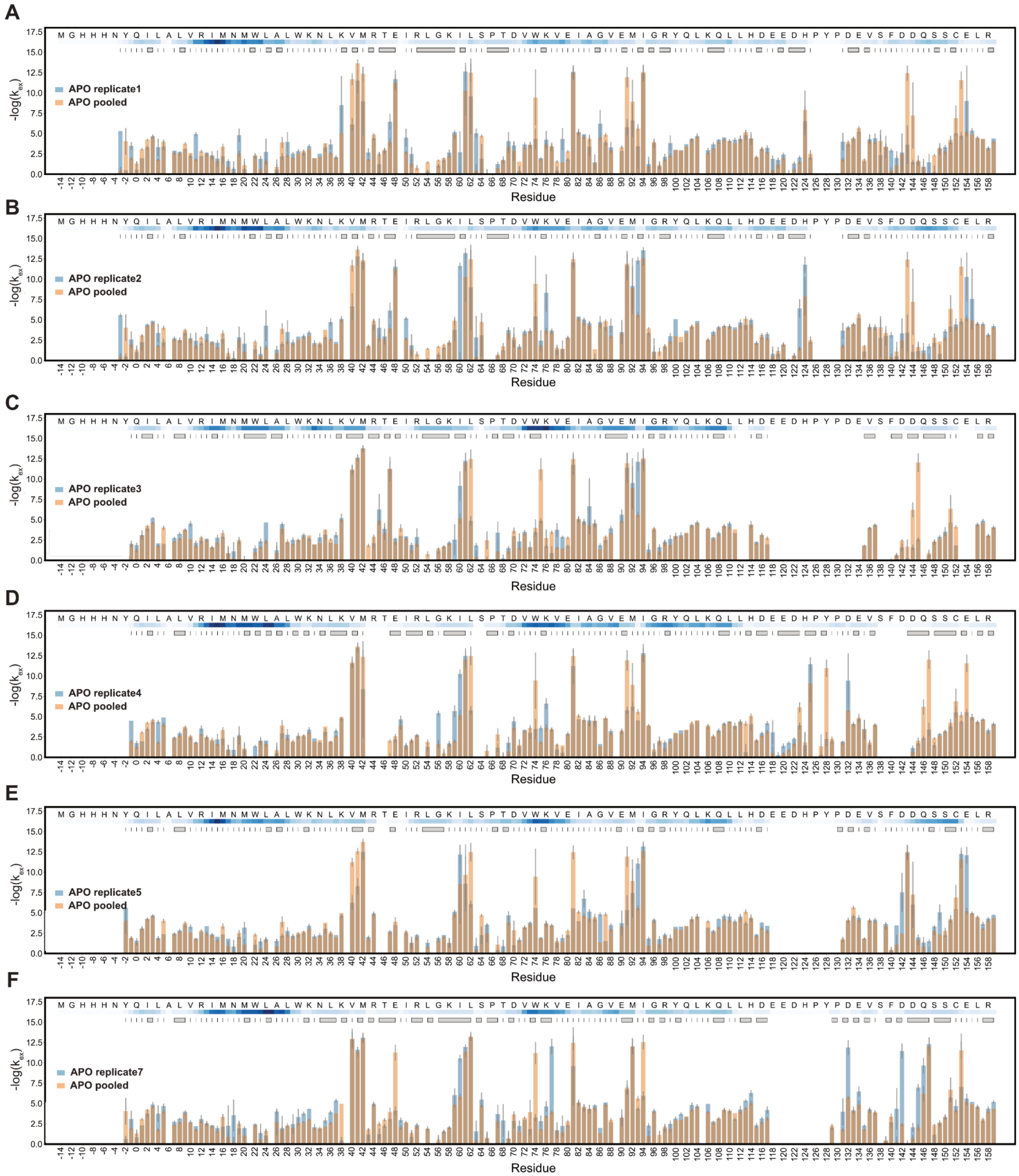

**Figure S6. Reproducibility analysis of ecDHFR datasets in the APO state.**

Comparisons between -log($k_{ex}$) from individual replicates to the pooled dataset. The residues are numbered as in PDB 6XG5.[7]

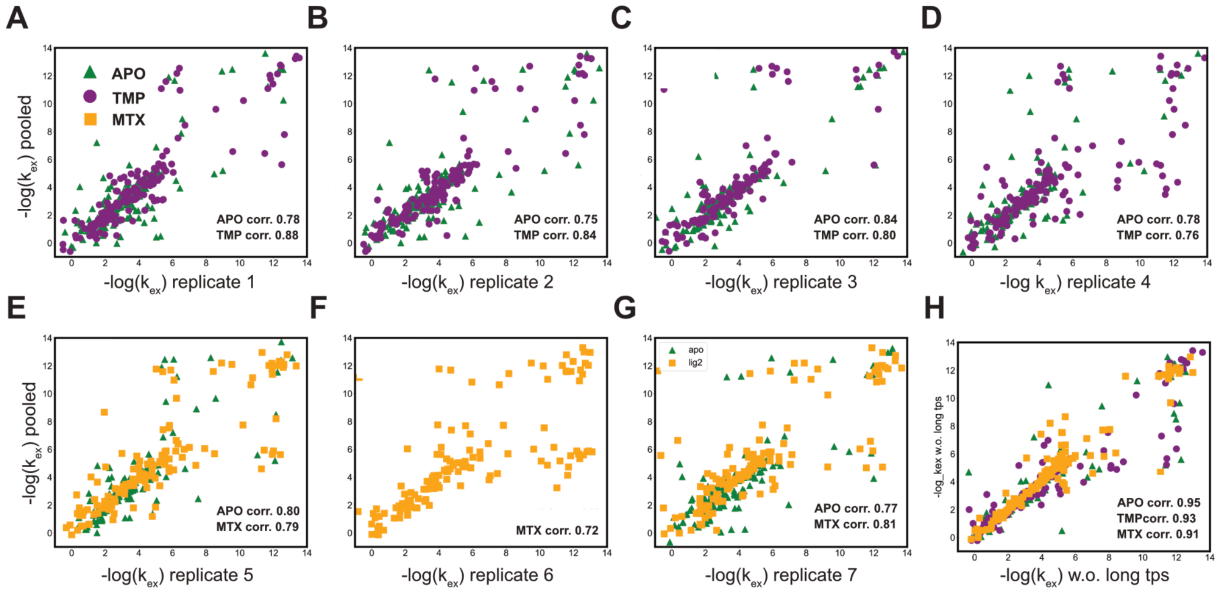

**Figure S7. Reproducibility analysis of all singlicate ecDHFR datasets vs. pooled data.**

Correlation of -log($k_{ex}$) corresponding to Figs. S6, S8, S14, S15.

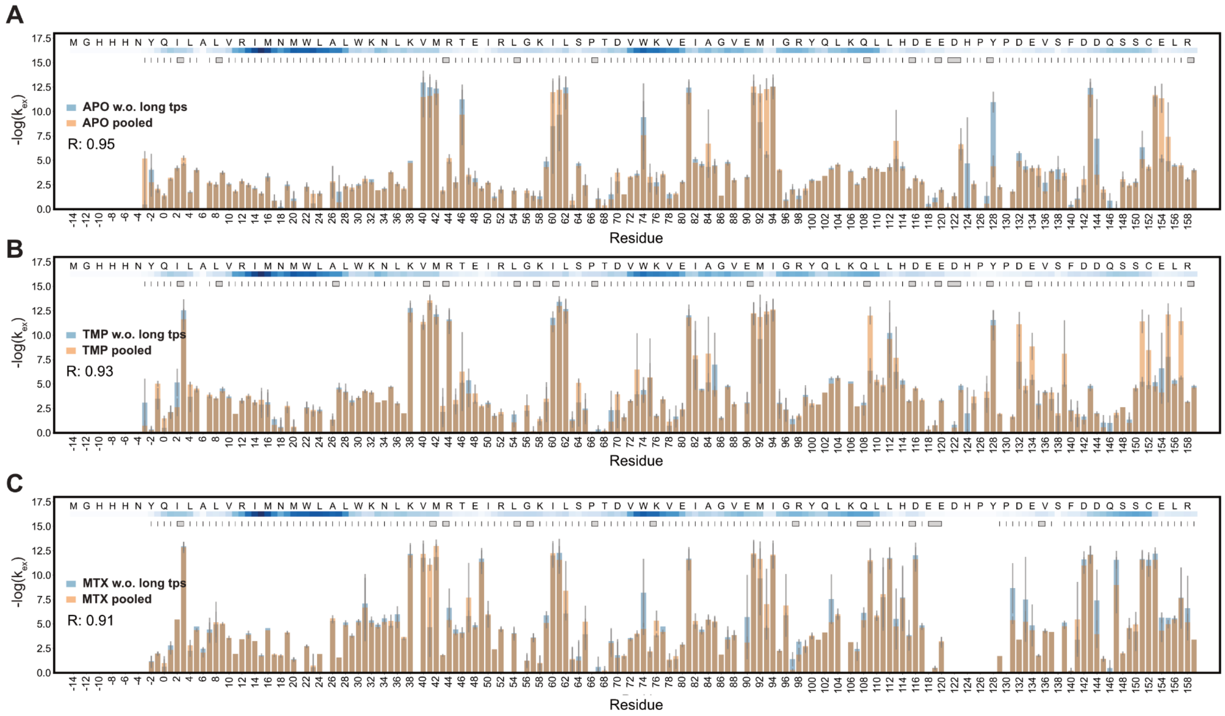

**Figure S8. The effect of including long HX time points on FEATHER results.**

Comparisons between -log($k_{ex}$) from with/without long time points (>4 hours) for ecDHFR. The residues are numbered as in PDB ID 6XG4.

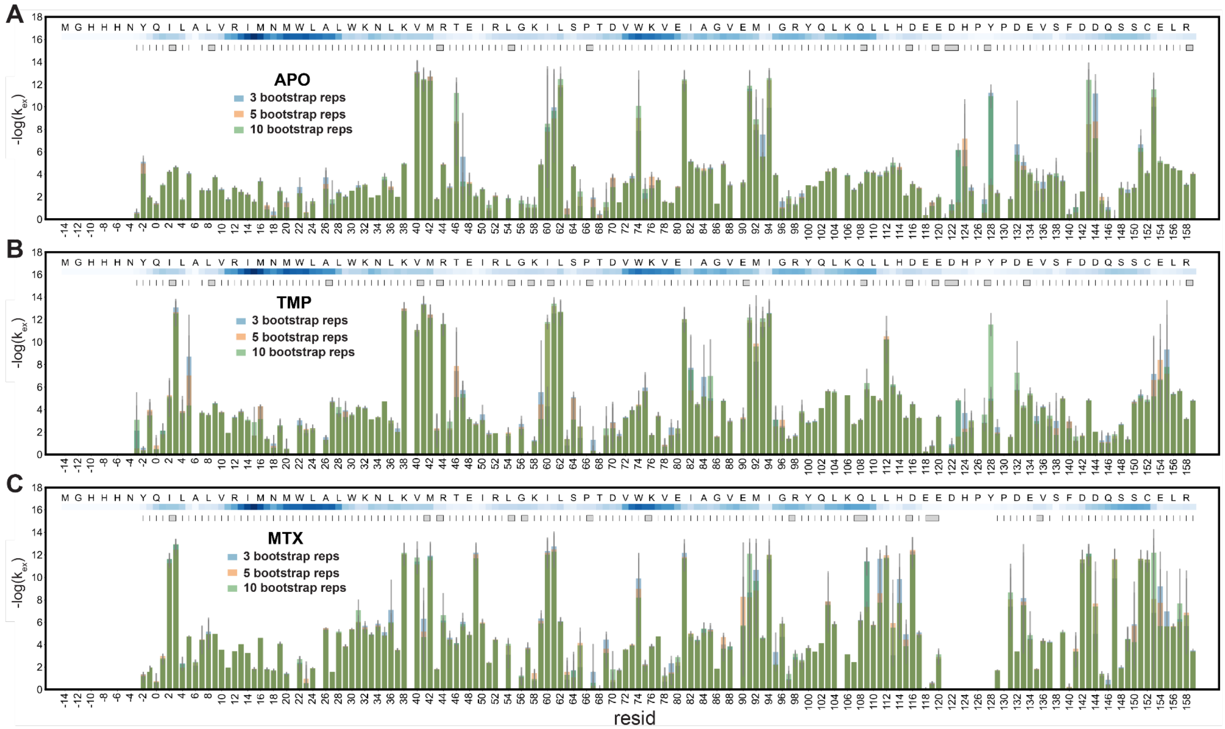

**Figure S9. The effect of the number of bootstrap replicates on FEATHER results.**

The results obtained from running PIGEON-FEATHER with 3, 5, and 10 bootstrap replicates were compared for ecDHFR APO, TMP, and MTX states, respectively. Increasing beyond 3 replicates does not yield to improved accuracy.

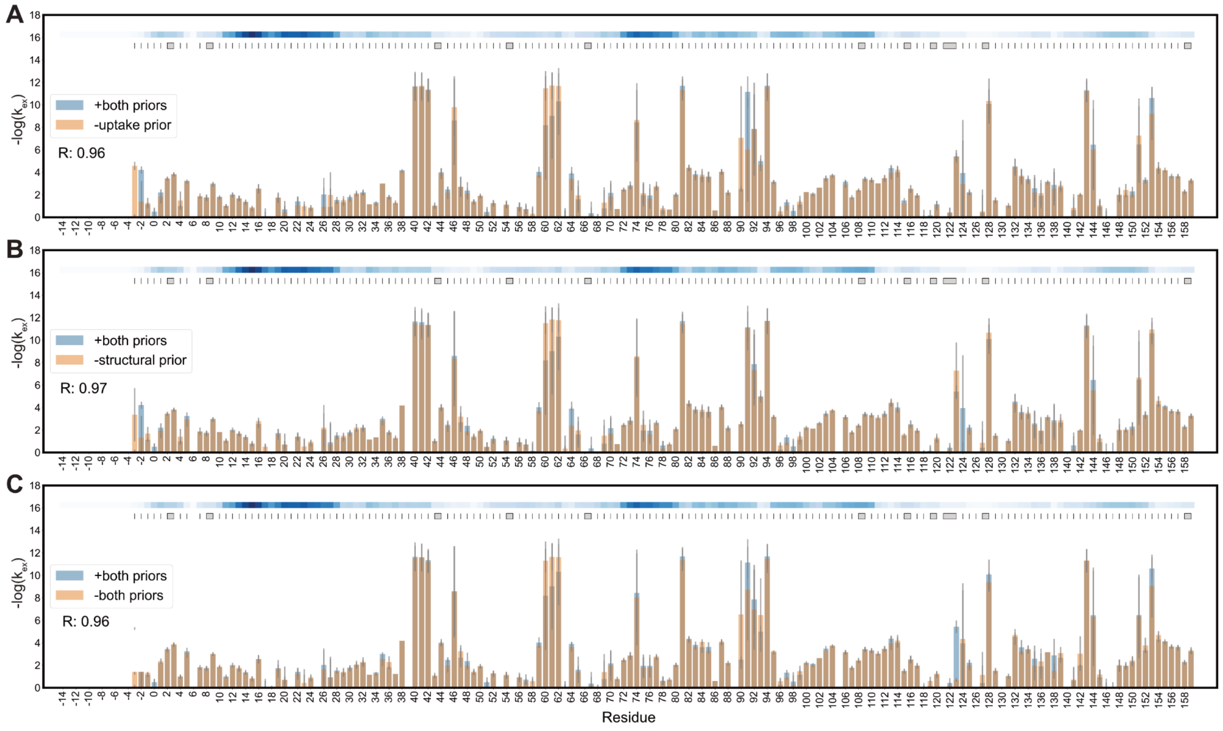

**Figure S10. The effects of including the two Bayesian priors.**

**A.-C.** The comparison of results obtained from running PIGEON-FEATHER without uptake priors, structural priors, and both priors to those obtained with both priors for the ecDHFR APO state. R values of the correlation are annotated below the legend.

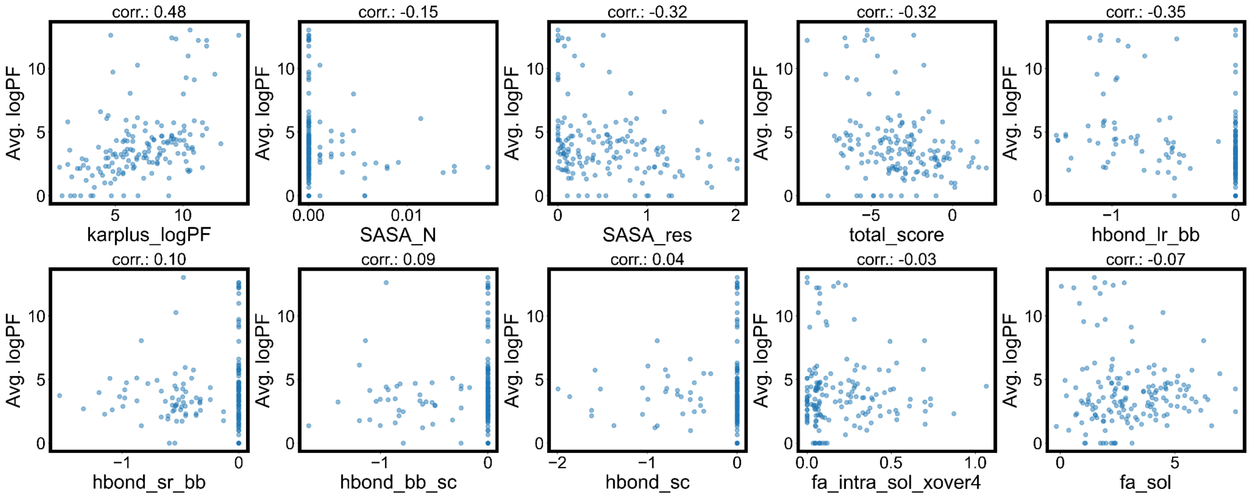

**Figure S11. Computationally determined metrics as predictors for PFs.**

The correlation analysis between PIGEON-FEATHER-determined log(PF), and log(PF) calculated using the empirical model (karplus_logPF)[8], SASA, and related Rosetta energy terms[9].

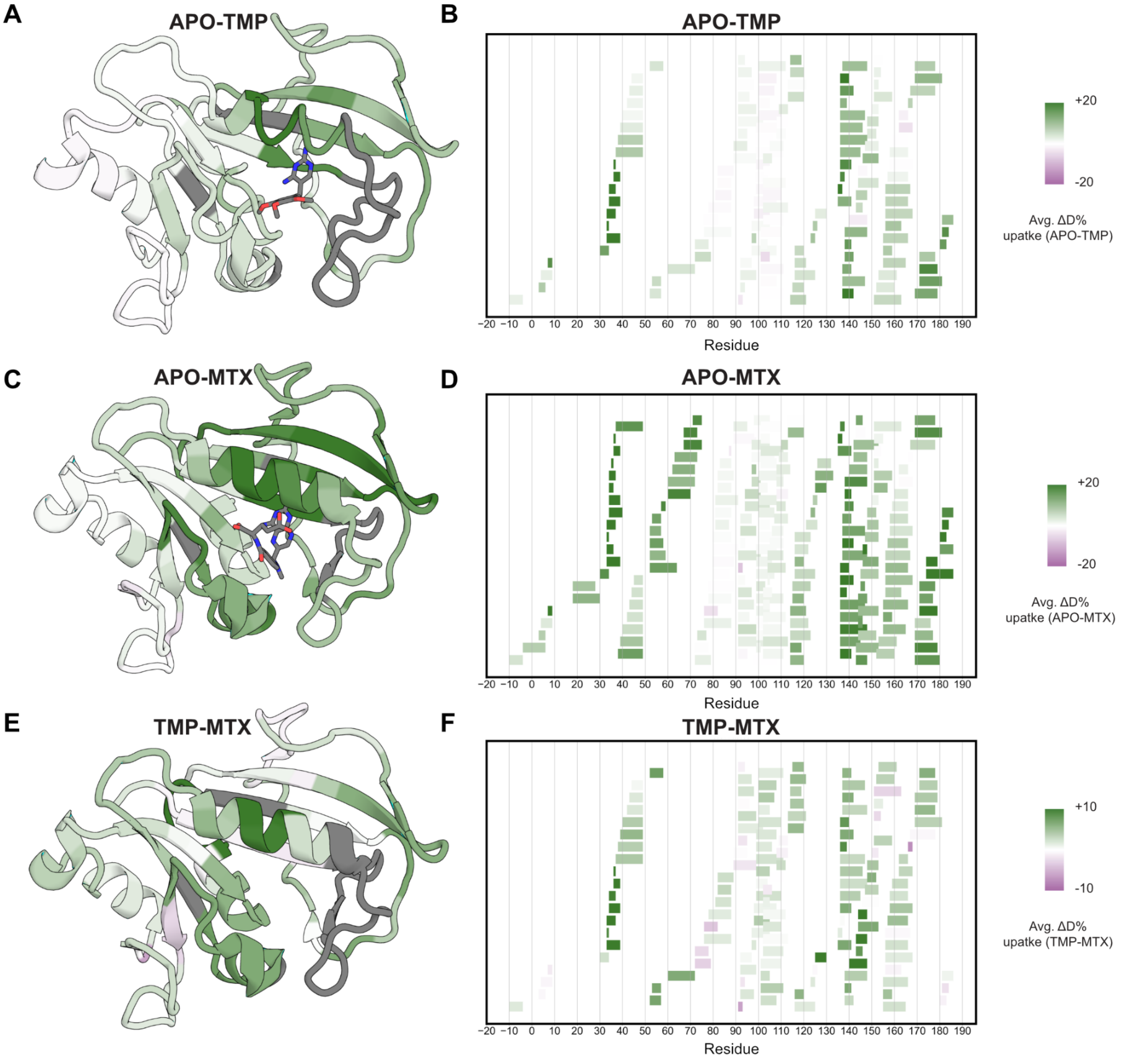

**Figure S12. Peptide level analysis of hDHFR datasets.**

**A, C, E.** Average peptide deuteration uptake percentage difference (ΔD%) between APO and TMP state, APO and MTX state, TMP state and MTX state, respectively. The ΔD% was calculated for each peptide by averaging the experimental deuterium uptake at each timepoint and dividing by the total number of exchangeable protons in the peptide. The green color of most sites indicates that the first state undergoes more H-D exchange, while the second state is more protected from exchange. **B, D, F.** Woods plots showing the ΔD% of all the peptides.

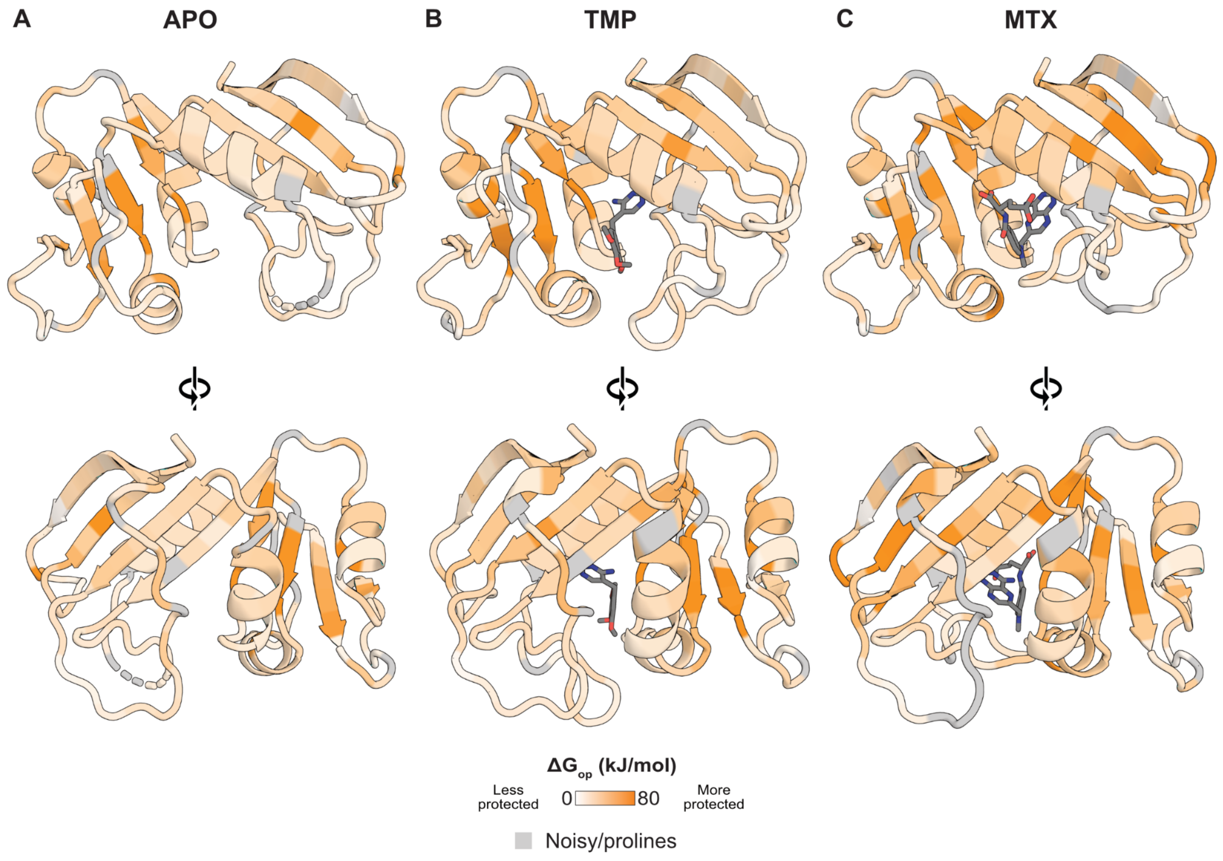

**Figure S13. Absolute $\Delta G_{op}$ for ecDHFR functional states.**

**A.** APO ecDHFR (PDB ID: 5DFR[10]). **B.** TMP-DHFR (PDB: 6XG5[7]). **C.** MTX ecDHFR (PDB ID: 1RG7[11]).

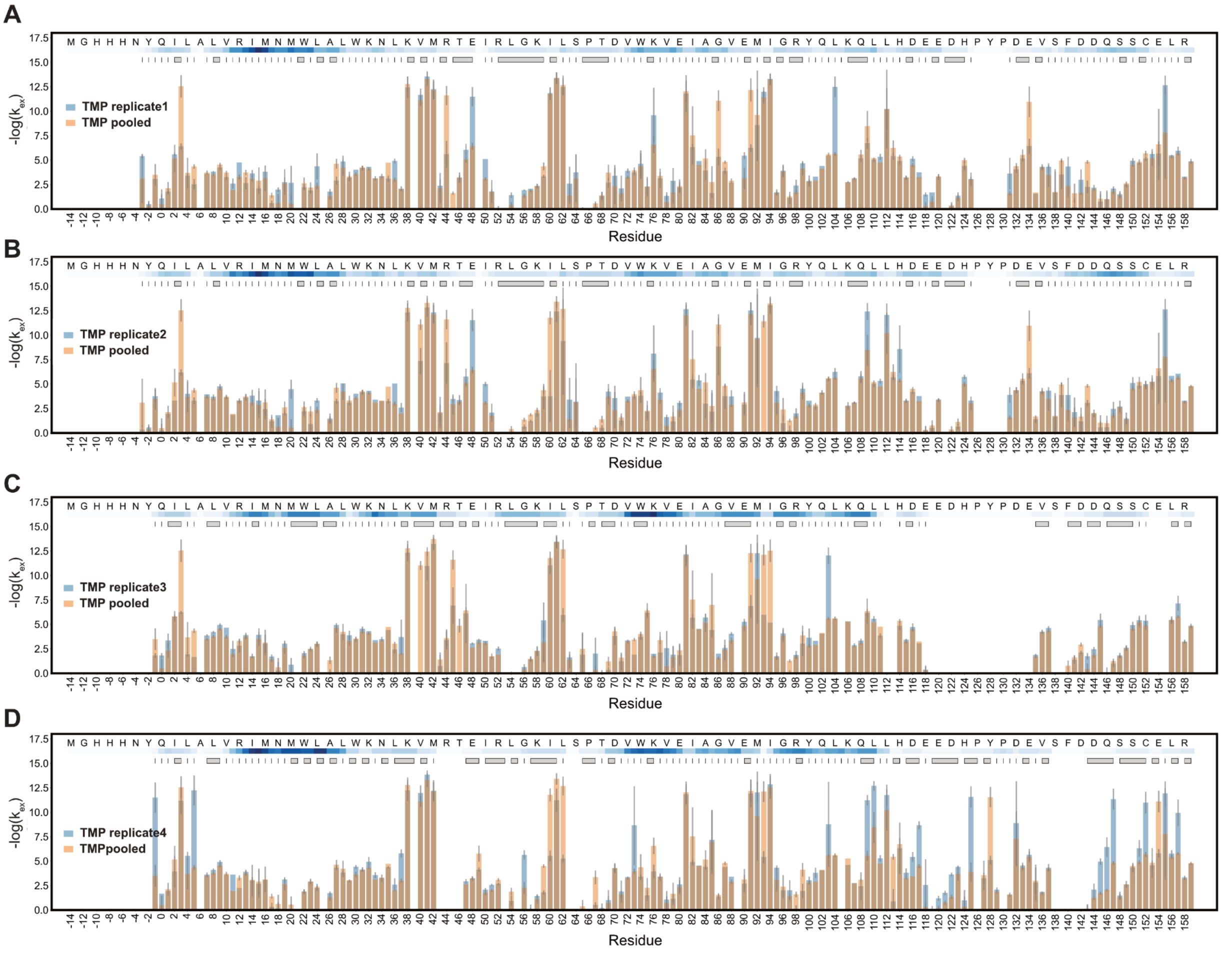

**Figure S14. Reproducibility analysis of ecDHFR datasets in the TMP state.**

Comparisons between log(PF)s from individual replicates to the pooled dataset. The residues are aligned to PDB 6XG4.

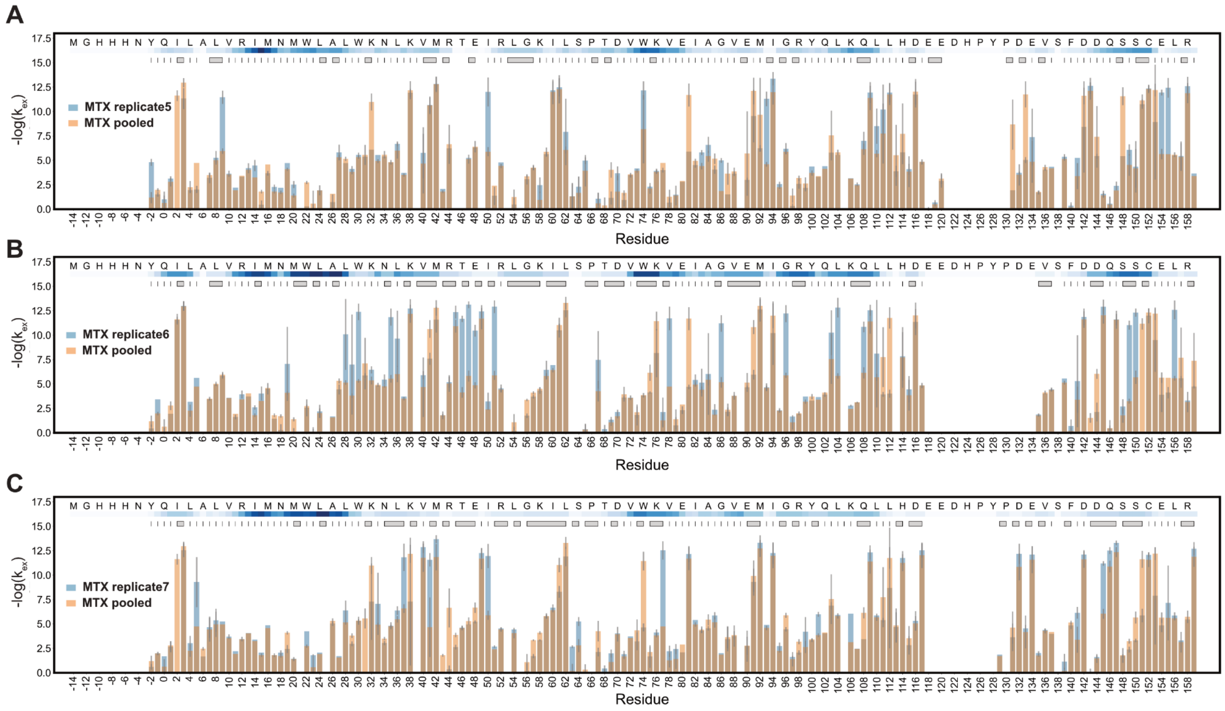

**Figure S15. Reproducibility analysis of ecDHFR datasets in the MTX state.**

Comparisons between log(PF)s from individual replicates to the pooled dataset. The residues are aligned to PDB 6XG5[7].

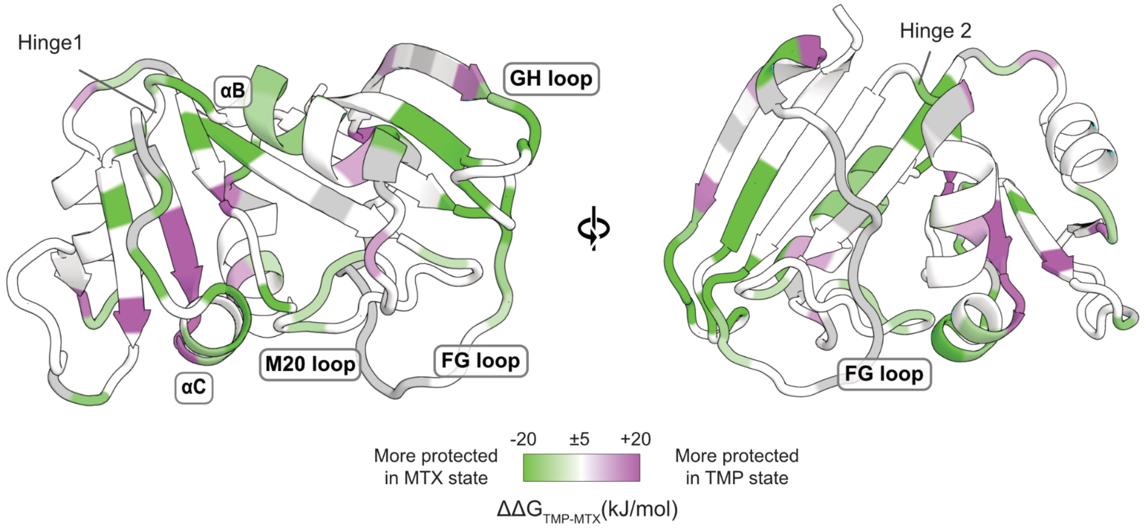

**Figure S16. Differential effects of two competitive inhibitors on ecDHFR.**

Noisy/low data regions and prolines are colored gray. PDB ID: 1RG7[11].

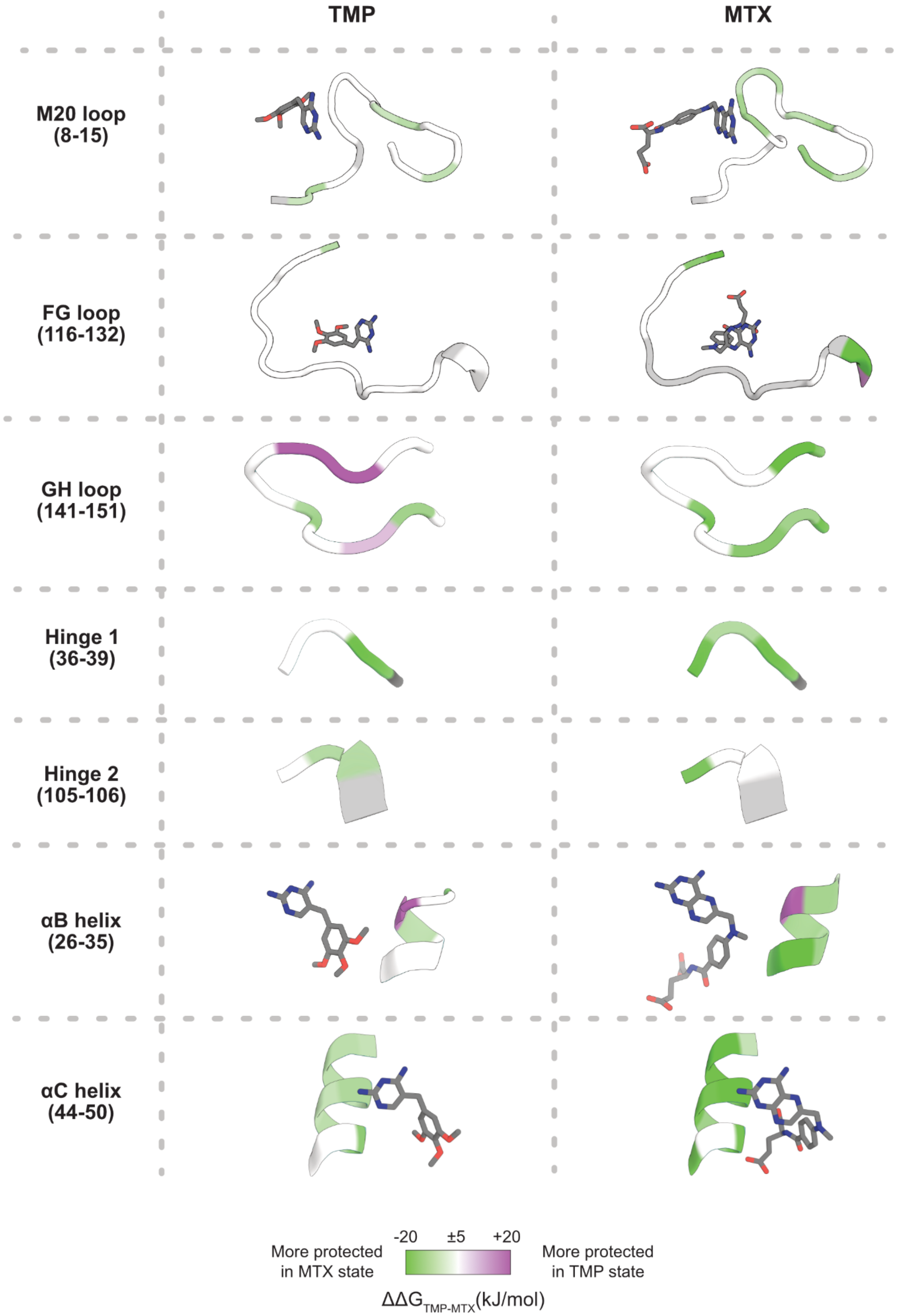

**Figure S17. Comparing inhibitor- vs. APO state ecDHFR by secondary structures.**

TMP and MTX are shown as yellow sticks in some structures. PDB ID: 1RG7[11] (MTX-ecDHFR), 6XG5[7] (TMP-ecDHFR). Noisy/low data regions and prolines are colored gray.

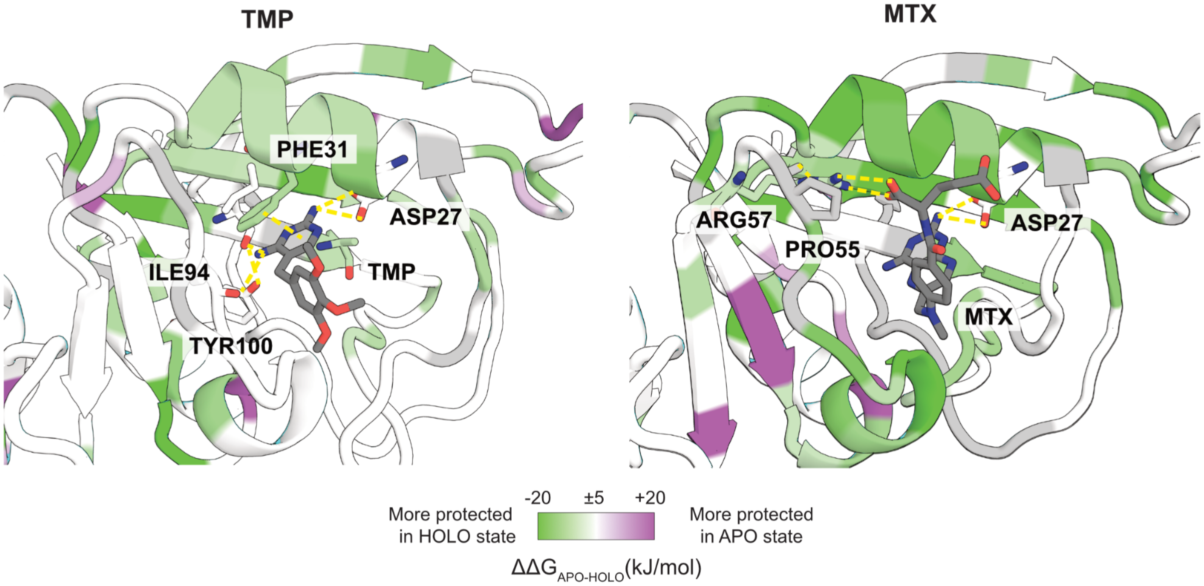

**Figure S18. Comparing inhibitor- vs. APO state ecDHFR, binding pocket.**

TMP and MTX are shown as yellow sticks. Some amino acids that directly interact with the inhibitor molecules are also shown as sticks. Hydrogen bonds shown as yellow dashes. PDB ID: 1RG7[11] (MTX-ecDHFR, top), 6XG5[7] (TMP-ecDHFR, bottom).

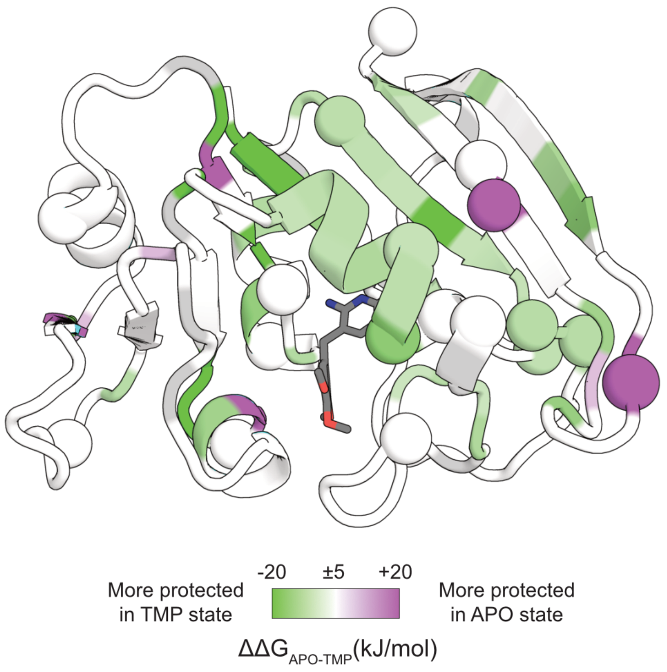

**Figure S19. TMP resistance mutations are enriched in the β sheet and binding pocket.**

Positions where TMP resistance mutations commonly occur are shown as spheres. Noisy/low data regions and prolines are colored gray. PDB ID: 1RG7[11].

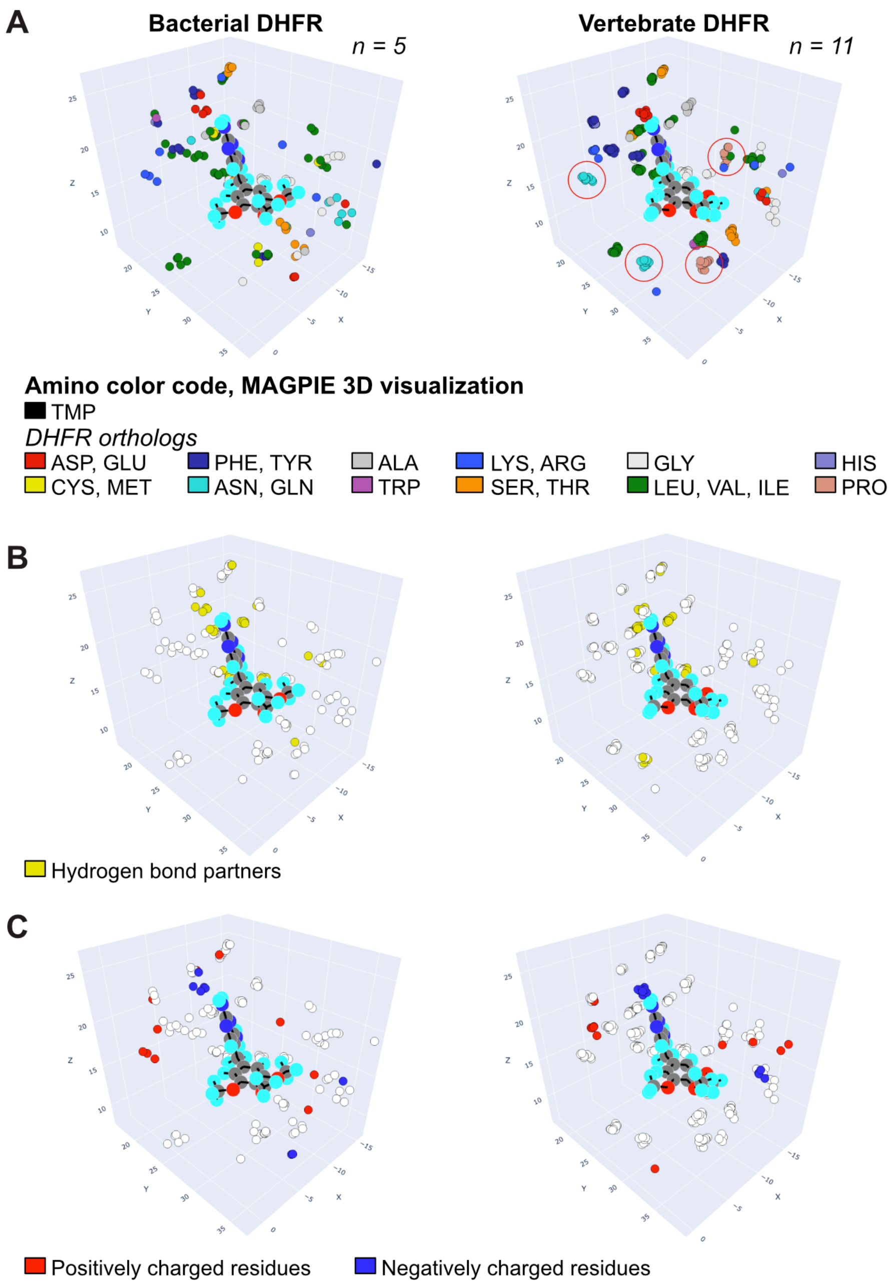

**Figure S20. TMP-bound chemical environment in bacterial and vertebrate DHFR.**

3D visualization of the TMP chemical environment for experimentally solved structures and computationally predicted structural models of five TMP-sensitive bacterial and eleven vertebrate DHFR orthologs (Fig. S21). **A.** Amino acids in the binding pocket, with main differences circled in red, corresponding to a loop that is longer in vertebrate DHFRs to facilitate the closed conformation[12]. **B.** Hydrogen bond donors/acceptors. **C.** Charged residues. Computational models were generated by structural alignment of TMP in an AlphaFold3[13] model with the crystallographic TMP from the most closely evolutionarily related experimentally solved structure available, followed by Rosetta relax[9]. Visualization performed in MAGPIE.[14]

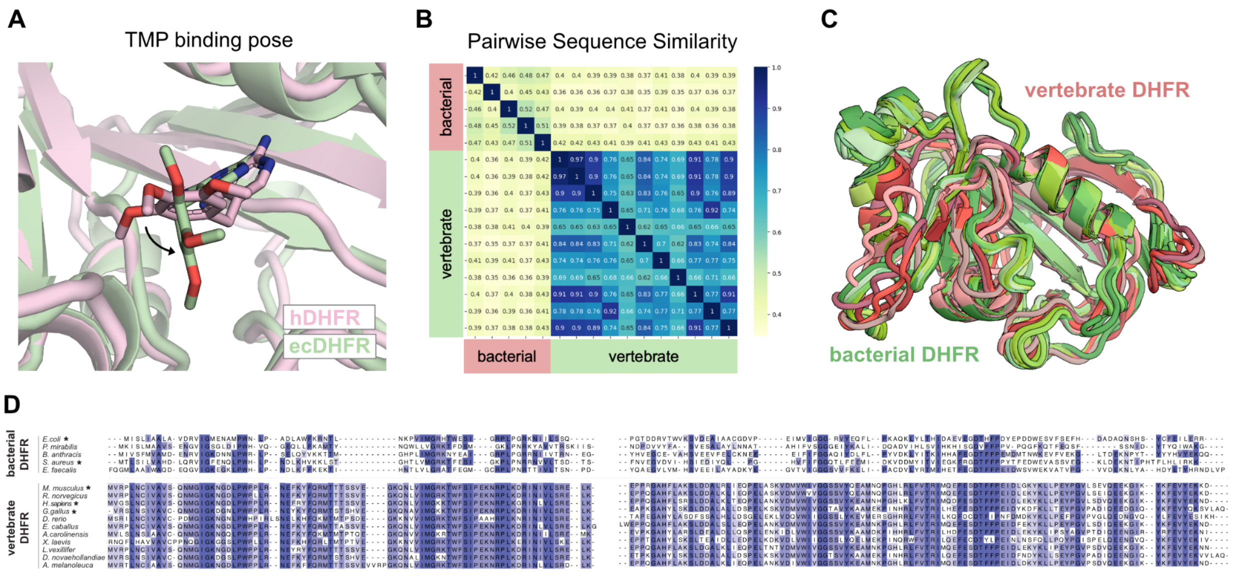

**Figure S21. Sequence/structure comparison for DHFR orthologs.**

**A.** TMP binding pose of hDFR (PDB: 2W3A) and ecDHFR (PDB: 6XG5).[7,15] **B.** Pairwise sequence similarity among DHFR orthologs. **C.** Structure overlap for the same proteins in B., with vertebrate DHFRs shown in red and bacterial DHFRs shown in green. **D.** Multi-sequence alignment. An asterisk (*) indicates that the experimentally solved structure is used in panel C: *E. coli* (PDB: 6XG5), *S. aureus* (PDB: 2W9S), *M. musculus* (PDB: 1U70), *H. sapiens* (PDB: 1BOZ), *G. gallus* (PDB: 1DR1).[16–18] Otherwise, AF3 models are used.[13]

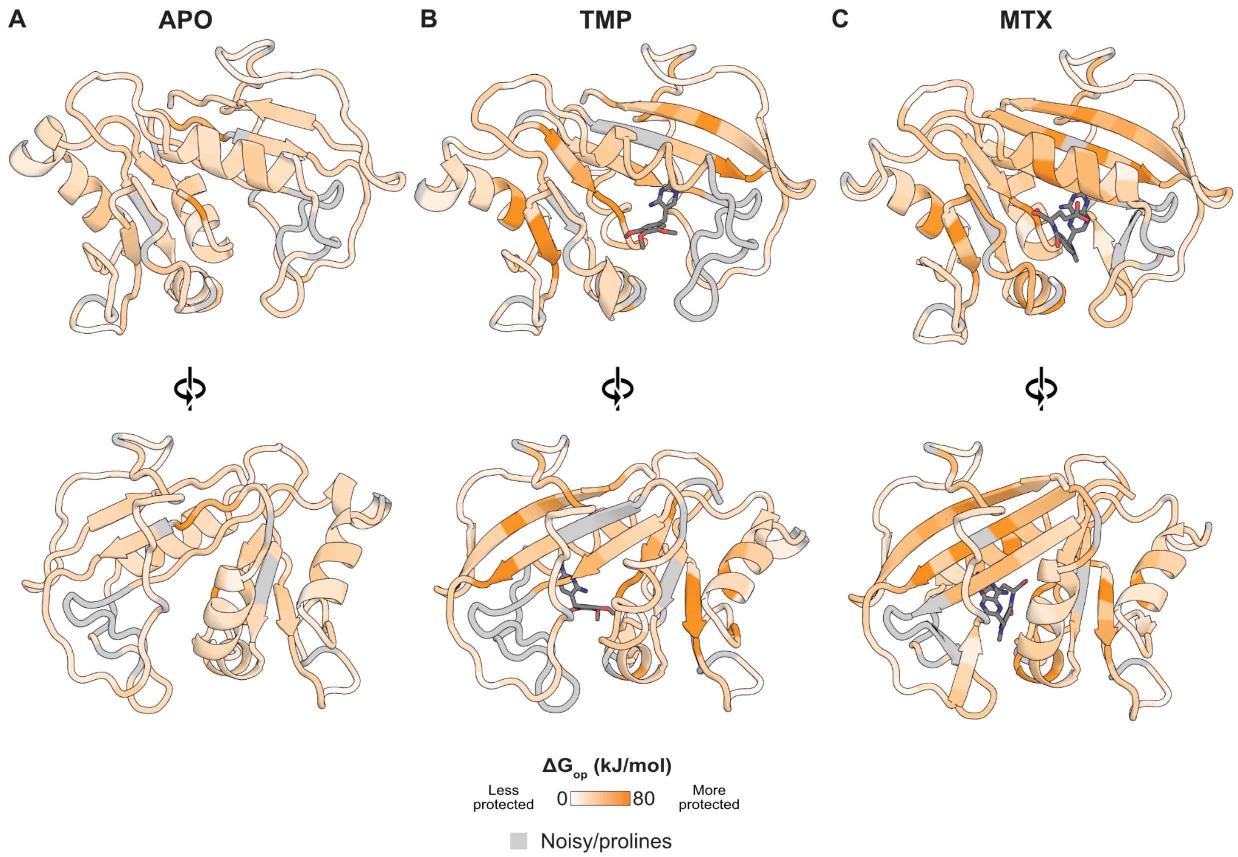

**Figure S22. Absolute $\Delta G_{op}$ for hDHFR functional states.**

**A.** APO hDHFR (PDB ID: 4M6J[19]). **B.** TMP-hDHFR (PDB: 2W3A). **C.** MTX-hDHFR (PDB ID: 1U72[20]).

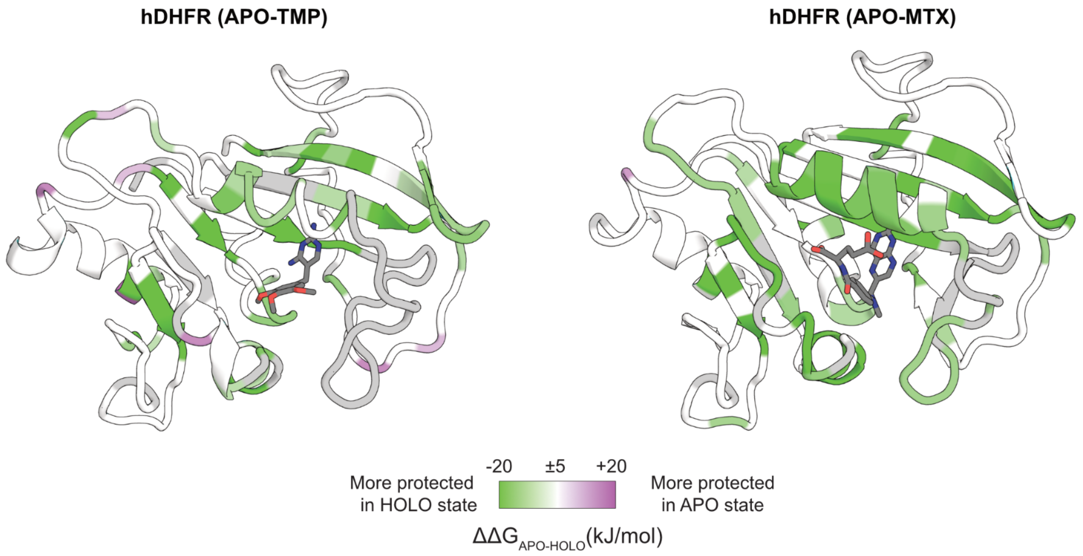

**Figure S23. Comparing inhibitor- vs. APO state hDHFR.**

Noisy/low data regions and prolines are colored gray.

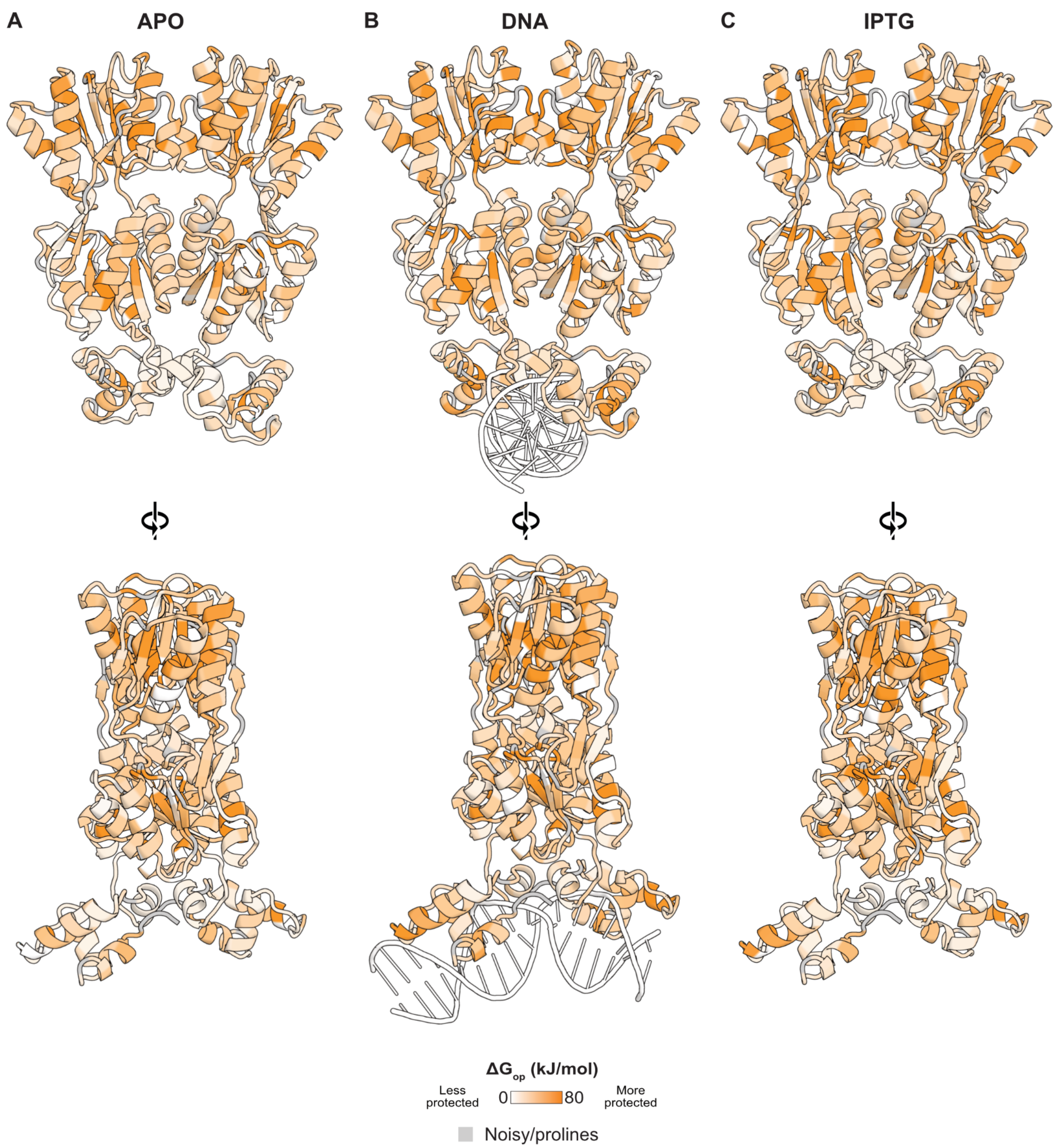

**Figure S24. Absolute $\Delta G_{op}$ for LacI functional states.**

**A.** APO, **B.** DNA-bound, and **C.** IPTG-bound LacI. The DNA-bound state (PDB ID: 1EFA[21]) was used to model all states for ease of comparison because the available APO- and IPTG-LacI X-ray crystal structures do not include a DNA-binding domain.

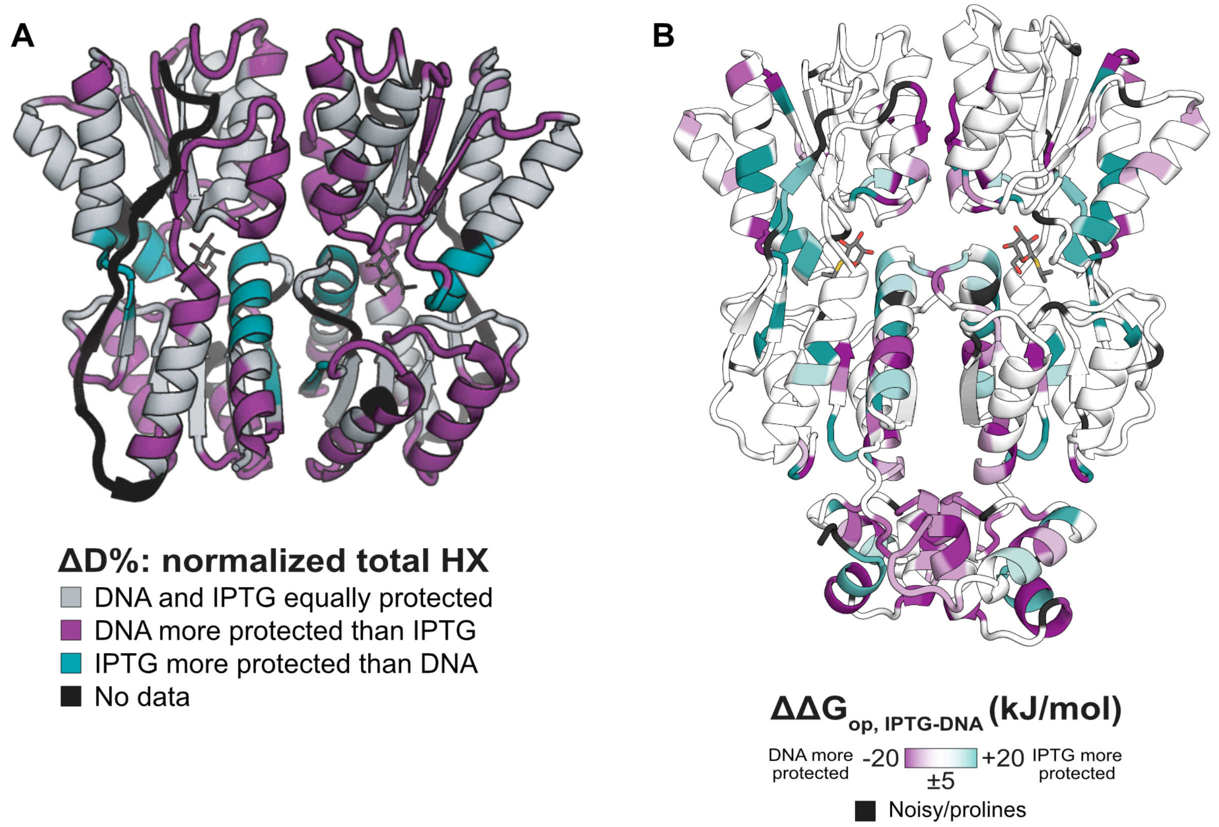

**Figure S25. Comparing LacI HX from Glasgow *et al.*, 2023 to this study.**

**A.** Normalized and back-exchange-corrected D uptake from 57 curated peptides comparing IPTG and DNA states of LacI from Glasgow *et al.*, 2023[22], for all effects larger than 20% at 3+ timepoints in the exchange timecourse. PDB ID: 2P9H[23]. Representations of the same data using more stringent thresholds are available in Fig. S6 of that study. **B.** $\Delta\Delta G_{op,\ IPTG\text{-}DNA}$ from this work, colored similarly to (A) for comparison (for all effects >5 kJ/mol or <-5 kJ/mol). DNA binding domain included. PDB ID: 1EFA.

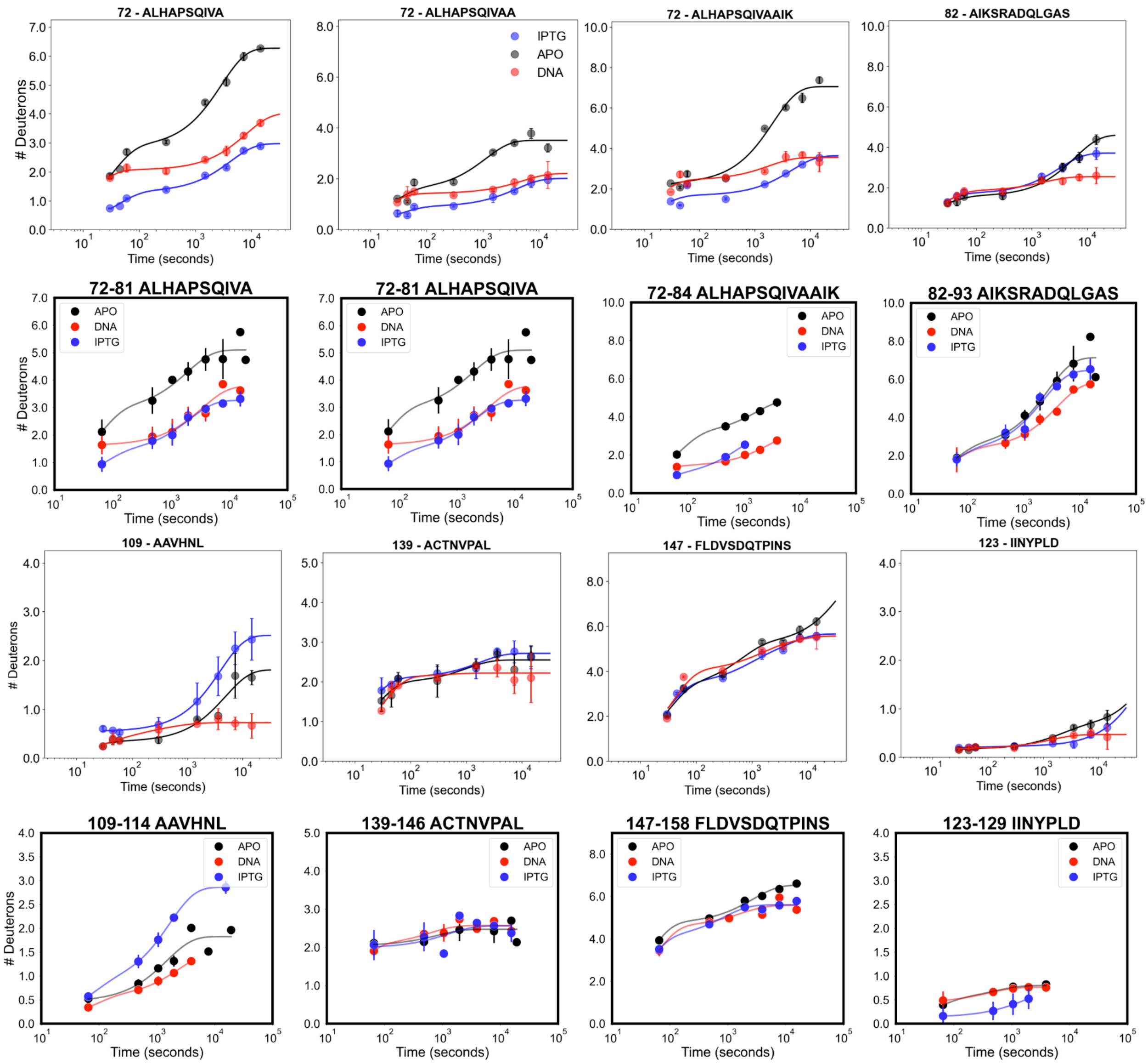

**Figure S26. D uptake plots, this study vs. Glasgow *et al*., 2023.**

Apo, IPTG-, and DNA-LacI show the same trends for each functional state. For each pair of rows, top rows: Glasgow *et al.*[22] and bottom rows: Lu, Wells, *et al.*

Figure continued on the next page.

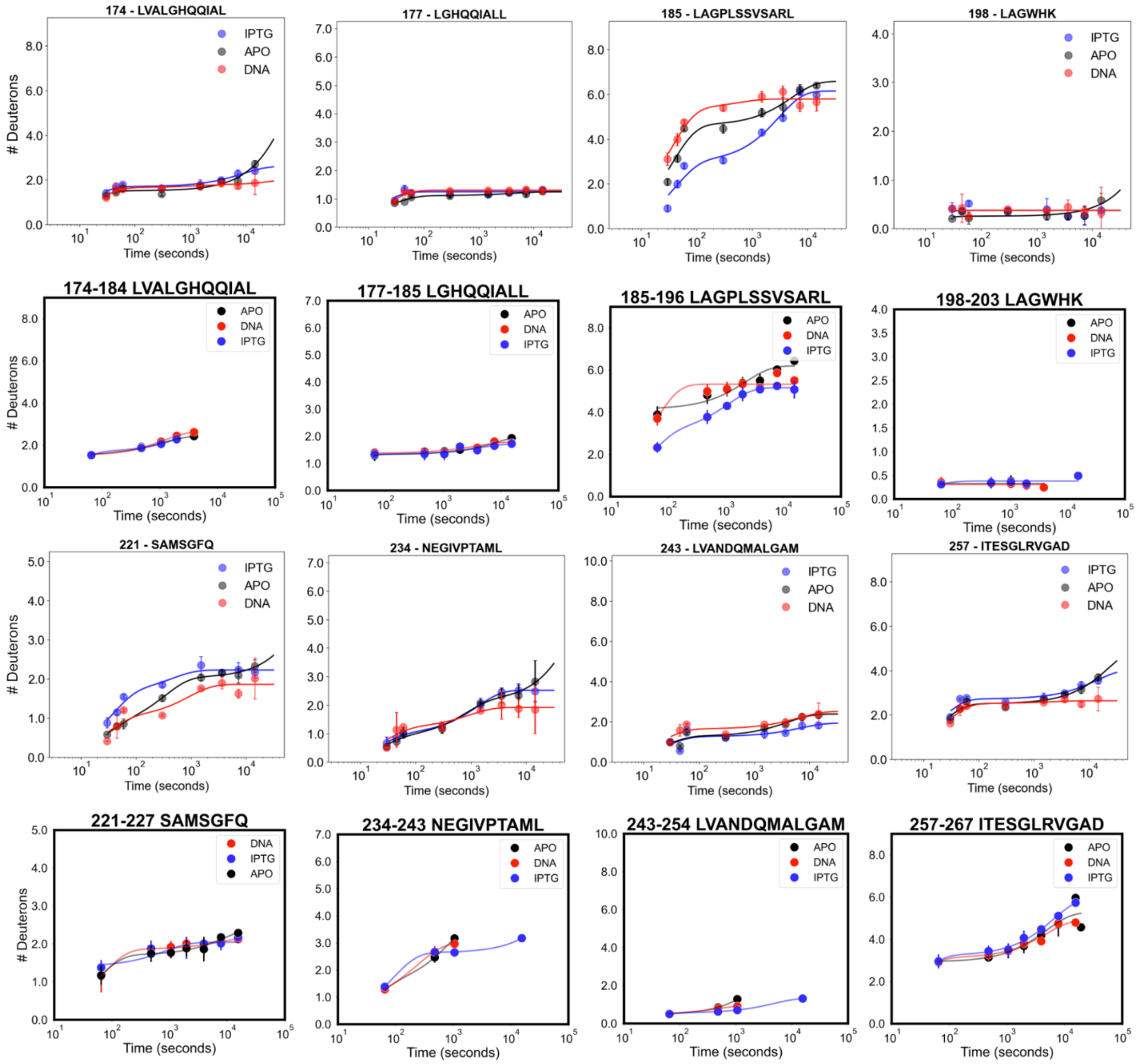

Figure continued on the next page.

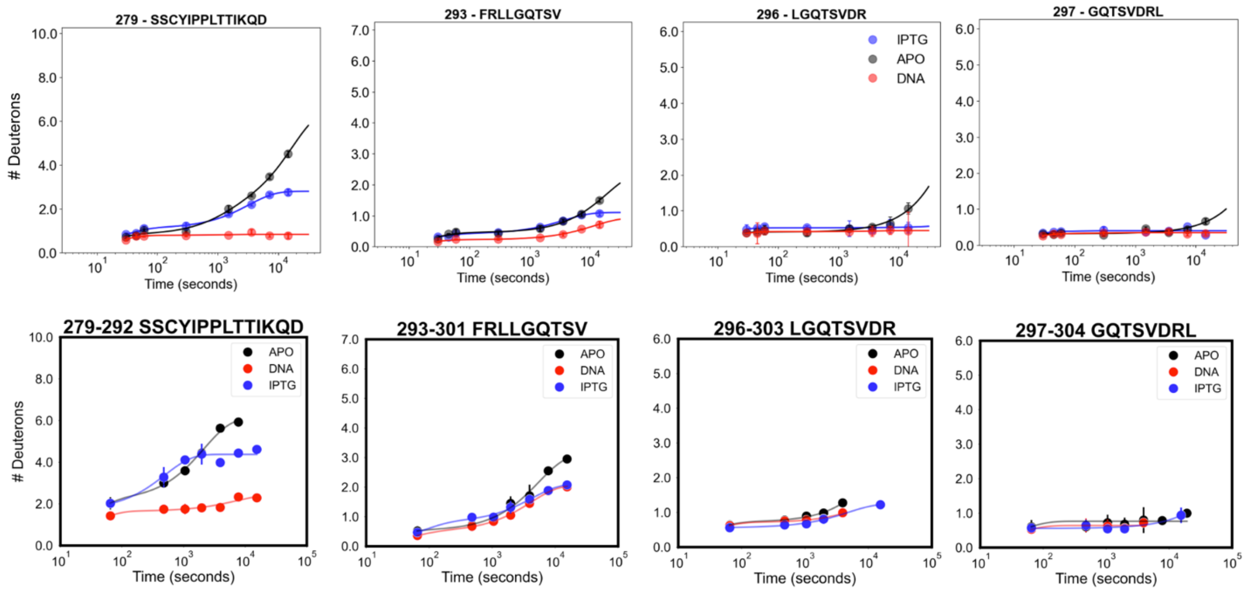

279 - SSCYIPPLTTIKQD
293 - FRLLGQTSV
296 - LGQTSVDR
297 - GQTSVDRL
IPTG
APO
DNA
### Deuterons
Time (seconds)
279-292 SSCYIPPLTTIKQD
293-301 FRLLGQTSV
296-303 LGQTSVDR
297-304 GQTSVDRL

### 4. Supplemental tables

**Table S1. Peptide features throughout PIGEON analysis stages.**

- Analysis stage: the step in PIGEON after which numbers are reported
    - *input*: initial pooled data (Extended Data Fig. 1, Fig. S1);
    - *high score*: matches selected to perform the calibration correction fit;
    - *mz threshold*: peptides remaining after the calibration correction;
    - *single match*: number of consensus matches before disambiguation;
    - *clean*: final set of peptides after disambiguation;
    - *degenerate*: peptides discarded in mass disambiguation (Extended Data Fig. 1e, iii, iv, g).
- N(peps): number of peptides in the set
- %0: percentage of peptides in the set with fragment score 0 (Extended Data Fig. 1d)
- mean score: mean fragment score
- *m*/*z* dev.: RMS distance for all peptides in the set from the heuristic fit line (in ppm)

| HX/MS dataset | Analysis stage | N(peps) | 0% | mean score | m/z dev. |
|---|---|---|---|---|---|
| LacI Rep1 | input | 19804 | 21.5 | 0.009 | 17.7 |
| | high score | 673 | 0.0 | 0.067 | 1.58 |
| | mz threshold | 10866 | 14.9 | 0.0140 | 2.18 |
| | single match | 635 | 10.2 | 0.022 | 2.58 |
| | clean | 439 | 0.5 | 0.031 | 1.84 |
| | degenerate | 196 | 32.1 | 0.002 | 3.73 |
| LacI Rep2 | input | 16144 | 11.2 | 0.013 | 13.5 |
| | high score | 964 | 0.0 | 0.069 | 1.1 |
| | mz threshold | 10896 | 7.2 | 0.018 | 2.0 |
| | single match | 383 | 7.0 | 0.025 | 2.1 |
| | clean | 298 | 0.3 | 0.031 | 1.8 |
| | degenerate | 85 | 30.6 | 0.005 | 3.1 |
| LacI Rep3 | input | 9160 | 21.1 | 0.007 | 12.9 |
| | high score | 104 | 0.0 | 0.062 | 2.5 |
| | mz threshold | 5591 | 15.4 | 0.010 | 2.1 |
| | single match | 493 | 7.1 | 0.016 | 2.2 |
| | clean | 379 | 2.9 | 0.020 | 2.0 |
| | degenerate | 114 | 21.1 | 0.004 | 2.9 |

| | | | | | |
|---|---|---|---|---|---|
| ecDHFR Rep1/Rep2 | input | 2961 | 20.8 | 0.010 | 10.8 |
| | high score | 96 | 0.0 | 0.068 | 0.9 |
| | mz threshold | 2244 | 15.0 | 0.013 | 1.2 |
| | single match | 276 | 11.6 | 0.020 | 1.4 |
| | clean | 232 | 6.0 | 0.023 | 1.2 |
| | degenerate | 44 | 40.9 | 0.006 | 2.2 |
| ecDHFR Rep3 | input | 2704 | 8.5 | 0.011 | 12.1 |
| | high score | 93 | 0.0 | 0.067 | 0.7 |
| | mz threshold | 1884 | 3.6 | 0.015 | 1.1 |
| | single match | 198 | 3.5 | 0.021 | 1.5 |
| | clean | 185 | 3.8 | 0.022 | 1.4 |
| | degenerate | 13 | 0.0 | 0.008 | 3.0 |
| ecDHFR Rep4 | input | 3300 | 21.2 | 0.011 | 13.7 |
| | high score | 129 | 0.0 | 0.067 | 2.6 |
| | mz threshold | 2357 | 13.1 | 0.015 | 2.0 |
| | single match | 258 | 11.2 | 0.021 | 2.4 |
| | clean | 219 | 8.7 | 0.023 | 1.9 |
| | degenerate | 39 | 25.6 | 0.006 | 4.1 |
| ecDHFR Rep5 | input | 3317 | 12.6 | 0.008 | 10.6 |
| | high score | 63 | 0.0 | 0.066 | 0.5 |
| | mz threshold | 2481 | 8.9 | 0.011 | 1.5 |
| | single match | 327 | 5.2 | 0.017 | 1.8 |
| | clean | 274 | 2.6 | 0.019 | 1.5 |
| | degenerate | 53 | 18.9 | 0.005 | 2.7 |
| ecDHFR Rep6 | input | 1324 | 26.4 | 0.008 | 11.1 |
| | high score | 14 | 0.0 | 0.059 | 0.7 |
| | mz threshold | 962 | 19.9 | 0.010 | 1.2 |
| | single match | 182 | 15.4 | 0.016 | 1.3 |
| | clean | 150 | 9.3 | 0.018 | 1.2 |
| | degenerate | 32 | 43.8 | 0.006 | 1.8 |
| ecDHFR Rep7 | input | 2944 | 19.4 | 0.009 | 10.8 |
| | high score | 64 | 0.0 | 0.068 | 0.8 |

| | | | | | |
|---|---|---|---|---|---|
| | mz threshold | 2138 | 14.8 | 0.012 | 1.5 |
| | single match | 242 | 12.0 | 0.018 | 2.0 |
| | clean | 196 | 6.6 | 0.021 | 1.6 |
| | degenerate | 46 | 34.8 | 0.006 | 3.1 |
| hDHFR Rep1 | input | 3072 | 24.2 | 0.003 | 14.9 |
| | high score | 23 | 0.0 | 0.062 | 0.7 |
| | mz threshold | 1315 | 17.1 | 0.007 | 2.5 |
| | single match | 372 | 11.8 | 0.008 | 2.9 |
| | clean | 229 | 2.6 | 0.013 | 2.3 |
| | degenerate | 143 | 26.6 | 0.002 | 3.8 |
| hDHFR Rep2 | input | 3430 | 46.7 | 0.003 | 15.4 |
| | high score | 19 | 0.0 | 0.064 | 0.4 |
| | mz threshold | 1351 | 34.9 | 0.006 | 2.4 |
| | single match | 318 | 32.7 | 0.008 | 2.9 |
| | clean | 167 | 14.4 | 0.014 | 2.4 |
| | degenerate | 151 | 53.0 | 0.001 | 3.5 |
| hDHFR Rep3 | input | 12488 | 26.4 | 0.006 | 16.3 |
| | high score | 1116 | 0.0 | 0.039 | 5.6 |
| | mz threshold | 4218 | 22.6 | 0.010 | 3.9 |
| | single match | 600 | 19.3 | 0.009 | 4.1 |
| | clean | 301 | 5.6 | 0.016 | 4.0 |
| | degenerate | 299 | 33.1 | 0.002 | 4.2 |

**Table S2. Protein features of FEATHER simulated datasets.**

| Protein | # peptides | Sequence length | # simulated peptides | # single resolved |
|---|---|---|---|---|
| Sim100-50P | 50 | 100 | 50 | 32 |
| Sim100-100P | 100 | 100 | 100 | 65 |
| Sim100-150P | 150 | 100 | 150 | 85 |
| Sim200-100P | 100 | 200 | 100 | 69 |
| Sim200-200P | 200 | 200 | 200 | 135 |
| Sim200-300P | 300 | 200 | 300 | 170 |
| Sim300-100P | 100 | 300 | 100 | 66 |
| Sim200-200P | 200 | 300 | 200 | 164 |
| Sim300-300P | 300 | 300 | 300 | 196 |
| Sim300-500P | 500 | 300 | 500 | 267 |
| Sim500-300P | 300 | 500 | 300 | 223 |
| Sim500-500P | 500 | 500 | 500 | 328 |
| SimT4100-50P | 50 | 100 | 100 | 80 |

**Table S3. FEATHER simulated datasets.**

We tested synthetic datasets of varying quality, encompassing diverse log(PF) ranges, numbers of time points, time window ranges, and noise levels.

| Data quality | log(P) range | # timepoints | Time window (s) | Noise level (%) |
|---|---|---|---|---|
| perfect | 2-10 | 20 | 1e1 to 1e12 | 0 |
| sparse | 2-10 | 10 | 1e1 to 1e12 | 0 |
| tp_1e6 | 2-10 | 10 | 1e1 to 1e6 | 0 |
| sparse_tp_1e5 | 2-10 | 5 | 1e1 to 1e5 | 0 |
| noise10 | 2-10 | 20 | 1e1 to 1e12 | 10 |
| noise20 | 2-10 | 20 | 1e1 to 1e12 | 20 |
| noise30 | 2-10 | 20 | 1e1 to 1e12 | 30 |
| SimT4100-50P | 2-6 | 7 | 1e1 to 1e5 | 0 |

**Table S4. The contribution of each feature of FEATHER to its overall performance.**

The improvement is calculated by the increase in the R value and decrease in the RMSE compared to the original BayesianHDX method.[24] The benchmark sets tested in the Protein column are detailed in Table S2.

| **Protein** | Overall improvement | Peptide subtraction | Isotopic mass envelope | Swapping mechanism |
|---|---|---|---|---|
| Sim100-50P | **0.20** | **0.03** | **0.02** | **0.07** |
| Sim100-100P | **0.44** | **0.05** | **0.09** | **0.10** |
| Sim100-150P | **0.56** | **0.02** | **0.07** | **0.12** |
| Sim200-100P | **0.26** | **0.02** | **0.02** | **0.06** |
| Sim200-200P | **0.27** | **0.01** | **0.03** | **0.07** |
| Sim200-300P | **0.30** | **-0.01** | **0.08** | **0.10** |
| Sim300-100P | **0.15** | **-0.01** | **0.02** | **0.02** |
| Sim300-200P | **0.27** | **0.02** | **0.06** | **0.07** |
| Sim300-300P | **0.34** | **0.02** | **0.02** | **0.09** |
| Sim300-500P | **0.42** | **0.04** | **0.05** | **0.10** |
| Sim500-300P | **0.28** | **-0.02** | **0.03** | **0.04** |
| Sim500-500P | **0.25** | **0.02** | **0.05** | **0.06** |
| Average | **0.31** | **0.02** | **0.05** | **0.08** |

**Table S5. Benchmarks on simulated datasets with varying data quality.**

The benchmark sets tested in the Protein column are detailed in Table S2. The bolded first number in each column indicates the R correlation value. The second number is the RMSE.

| Protein | perfect | sparse | tp_1e6 | sparse_tp_1e5 | noise10 | noise20 | noise30 |
|---|---|---|---|---|---|---|---|
| Sim100-50P | **0.99**/0.49 | **0.99**/0.51 | **0.95**/1.25 | **0.94/**1.28 | **0.99**/0.57 | **0.99**/0.45 | **0.98**/0.63 |
| Sim100-100P | **0.99**/0.51 | **1.00**/0.39 | **0.92**/1.35 | **0.96**/1.35 | **0.95**/0.87 | **0.99**/0.53 | **0.97**/0.68 |
| Sim100-150P | **1.00**/0.39 | **0.99**/0.48 | **0.97**/0.95 | **0.95**/1.00 | **0.99**/0.49 | **0.99**/0.46 | **1.00**/0.40 |
| Sim200-100P | **0.96**/0.79 | **0.97**/0.73 | **0.95**/1.14 | **0.87**/1.73 | **0.93**/0.98 | **0.93**/1.02 | **0.96**/0.80 |
| Sim200-200P | **0.98**/0.57 | **0.99**/0.48 | **0.96**/0.95 | **0.95**/1.52 | **0.99**/0.51 | **0.97**/0.69 | **0.96**/0.83 |
| Sim200-300P | **0.98**/0.66 | **0.97**/0.72 | **0.94**/1.06 | **0.94**/1.18 | **0.97**/0.68 | **0.95**/0.84 | **0.96**/0.76 |
| Sim300-100P | **0.89**/1.24 | **0.91**/1.15 | **0.85**/1.76 | **0.84**/2.05 | **0.88**/1.29 | **0.90**/1.19 | **0.88**/1.29 |
| Sim300-200P | **0.92**/1.04 | **0.91**/1.11 | **0.92**/1.28 | **0.90**/1.43 | **0.95**/0.90 | **0.93**/1.02 | **0.92**/1.05 |
| Sim300-300P | **0.97**/0.75 | **0.97**/0.70 | **0.95**/1.13 | **0.93**/1.23 | **0.99**/0.53 | **0.97**/0.77 | **0.94**/0.91 |
| Sim300-500P | **1.00**/0.40 | **0.99**/0.51 | **0.97**/0.82 | **0.95**/1.05 | **0.99**/0.50 | **0.99**/0.54 | **0.99**/0.53 |
| Sim500-300P | **0.91**/1.10 | **0.92**/1.02 | **0.84**/1.66 | **0.84**/1.85 | **0.91**/1.13 | **0.90**/1.15 | **0.91**/1.10 |
| Sim500-500P | **0.96**/0.78 | **0.98**/0.64 | **0.91**/1.27 | **0.91**/1.39 | **0.96**/0.78 | **0.95**/0.87 | **0.96**/0.76 |

Representative example of D uptake plots for one peptide from the benchmark datasets

**perfect** **sparse** **tp_1e6** **sparse_tp_1e5** **noise10** **noise20** **noise30**

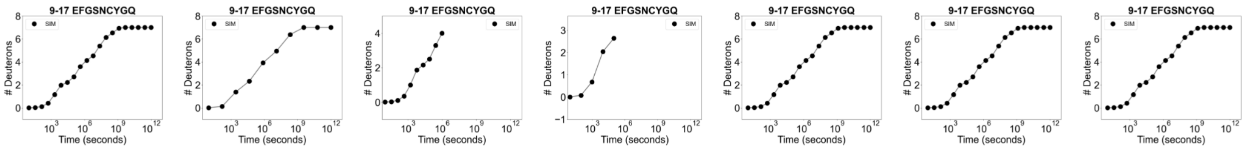

**Table S6. Benchmarks comparison to other methods using the perfect simulated dataset.**

The benchmark sets tested in the Protein column are detailed in Table S2. The bolded first number in each column indicates the R correlation value. The second number is the RMSE. HDSite could not be run on any larger datasets due to the computational cost.

| Protein/Model | FEATHER | BayesianHDX | PyHDX | ExPfact | HDSite |
|---|---|---|---|---|---|
| SimT4100-50P | **0.98**/0.45 | **0.70**/1.24 | **0.71**/0.97 | **0.18**/3.95 | **0.84**/0.69 |
| Sim100-50P | **0.99**/0.49 | **0.79**/1.72 | **0.56**/2.33 | **0.35**/4.65 | - |
| Sim100-100P | **0.99**/0.51 | **0.55**/2.48 | **0.45**/2.65 | **0.34**/4.58 | - |
| Sim100-150P | **1.00**/0.39 | **0.44**/2.75 | **0.56**/2.37 | **0.31**/4.74 | - |
| Sim200-100P | **0.96**/0.79 | **0.70**/1.97 | **0.56**/2.38 | **0.29**/4.74 | - |
| Sim200-200P | **0.98**/0.57 | **0.71**/1.96 | **0.41**/2.65 | **0.25**/4.86 | - |
| Sim200-300P | **0.98**/0.66 | **0.68**/2.03 | **0.47**/2.67 | **0.31**/4.95 | - |
| Sim300-100P | **0.89**/1.24 | **0.74**/1.92 | **0.57**/2.24 | **0.36**/4.85 | - |
| Sim200-200P | **0.92**/1.04 | **0.65**/2.17 | **0.46**/2.57 | **0.28**/4.91 | - |
| Sim300-300P | **0.97**/0.75 | **0.63**/2.23 | **0.39**/2.70 | **0.36**/4.84 | - |
| Sim300-500P | **1.00**/0.40 | **0.58**/2.37 | **0.40**/2.70 | **0.37**/4.85 | - |
| Sim500-300P | **0.91**/1.10 | **0.63**/2.22 | **0.44**/2.49 | **0.33**/4.71 | - |
| Sim500-500P | **0.96**/0.78 | **0.71**/1.93 | **0.37**/2.63 | **0.28**/4.70 | - |

**Table S7. HX experiment summary table for ecDHFR.**

Tables S7, S9, and S11 include all experimental reporting suggestions included in Masson *et al.*[25] All experiments run with KEEP setting in PIGEON. Abbreviations: FP, immobilized protease type XIII. Nep, nepenthesin 2. AP, alanyl aminopeptidase.

| Dataset | **DHFR Rep1** | **DHFR Rep2** | **DHFR Rep3** | **DHFR Rep4** | **DHFR Rep5** | **DHFR Rep6** | **DHFR Rep7** |
|---|---|---|---|---|---|---|---|
| Protein states | APO/TMP | APO/TMP | APO/TMP | APO/TMP | APO/MTX | MTX | APO/MTX |
| Date | 01/12/24 | 01/12/24 | 02/26/24 | 03/04/24 | 06/13/24 | 06/20/24 | 06/20/24 |
| Protease column | FP/pepsin | FP/pepsin | Nep/ pepsin | AP/pepsin | FP/pepsin | Nep/ pepsin | AP/pepsin |
| Reaction details | 50 mM potassium phosphate, 1 mM NaCl, 10 mM BME, pH 7.0, temperature 20 °C, final $D_2O$ concentration 90% | | | | | | |
| Time course (s) | 4.6e1-4.3e4 | 4.6e1-4.3e4 | 4.6e1-2.9e4 | 4.6e1-2.9e4 | 4.3e1-4.0e4 | 4.3e1-1.8e4 | 4.2e1-5.4e4 |
| # of time points | 7 | 7 | 6 | 6 | 5 | 5 | 6 |
| Control samples | Maximally-labeled sample (full-D) | | | | | | |
| Back-exchange (mean / IQR) | 0.23/0.12 | 0.23/0.12 | 0.23/0.11 | 0.23/0.08 | 0.23/0.11 | 0.22/0.10 | 0.22/0.09 |
| # of peptides | 140 | 144 | 115 | 149 | 164 | 107 | 174 |
| Sequence coverage | 0.90 | 0.90 | 0.78 | 0.86 | 0.85 | 0.80 | 0.86 |
| Average peptide length / Redundancy | 1.3 | 1.2 | 1.5 | 1.2 | 1.1 | 1.6 | 1.3 |
| Significant difference in $\Delta\ log\ PF$ | $\Delta\ log\ PF > \frac{\sum_{states} std(log\ PF)}{\sqrt{n_{bootstrap}}}$ | | | | | | |

**Table S8. $\Delta G_{op}$ values for ecDHFR.**

| Res. name | Res. number | Apo | Std Dev | Single resolved | TMP | Std Dev | Single resolved | MTX | Std Dev | Single resolved |
|---|---|---|---|---|---|---|---|---|---|---|
| MET | 1 | 24 | 0 | TRUE | 19 | 3 | TRUE | 23 | 2 | TRUE |
| ILE | 2 | 26 | 0 | FALSE | 48 | 5 | FALSE | 70 | 2 | FALSE |
| SER | 3 | 30 | 0 | FALSE | 51 | 5 | FALSE | 73 | 2 | FALSE |
| LEU | 4 | 12 | 1 | TRUE | 23 | 9 | TRUE | 15 | 3 | TRUE |
| ILE | 5 | 21 | 0 | TRUE | 23 | 1 | TRUE | 25 | 0 | TRUE |
| ALA | 6 | | | | | | | 16 | 1 | TRUE |
| ALA | 7 | 19 | 0 | TRUE | 25 | 1 | TRUE | 29 | 7 | TRUE |
| LEU | 8 | 18 | 0 | FALSE | 23 | 1 | FALSE | 29 | 7 | TRUE |
| ALA | 9 | 21 | 0 | FALSE | 26 | 1 | FALSE | 31 | 1 | TRUE |
| VAL | 10 | 15 | 0 | TRUE | 21 | 1 | TRUE | 20 | 0 | TRUE |
| ASP | 11 | 14 | 1 | TRUE | 15 | 0 | TRUE | 15 | 0 | TRUE |
| ARG | 12 | 19 | 1 | TRUE | 22 | 1 | TRUE | 23 | 0 | TRUE |
| VAL | 13 | 15 | 1 | TRUE | 22 | 1 | TRUE | 24 | 0 | TRUE |
| ILE | 14 | 11 | 1 | TRUE | 16 | 1 | TRUE | 17 | 0 | TRUE |
| GLY | 15 | 11 | 1 | TRUE | 17 | 7 | TRUE | 13 | 1 | TRUE |
| MET | 16 | 24 | 2 | TRUE | 24 | 6 | TRUE | 31 | 0 | TRUE |
| GLU | 17 | 9 | 4 | TRUE | 11 | 2 | TRUE | 14 | 1 | TRUE |
| ASN | 18 | 7 | 3 | TRUE | 10 | 5 | TRUE | 16 | 1 | TRUE |
| ALA | 19 | 20 | 2 | TRUE | 22 | 1 | TRUE | 29 | 1 | TRUE |
| MET | 20 | 12 | 1 | TRUE | 8 | 4 | TRUE | 12 | 1 | TRUE |
| PRO | 21 | | | | | | | | | |
| TRP | 22 | 13 | 1 | TRUE | 15 | 3 | TRUE | 16 | 1 | TRUE |
| ASN | 23 | 10 | 6 | TRUE | 18 | 3 | TRUE | 10 | 7 | TRUE |
| LEU | 24 | 12 | 1 | TRUE | 17 | 1 | TRUE | 13 | 0 | TRUE |
| PRO | 25 | | | | | | | | | |
| ALA | 26 | 18 | 8 | TRUE | 19 | 2 | FALSE | 32 | 1 | TRUE |
| ASP | 27 | 14 | 9 | TRUE | 21 | 2 | FALSE | 13 | 0 | TRUE |
| LEU | 28 | 13 | 1 | TRUE | 24 | 5 | TRUE | 28 | 1 | TRUE |

| | | | | | | | | | | |
|---|---|---|---|---|---|---|---|---|---|---|
| ALA | 29 | 15 | 2 | TRUE | 21 | 2 | TRUE | 24 | 0 | TRUE |
| TRP | 30 | 16 | 1 | TRUE | 22 | 1 | TRUE | 32 | 0 | TRUE |
| PHE | 31 | 18 | 1 | TRUE | 26 | 1 | TRUE | 36 | 4 | TRUE |
| LYS | 32 | 21 | 1 | TRUE | 27 | 1 | TRUE | 34 | 2 | TRUE |
| ARG | 33 | 16 | 0 | TRUE | 23 | 0 | TRUE | 33 | 1 | TRUE |
| ASN | 34 | 20 | 0 | TRUE | 27 | 0 | TRUE | 39 | 2 | TRUE |
| THR | 35 | 27 | 0 | TRUE | 32 | 0 | TRUE | 32 | 1 | TRUE |
| LEU | 36 | 16 | 1 | TRUE | 17 | 3 | TRUE | 35 | 5 | TRUE |
| ASP | 37 | 15 | 0 | TRUE | 16 | 3 | TRUE | 23 | 1 | TRUE |
| LYS | 38 | 31 | 0 | TRUE | 72 | 5 | TRUE | 72 | 5 | TRUE |
| PRO | 39 | | | | | | | | | |
| VAL | 40 | 69 | 7 | TRUE | 68 | 2 | FALSE | 66 | 8 | TRUE |
| ILE | 41 | 69 | 7 | TRUE | 69 | 2 | FALSE | 46 | 9 | FALSE |
| MET | 42 | 71 | 5 | TRUE | 71 | 6 | TRUE | 50 | 9 | FALSE |
| GLY | 43 | 23 | 1 | FALSE | 43 | 5 | FALSE | 28 | 5 | FALSE |
| ARG | 44 | 24 | 1 | FALSE | 44 | 5 | FALSE | 29 | 5 | FALSE |
| HIS | 45 | 24 | 1 | TRUE | 25 | 6 | TRUE | 33 | 2 | TRUE |
| THR | 46 | 58 | 22 | TRUE | 35 | 1 | TRUE | 30 | 3 | TRUE |
| TRP | 47 | 22 | 5 | TRUE | 32 | 7 | TRUE | 35 | 2 | TRUE |
| GLU | 48 | 20 | 4 | TRUE | 21 | 1 | TRUE | 30 | 1 | TRUE |
| SER | 49 | 17 | 1 | TRUE | 20 | 0 | TRUE | 70 | 6 | TRUE |
| ILE | 50 | 17 | 1 | TRUE | 19 | 0 | TRUE | 35 | 3 | TRUE |
| GLY | 51 | 10 | 2 | TRUE | 13 | 1 | TRUE | 16 | 0 | TRUE |
| ARG | 52 | 17 | 1 | TRUE | 15 | 1 | TRUE | 30 | 1 | TRUE |
| PRO | 53 | | | | | | | | | |
| LEU | 54 | 10 | 1 | FALSE | 10 | 1 | FALSE | 22 | 1 | FALSE |
| PRO | 55 | | | | | | | | | |
| GLY | 56 | 12 | 2 | TRUE | 15 | 3 | TRUE | 16 | 2 | FALSE |
| ARG | 57 | 14 | 2 | TRUE | 8 | 1 | FALSE | 19 | 2 | FALSE |
| LYS | 58 | 10 | 4 | TRUE | 8 | 1 | FALSE | 12 | 3 | TRUE |
| ASN | 59 | 34 | 2 | TRUE | 24 | 13 | TRUE | 41 | 3 | TRUE |

| | | | | | | | | | | |
|---|---|---|---|---|---|---|---|---|---|---|
| ILE | 60 | 52 | 27 | TRUE | 71 | 2 | FALSE | 67 | 7 | TRUE |
| ILE | 61 | 54 | 21 | TRUE | 68 | 2 | FALSE | 66 | 8 | TRUE |
| LEU | 62 | 62 | 16 | TRUE | 71 | 6 | TRUE | 33 | 1 | TRUE |
| SER | 63 | 8 | 5 | TRUE | 14 | 3 | TRUE | 12 | 7 | TRUE |
| SER | 64 | 34 | 1 | TRUE | 26 | 13 | TRUE | 17 | 9 | TRUE |
| GLN | 65 | 19 | 6 | TRUE | 19 | 9 | TRUE | 30 | 7 | TRUE |
| PRO | 66 | | | | | | | | | |
| GLY | 67 | 9 | 3 | FALSE | 8 | 3 | FALSE | 5 | 1 | FALSE |
| THR | 68 | 6 | 3 | TRUE | 5 | 2 | TRUE | 5 | 2 | TRUE |
| ASP | 69 | 14 | 9 | TRUE | 18 | 11 | TRUE | 23 | 7 | TRUE |
| ASP | 70 | 20 | 6 | TRUE | 16 | 9 | TRUE | 14 | 9 | TRUE |
| ARG | 71 | 12 | 0 | TRUE | 12 | 2 | TRUE | 13 | 1 | TRUE |
| VAL | 72 | 19 | 1 | TRUE | 20 | 1 | TRUE | 21 | 1 | TRUE |
| THR | 73 | 23 | 2 | TRUE | 25 | 1 | TRUE | 25 | 1 | TRUE |
| TRP | 74 | 55 | 19 | TRUE | 28 | 8 | TRUE | 45 | 22 | TRUE |
| VAL | 75 | 15 | 5 | TRUE | 30 | 8 | TRUE | 16 | 1 | FALSE |
| LYS | 76 | 18 | 5 | TRUE | 13 | 1 | TRUE | 20 | 1 | FALSE |
| SER | 77 | 26 | 1 | TRUE | 25 | 2 | TRUE | 33 | 0 | TRUE |
| VAL | 78 | 10 | 2 | TRUE | 6 | 3 | TRUE | 9 | 4 | TRUE |
| ASP | 79 | 12 | 0 | TRUE | 13 | 6 | TRUE | 11 | 5 | TRUE |
| GLU | 80 | 18 | 1 | TRUE | 15 | 4 | TRUE | 19 | 0 | TRUE |
| ALA | 81 | 73 | 4 | TRUE | 70 | 5 | TRUE | 69 | 6 | TRUE |
| ILE | 82 | 29 | 1 | TRUE | 43 | 17 | TRUE | 28 | 1 | TRUE |
| ALA | 83 | 29 | 1 | TRUE | 28 | 1 | TRUE | 27 | 1 | TRUE |
| ALA | 84 | 29 | 2 | TRUE | 33 | 4 | TRUE | 34 | 3 | TRUE |
| CYS | 85 | 32 | 2 | TRUE | 44 | 18 | TRUE | 37 | 3 | TRUE |
| GLY | 86 | 15 | 0 | TRUE | 15 | 1 | TRUE | 17 | 0 | TRUE |
| ASP | 87 | 33 | 1 | TRUE | 32 | 1 | TRUE | 26 | 6 | TRUE |
| VAL | 88 | 16 | 1 | TRUE | 15 | 1 | TRUE | 21 | 2 | TRUE |
| PRO | 89 | | | | | | | | | |
| GLU | 90 | 20 | 1 | TRUE | 45 | 5 | FALSE | 34 | 24 | TRUE |

| | | | | | | | | | | |
|---|---|---|---|---|---|---|---|---|---|---|
| ILE | 91 | 66 | 7 | TRUE | 42 | 5 | FALSE | 68 | 7 | TRUE |
| MET | 92 | 51 | 17 | TRUE | 59 | 22 | TRUE | 57 | 21 | TRUE |
| VAL | 93 | 33 | 3 | TRUE | 71 | 6 | TRUE | 27 | 1 | TRUE |
| ILE | 94 | 69 | 6 | TRUE | 70 | 6 | TRUE | 65 | 8 | TRUE |
| GLY | 95 | 25 | 1 | TRUE | 20 | 6 | TRUE | 15 | 1 | TRUE |
| GLY | 96 | 11 | 3 | TRUE | 18 | 6 | TRUE | 38 | 1 | TRUE |
| GLY | 97 | 17 | 1 | TRUE | 13 | 2 | TRUE | 17 | 2 | FALSE |
| ARG | 98 | 13 | 3 | TRUE | 14 | 1 | TRUE | 18 | 2 | FALSE |
| VAL | 99 | 12 | 2 | TRUE | 23 | 5 | TRUE | 16 | 3 | TRUE |
| TYR | 100 | 19 | 0 | TRUE | 17 | 1 | TRUE | 23 | 1 | TRUE |
| GLU | 101 | 20 | 0 | TRUE | 20 | 0 | TRUE | 23 | 0 | TRUE |
| GLN | 102 | 23 | 0 | TRUE | 27 | 0 | TRUE | 27 | 0 | TRUE |
| PHE | 103 | 28 | 1 | TRUE | 35 | 1 | TRUE | 47 | 16 | TRUE |
| LEU | 104 | 26 | 1 | TRUE | 33 | 1 | TRUE | 34 | 1 | TRUE |
| PRO | 105 | | | | | | | | | |
| LYS | 106 | 25 | 1 | TRUE | 32 | 0 | TRUE | 20 | 0 | TRUE |
| ALA | 107 | 19 | 1 | TRUE | 20 | 0 | TRUE | 33 | 1 | FALSE |
| GLN | 108 | 25 | 0 | FALSE | 32 | 4 | FALSE | 33 | 1 | FALSE |
| LYS | 109 | 26 | 0 | FALSE | 32 | 4 | FALSE | 33 | 1 | FALSE |
| LEU | 110 | 24 | 1 | TRUE | 30 | 1 | TRUE | 34 | 2 | TRUE |
| TYR | 111 | 23 | 0 | TRUE | 28 | 0 | TRUE | 43 | 18 | TRUE |
| LEU | 112 | 25 | 1 | TRUE | 62 | 11 | TRUE | 69 | 6 | TRUE |
| THR | 113 | 31 | 2 | TRUE | 37 | 4 | TRUE | 34 | 18 | TRUE |
| HIS | 114 | 32 | 2 | TRUE | 38 | 1 | TRUE | 51 | 17 | TRUE |
| ILE | 115 | 18 | 1 | FALSE | 25 | 0 | FALSE | 46 | 5 | FALSE |
| ASP | 116 | 18 | 1 | FALSE | 25 | 0 | FALSE | 47 | 5 | FALSE |
| ALA | 117 | 18 | 1 | TRUE | 21 | 0 | TRUE | 30 | 1 | TRUE |
| GLU | 118 | 6 | 3 | TRUE | 5 | 2 | TRUE | 10 | 1 | FALSE |
| VAL | 119 | 7 | 1 | FALSE | 10 | 2 | FALSE | 6 | 1 | FALSE |
| GLU | 120 | 11 | 1 | FALSE | 14 | 2 | FALSE | 9 | 1 | FALSE |
| GLY | 121 | 17 | 1 | FALSE | 13 | 1 | FALSE | | | |

| | | | | | | | | | | |
|---|---|---|---|---|---|---|---|---|---|---|
| ASP | 122 | 20 | 1 | FALSE | 16 | 1 | FALSE | | | |
| THR | 123 | 17 | 1 | FALSE | 13 | 1 | FALSE | | | |
| HIS | 124 | 32 | 26 | TRUE | 19 | 15 | TRUE | | | |
| PHE | 125 | 21 | 1 | TRUE | 23 | 3 | TRUE | | | |
| PRO | 126 | | | | | | | | | |
| ASP | 127 | 37 | 6 | FALSE | 44 | 6 | FALSE | | | |
| TYR | 128 | 35 | 6 | FALSE | 42 | 6 | FALSE | | | |
| GLU | 129 | 16 | 1 | TRUE | 15 | 0 | TRUE | 13 | 1 | TRUE |
| PRO | 130 | | | | | | | | | |
| ASP | 131 | 13 | 1 | TRUE | 12 | 1 | TRUE | 54 | 14 | TRUE |
| ASP | 132 | 33 | 4 | TRUE | 44 | 16 | TRUE | 23 | 0 | TRUE |
| TRP | 133 | 25 | 3 | TRUE | 28 | 2 | FALSE | 46 | 23 | TRUE |
| GLU | 134 | 26 | 2 | TRUE | 30 | 2 | FALSE | 29 | 11 | TRUE |
| SER | 135 | 24 | 6 | TRUE | 22 | 7 | TRUE | 22 | 1 | FALSE |
| VAL | 136 | 17 | 6 | TRUE | 25 | 1 | TRUE | 19 | 1 | FALSE |
| PHE | 137 | 24 | 0 | TRUE | 21 | 6 | TRUE | 25 | 0 | TRUE |
| SER | 138 | 27 | 8 | TRUE | 17 | 12 | TRUE | | | |
| GLU | 139 | 24 | 2 | TRUE | 33 | 1 | TRUE | 34 | 1 | TRUE |
| PHE | 140 | 4 | 2 | TRUE | 15 | 6 | TRUE | 3 | 2 | TRUE |
| HIS | 141 | 13 | 6 | TRUE | 16 | 5 | TRUE | 26 | 7 | TRUE |
| ASP | 142 | 21 | 0 | TRUE | 17 | 1 | TRUE | 73 | 4 | TRUE |
| ALA | 143 | 71 | 6 | TRUE | 30 | 1 | TRUE | 71 | 5 | TRUE |
| ASP | 144 | 45 | 22 | TRUE | 16 | 0 | TRUE | 45 | 18 | TRUE |
| ALA | 145 | 13 | 4 | TRUE | 8 | 4 | TRUE | 11 | 1 | TRUE |
| GLN | 146 | 8 | 5 | TRUE | 10 | 4 | TRUE | 7 | 4 | TRUE |
| ASN | 147 | 8 | 4 | TRUE | 18 | 1 | TRUE | 72 | 5 | TRUE |
| SER | 148 | 23 | 3 | TRUE | 23 | 0 | TRUE | 19 | 0 | TRUE |
| HIS | 149 | 21 | 1 | TRUE | 16 | 2 | TRUE | 34 | 6 | TRUE |
| SER | 150 | 25 | 2 | TRUE | 34 | 1 | TRUE | 34 | 9 | TRUE |
| TYR | 151 | 45 | 19 | TRUE | 34 | 3 | TRUE | 70 | 6 | TRUE |
| CYS | 152 | 31 | 1 | TRUE | 35 | 1 | TRUE | 72 | 4 | TRUE |

| | | | | | | | | | | |
|---|---|---|---|---|---|---|---|---|---|---|
| PHE | 153 | 70 | 5 | TRUE | 35 | 4 | TRUE | 73 | 4 | TRUE |
| GLU | 154 | 32 | 1 | TRUE | 42 | 19 | TRUE | 36 | 3 | TRUE |
| ILE | 155 | 27 | 0 | TRUE | 47 | 24 | TRUE | 31 | 0 | TRUE |
| LEU | 156 | 24 | 1 | TRUE | 30 | 1 | TRUE | 31 | 1 | TRUE |
| GLU | 157 | 27 | 1 | TRUE | 34 | 1 | TRUE | 44 | 20 | TRUE |
| ARG | 158 | 24 | 1 | FALSE | 26 | 0 | FALSE | 42 | 19 | TRUE |
| ARG | 159 | 15 | 1 | FALSE | 18 | 0 | FALSE | 15 | 0 | TRUE |

**Table S9. HX experiment summary table for hDHFR.**

All experiments run with KEEP setting in PIGEON. Abbreviations: FP, immobilized protease type XIII. Nep, nepenthesin 2. AP, alanyl aminopeptidase.

| Dataset | **hDHFR Rep1** | **hDHFR Rep2** | **hDHFR Rep3** |
|---|---|---|---|
| Protein states | APO/MTX | APO/TMP/MTX | APO/TMP/MTX |
| Date | 07/05/24 | 07/06/24 | 10/20/24 |
| Protease column | FP/pepsin | Nep/pepsin | AP/pepsin |
| Reaction details | 50 mM potassium phosphate, 1 mM NaCl, 10 mM BME, pH 7.0, temperature 20 °C, final $D_2O$ concentration 90% | | |
| Time course (s) | 4.3e1-4.0e4 | 4.3e1-1.0e4 | 4.2e1-2.2e4 |
| # of time points | 5 | 4 | 5 |
| Control samples | Maximally-labeled sample (full-D) | | |
| Back-exchange (mean / IQR) | 0.23/0.10 | 0.22/0.09 | 0.21/0.09 |
| # of peptides | 151 | 103 | 112 |
| Sequence coverage | 0.88 | 0.74 | 0.68 |
| Average peptide length / Redundancy | 1.4 | 2.2 | 2.0 |
| Significant difference in $\Delta$ $log\ PF$ | $\Delta\ log\ PF > \frac{\sum_{states} std(log\ PF)}{\sqrt{n_{bootstrap}}}$ | | |

**Table S10. $\Delta G_{op}$ values for hDHFR.**

| Res. name | Res. number | Apo | Std Dev | Single resolved | TMP | Std Dev | Single resolved | MTX | Std Dev | Single resolved |
|---|---|---|---|---|---|---|---|---|---|---|
| VAL | 1 | 26 | 2 | FALSE | | | | 25 | 1 | FALSE |
| GLY | 2 | 28 | 2 | FALSE | | | | 28 | 1 | FALSE |
| SER | 3 | 13 | 3 | TRUE | 16 | 0 | TRUE | 34 | 25 | TRUE |
| LEU | 4 | 22 | 1 | FALSE | 22 | 1 | FALSE | 19 | 1 | FALSE |
| ASN | 5 | 25 | 1 | FALSE | 25 | 1 | FALSE | 22 | 1 | FALSE |
| CYS | 6 | 29 | 1 | FALSE | 29 | 1 | FALSE | 26 | 1 | FALSE |
| ILE | 7 | 27 | 1 | FALSE | 46 | 3 | FALSE | 45 | 3 | FALSE |
| VAL | 8 | 23 | 1 | FALSE | 42 | 3 | FALSE | 41 | 3 | FALSE |
| ALA | 9 | 26 | 0 | TRUE | 46 | 3 | FALSE | 45 | 3 | FALSE |
| VAL | 10 | | | | | | | | | |
| SER | 11 | | | | | | | | | |
| GLN | 12 | | | | | | | | | |
| ASN | 13 | | | | | | | | | |
| MET | 14 | | | | | | | | | |
| GLY | 15 | | | | | | | | | |
| ILE | 16 | | | | | | | | | |
| GLY | 17 | | | | | | | | | |
| LYS | 18 | 19 | 0 | FALSE | | 0 | | 28 | 1 | FALSE |
| ASN | 19 | 22 | 0 | FALSE | | 0 | | 31 | 1 | FALSE |
| GLY | 20 | 20 | 0 | FALSE | | 0 | | 29 | 1 | FALSE |
| ASP | 21 | 20 | 0 | FALSE | | 0 | | 29 | 1 | FALSE |
| LEU | 22 | 14 | 0 | FALSE | | 0 | | 23 | 1 | FALSE |
| PRO | 23 | | | | | | | | | |
| TRP | 24 | 15 | 0 | FALSE | | 0 | | 24 | 1 | FALSE |
| PRO | 25 | | | | | | | | | |
| PRO | 26 | | | | | | | | | |
| LEU | 27 | 14 | 0 | FALSE | | 0 | | 23 | 1 | FALSE |
| ARG | 28 | 18 | 0 | FALSE | | 0 | | 27 | 1 | FALSE |
| ASN | 29 | 7 | 3 | TRUE | | | | 8 | 3 | TRUE |

| | | | | | | | | | | |
|---|---|---|---|---|---|---|---|---|---|---|
| GLU | 30 | 18 | 0 | TRUE | 32 | 4 | FALSE | 18 | 0 | TRUE |
| PHE | 31 | 28 | 0 | FALSE | 29 | 4 | FALSE | 41 | 1 | FALSE |
| ARG | 32 | 31 | 0 | FALSE | 32 | 4 | FALSE | 44 | 1 | FALSE |
| TYR | 33 | 18 | 0 | TRUE | 26 | 0 | TRUE | 29 | 1 | TRUE |
| PHE | 34 | 22 | 0 | TRUE | 30 | 0 | TRUE | 36 | 0 | TRUE |
| GLN | 35 | 20 | 1 | TRUE | 29 | 1 | TRUE | 73 | 4 | TRUE |
| ARG | 36 | 17 | 1 | TRUE | 26 | 0 | TRUE | 34 | 0 | TRUE |
| MET | 37 | 22 | 0 | TRUE | 30 | 2 | TRUE | 34 | 0 | TRUE |
| THR | 38 | 18 | 0 | TRUE | 32 | 0 | TRUE | 36 | 0 | TRUE |
| THR | 39 | 22 | 0 | TRUE | 26 | 0 | TRUE | 36 | 0 | TRUE |
| THR | 40 | 18 | 0 | TRUE | 13 | 2 | FALSE | 11 | 8 | TRUE |
| SER | 41 | 10 | 5 | TRUE | 16 | 2 | FALSE | 18 | 8 | TRUE |
| SER | 42 | 19 | 2 | TRUE | 20 | 1 | TRUE | 22 | 0 | TRUE |
| VAL | 43 | 11 | 4 | TRUE | 5 | 3 | TRUE | 11 | 1 | TRUE |
| GLU | 44 | 11 | 2 | TRUE | 27 | 1 | FALSE | 6 | 3 | TRUE |
| GLY | 45 | 23 | 1 | FALSE | 27 | 1 | FALSE | 32 | 2 | FALSE |
| LYS | 46 | 25 | 1 | FALSE | 29 | 1 | FALSE | 33 | 2 | FALSE |
| GLN | 47 | 25 | 1 | FALSE | 29 | 1 | FALSE | 34 | 2 | FALSE |
| ASN | 48 | 28 | 1 | FALSE | 32 | 1 | FALSE | 36 | 2 | FALSE |
| LEU | 49 | 22 | 1 | FALSE | 27 | 1 | FALSE | 31 | 2 | FALSE |
| VAL | 50 | | | | | | | | | |
| ILE | 51 | | | | | | | | | |
| MET | 52 | 23 | 2 | TRUE | 18 | 2 | TRUE | 32 | 3 | TRUE |
| GLY | 53 | 17 | 1 | FALSE | 23 | 1 | FALSE | 20 | 2 | FALSE |
| LYS | 54 | 17 | 1 | FALSE | 23 | 1 | FALSE | 20 | 2 | FALSE |
| LYS | 55 | 17 | 1 | FALSE | 23 | 1 | FALSE | 33 | 9 | FALSE |
| THR | 56 | 17 | 1 | FALSE | 23 | 1 | FALSE | 32 | 9 | FALSE |
| TRP | 57 | 19 | 4 | TRUE | 22 | 1 | FALSE | 45 | 20 | TRUE |
| PHE | 58 | 27 | 2 | TRUE | 58 | 14 | TRUE | 69 | 7 | TRUE |
| SER | 59 | 28 | 0 | TRUE | | 0 | | 62 | 14 | TRUE |
| ILE | 60 | 6 | 2 | FALSE | 19 | 0 | FALSE | 5 | 1 | FALSE |

| | | | | | | | | | | |
|---|---|---|---|---|---|---|---|---|---|---|
| PRO | 61 | 0 | 0 | FALSE | 0 | 0 | FALSE | 0 | 0 | FALSE |
| GLU | 62 | 13 | 3 | TRUE | 20 | 0 | FALSE | 24 | 9 | TRUE |
| LYS | 63 | 20 | 2 | FALSE | 21 | 0 | FALSE | 45 | 7 | FALSE |
| ASN | 64 | 24 | 2 | FALSE | 25 | 0 | FALSE | 49 | 7 | FALSE |
| ARG | 65 | 34 | 0 | FALSE | 24 | 0 | FALSE | 34 | 1 | FALSE |
| PRO | 66 | | | | | | | | | |
| LEU | 67 | 13 | 1 | FALSE | 17 | 0 | FALSE | 43 | 2 | FALSE |
| LYS | 68 | 17 | 1 | FALSE | 20 | 0 | FALSE | 46 | 2 | FALSE |
| GLY | 69 | 19 | 1 | FALSE | 22 | 0 | FALSE | 48 | 2 | FALSE |
| ARG | 70 | 19 | 1 | FALSE | 23 | 0 | FALSE | 49 | 2 | FALSE |
| ILE | 71 | 15 | 1 | TRUE | 19 | 0 | FALSE | 18 | 0 | TRUE |
| ASN | 72 | 34 | 0 | TRUE | 74 | 3 | TRUE | 72 | 4 | TRUE |
| LEU | 73 | 30 | 0 | TRUE | 72 | 1 | FALSE | 34 | 1 | TRUE |
| VAL | 74 | 27 | 0 | FALSE | 69 | 1 | FALSE | 51 | 4 | FALSE |
| LEU | 75 | 28 | 0 | FALSE | 70 | 1 | FALSE | 52 | 4 | FALSE |
| SER | 76 | 26 | 1 | FALSE | 28 | 1 | FALSE | 27 | 1 | FALSE |
| ARG | 77 | 27 | 1 | FALSE | 29 | 1 | FALSE | 29 | 1 | FALSE |
| GLU | 78 | 30 | 1 | TRUE | 29 | 4 | TRUE | 21 | 10 | TRUE |
| LEU | 79 | 3 | 1 | TRUE | 3 | 1 | TRUE | 12 | 4 | TRUE |
| LYS | 80 | 4 | 2 | TRUE | 4 | 2 | TRUE | 4 | 2 | TRUE |
| GLU | 81 | 17 | 1 | TRUE | 13 | 1 | TRUE | 15 | 1 | TRUE |
| PRO | 82 | | | | | | | | | |
| PRO | 83 | | | | | | | | | |
| GLN | 84 | 14 | 1 | FALSE | 12 | 0 | FALSE | 18 | 1 | FALSE |
| GLY | 85 | 16 | 1 | FALSE | 13 | 0 | FALSE | 20 | 1 | FALSE |
| ALA | 86 | 16 | 1 | FALSE | 13 | 0 | FALSE | 20 | 1 | FALSE |
| HIS | 87 | 18 | 5 | TRUE | 15 | 0 | FALSE | 16 | 7 | TRUE |
| PHE | 88 | 17 | 4 | TRUE | 14 | 0 | FALSE | 11 | 4 | TRUE |
| LEU | 89 | 21 | 4 | TRUE | 31 | 1 | TRUE | 17 | 2 | TRUE |
| SER | 90 | 27 | 6 | TRUE | 73 | 5 | TRUE | 27 | 8 | TRUE |
| ARG | 91 | 24 | 0 | TRUE | 30 | 2 | TRUE | 26 | 2 | TRUE |

| | | | | | | | | | | |
|---|---|---|---|---|---|---|---|---|---|---|
| SER | 92 | 13 | 2 | FALSE | 12 | 2 | FALSE | 11 | 2 | FALSE |
| LEU | 93 | 8 | 2 | FALSE | 8 | 2 | FALSE | 6 | 2 | FALSE |
| ASP | 94 | 15 | 1 | TRUE | 22 | 1 | TRUE | 15 | 1 | TRUE |
| ASP | 95 | 25 | 2 | TRUE | 10 | 6 | TRUE | 27 | 4 | TRUE |
| ALA | 96 | 34 | 0 | TRUE | 72 | 5 | TRUE | 65 | 13 | TRUE |
| LEU | 97 | 30 | 0 | TRUE | 47 | 18 | TRUE | 40 | 16 | TRUE |
| LYS | 98 | 26 | 0 | TRUE | 28 | 0 | TRUE | 26 | 3 | TRUE |
| LEU | 99 | 22 | 0 | TRUE | 20 | 0 | TRUE | 25 | 2 | TRUE |
| THR | 100 | 18 | 0 | TRUE | 18 | 0 | TRUE | 19 | 1 | TRUE |
| GLU | 101 | 22 | 0 | TRUE | 21 | 1 | TRUE | 21 | 1 | TRUE |
| GLN | 102 | 24 | 0 | TRUE | 22 | 2 | TRUE | 25 | 1 | TRUE |
| PRO | 103 | | | | | | | | | |
| GLU | 104 | 11 | 1 | TRUE | 10 | 0 | FALSE | 12 | 0 | TRUE |
| LEU | 105 | 14 | 0 | TRUE | 17 | 2 | TRUE | 15 | 1 | TRUE |
| ALA | 106 | 20 | 0 | TRUE | 20 | 1 | TRUE | 21 | 1 | TRUE |
| ASN | 107 | 18 | 4 | TRUE | 7 | 4 | TRUE | 9 | 4 | TRUE |
| LYS | 108 | 19 | 3 | TRUE | 19 | 3 | TRUE | 18 | 0 | TRUE |
| VAL | 109 | 17 | 3 | TRUE | 20 | 0 | FALSE | 18 | 0 | TRUE |
| ASP | 110 | 28 | 5 | TRUE | 23 | 0 | FALSE | 24 | 0 | TRUE |
| MET | 111 | 27 | 4 | TRUE | 70 | 6 | TRUE | 34 | 0 | TRUE |
| VAL | 112 | 26 | 1 | TRUE | 29 | 1 | TRUE | 29 | 1 | TRUE |
| TRP | 113 | 31 | 1 | TRUE | 51 | 19 | TRUE | 31 | 1 | TRUE |
| ILE | 114 | 23 | 3 | TRUE | 69 | 7 | TRUE | 48 | 25 | TRUE |
| VAL | 115 | 68 | 8 | TRUE | 69 | 6 | TRUE | 68 | 7 | TRUE |
| GLY | 116 | 17 | 1 | FALSE | 28 | 2 | FALSE | 24 | 1 | FALSE |
| GLY | 117 | 18 | 1 | FALSE | 29 | 2 | FALSE | 26 | 1 | FALSE |
| SER | 118 | 21 | 1 | FALSE | 32 | 2 | FALSE | 28 | 1 | FALSE |
| SER | 119 | 21 | 1 | FALSE | 32 | 2 | FALSE | 29 | 1 | FALSE |
| VAL | 120 | 12 | 4 | TRUE | 10 | 4 | TRUE | 10 | 0 | TRUE |
| TYR | 121 | 20 | 1 | TRUE | 22 | 2 | TRUE | 20 | 0 | TRUE |
| LYS | 122 | 16 | 0 | TRUE | 22 | 2 | TRUE | 14 | 0 | TRUE |

| | | | | | | | | | | |
|---|---|---|---|---|---|---|---|---|---|---|
| GLU | 123 | 21 | 2 | TRUE | 24 | 0 | TRUE | 20 | 0 | TRUE |
| ALA | 124 | 30 | 0 | TRUE | 31 | 5 | TRUE | 26 | 9 | TRUE |
| MET | 125 | 23 | 4 | TRUE | 19 | 1 | TRUE | 33 | 3 | TRUE |
| ASN | 126 | 8 | 3 | TRUE | 9 | 6 | TRUE | 18 | 15 | TRUE |
| HIS | 127 | 24 | 1 | FALSE | 24 | 1 | FALSE | 43 | 2 | FALSE |
| PRO | 128 | | | | | | | | | |
| GLY | 129 | 18 | 1 | FALSE | 18 | 1 | FALSE | 37 | 2 | FALSE |
| HIS | 130 | 23 | 1 | FALSE | 23 | 1 | FALSE | 42 | 2 | FALSE |
| LEU | 131 | 46 | 6 | FALSE | | | | 42 | 5 | FALSE |
| LYS | 132 | 45 | 6 | FALSE | | | | 41 | 5 | FALSE |
| LEU | 133 | 66 | 8 | TRUE | | | | 68 | 7 | TRUE |
| PHE | 134 | | | | | | | | | |
| VAL | 135 | 26 | 0 | TRUE | 33 | 3 | TRUE | 69 | 7 | TRUE |
| THR | 136 | 23 | 2 | TRUE | 30 | 0 | TRUE | 30 | 1 | TRUE |
| ARG | 137 | 28 | 0 | TRUE | 73 | 4 | TRUE | 73 | 4 | TRUE |
| ILE | 138 | 24 | 0 | TRUE | 71 | 5 | TRUE | 43 | 18 | TRUE |
| MET | 139 | 24 | 1 | TRUE | 26 | 0 | TRUE | 31 | 1 | TRUE |
| GLN | 140 | 21 | 1 | TRUE | 28 | 0 | TRUE | 30 | 0 | TRUE |
| ASP | 141 | 17 | 1 | TRUE | 17 | 1 | TRUE | 17 | 1 | TRUE |
| PHE | 142 | 13 | 1 | TRUE | 16 | 2 | TRUE | 14 | 2 | TRUE |
| GLU | 143 | 11 | 1 | TRUE | 5 | 2 | TRUE | 7 | 3 | TRUE |
| SER | 144 | 14 | 1 | TRUE | 18 | 2 | FALSE | 17 | 2 | TRUE |
| ASP | 145 | 29 | 5 | TRUE | 19 | 2 | FALSE | 34 | 0 | TRUE |
| THR | 146 | 16 | 0 | TRUE | 20 | 0 | TRUE | 24 | 0 | TRUE |
| PHE | 147 | 14 | 0 | TRUE | 14 | 0 | TRUE | 10 | 3 | TRUE |
| PHE | 148 | 19 | 1 | TRUE | 30 | 0 | TRUE | 32 | 0 | TRUE |
| PRO | 149 | | | | | | | | | |
| GLU | 150 | 14 | 0 | TRUE | 15 | 1 | TRUE | 15 | 1 | TRUE |
| ILE | 151 | 9 | 1 | TRUE | 11 | 4 | TRUE | 10 | 2 | TRUE |
| ASP | 152 | 17 | 4 | TRUE | 12 | 1 | TRUE | 17 | 4 | TRUE |
| LEU | 153 | 12 | 2 | TRUE | 12 | 3 | TRUE | 11 | 1 | TRUE |

| | | | | | | | | | | |
|---|---|---|---|---|---|---|---|---|---|---|
| GLU | 154 | 18 | 2 | TRUE | 16 | 1 | TRUE | 18 | 3 | TRUE |
| LYS | 155 | 16 | 0 | TRUE | 16 | 1 | TRUE | 18 | 3 | TRUE |
| TYR | 156 | 26 | 1 | TRUE | 57 | 22 | TRUE | 36 | 2 | TRUE |
| LYS | 157 | 24 | 4 | TRUE | 31 | 1 | TRUE | 36 | 2 | TRUE |
| LEU | 158 | 19 | 4 | TRUE | 16 | 1 | TRUE | 16 | 2 | TRUE |
| LEU | 159 | 24 | 0 | TRUE | 40 | 23 | TRUE | 43 | 23 | TRUE |
| PRO | 160 | | | | | | | | | |
| GLU | 161 | 8 | 4 | TRUE | 12 | 0 | TRUE | 8 | 3 | TRUE |
| TYR | 162 | 12 | 3 | FALSE | 9 | 2 | FALSE | 8 | 1 | FALSE |
| PRO | 163 | | | | | | | | | |
| GLY | 164 | 9 | 1 | FALSE | 8 | 1 | FALSE | 12 | 1 | FALSE |
| VAL | 165 | 7 | 1 | FALSE | 7 | 1 | FALSE | 11 | 1 | FALSE |
| LEU | 166 | 13 | 1 | TRUE | 15 | 1 | TRUE | 15 | 1 | TRUE |
| SER | 167 | 5 | 2 | TRUE | 7 | 4 | TRUE | 6 | 3 | TRUE |
| ASP | 168 | 17 | 1 | TRUE | 15 | 2 | TRUE | 15 | 1 | TRUE |
| VAL | 169 | 11 | 2 | FALSE | 11 | 2 | FALSE | 12 | 1 | FALSE |
| GLN | 170 | 15 | 2 | FALSE | 15 | 2 | FALSE | 16 | 1 | FALSE |
| GLU | 171 | 17 | 1 | FALSE | 28 | 1 | FALSE | 17 | 2 | FALSE |
| GLU | 172 | 15 | 1 | FALSE | 26 | 1 | FALSE | 15 | 2 | FALSE |
| LYS | 173 | 15 | 1 | FALSE | 26 | 1 | FALSE | 15 | 2 | FALSE |
| GLY | 174 | 19 | 1 | FALSE | 28 | 1 | FALSE | 37 | 3 | FALSE |
| ILE | 175 | 16 | 1 | FALSE | 24 | 1 | FALSE | 34 | 3 | FALSE |
| LYS | 176 | 17 | 1 | FALSE | 26 | 1 | FALSE | 35 | 3 | FALSE |
| TYR | 177 | 21 | 1 | TRUE | 27 | 1 | FALSE | 37 | 7 | TRUE |
| LYS | 178 | 23 | 1 | TRUE | 27 | 1 | FALSE | 40 | 5 | TRUE |
| PHE | 179 | 24 | 0 | TRUE | 70 | 6 | TRUE | 52 | 14 | TRUE |
| GLU | 180 | 28 | 0 | TRUE | 38 | 4 | TRUE | 43 | 17 | TRUE |
| VAL | 181 | 24 | 0 | TRUE | 54 | 20 | TRUE | 67 | 6 | TRUE |
| TYR | 182 | 21 | 1 | FALSE | 27 | 1 | FALSE | 50 | 3 | FALSE |
| GLU | 183 | 23 | 1 | FALSE | 29 | 1 | FALSE | 52 | 3 | FALSE |
| LYS | 184 | 25 | 2 | TRUE | 46 | 14 | TRUE | 32 | 25 | TRUE |

| | | | | | | | | | | |
|---|---|---|---|---|---|---|---|---|---|---|
| ASN | 185 | 22 | 1 | FALSE | 19 | 1 | FALSE | 27 | 5 | FALSE |
| ASP | 186 | 11 | 1 | FALSE | 8 | 1 | FALSE | 15 | 5 | FALSE |

**Table S11. HX experiment summary table for LacI.**

All experiments run with KEEP setting in PIGEON. Abbreviations: FP, immobilized protease type XIII. Nep, nepenthesin 2. AP, alanyl aminopeptidase.

| Dataset | **LacI Rep1** | **LacI Rep2** | **LacI Rep3** | **LacI Rep4** |
|---|---|---|---|---|
| Date | 11/18/22 | 01/14/23 | 03/15/24 | 08/08/24 |
| Protease column | FP/pepsin | FP/pepsin | FP/pepsin | AP/pepsin |
| Protein states | APO/DNA/IPTG | APO/DNA/IPTG | APO/DNA/IPTG | APO/DNA/IPTG |
| Reaction details | 50mM Tris, 150mM NaCl, pH 8.0, temperature 15 °C | | | |
| Time course (s) | 4.5e1 to 3.9e3 | 4.5e1 to 1.5e4 | 4.5e1 to 1.5e4 | 4.5e1 to 4.0e4 |
| # of time points | 6 | 5 | 8 | 9 |
| Control samples | Maximally labeled sample (full-D) | | | |
| Back-exchange (mean / IQR) | 0.20/0.12 | 0.20/0.11 | 0.23/0.10 | 0.24/0.10 |
| # of peptides | 189 | 187 | 240 | 240 |
| Sequence coverage | 0.81 | 0.79 | 0.91 | 0.87 |
| Average peptide length / Redundancy | 1.9 | 2.0 | 1.5 | 1.5 |
| Significant difference in $\Delta$ $log\ PF$ | $\Delta\ log\ PF > \frac{\sum_{states}\ std(log\ PF)}{\sqrt{n_{bootstrap}}}$ | | | |

**Table S12. $\Delta G_{op}$ values for LacI.**

| Res. name | Res. number | APO | Std Dev | Single resolved | DNA | Std Dev | Single resolved | IPTG | Std Dev | Single resolved |
|---|---|---|---|---|---|---|---|---|---|---|
| MET | 1 | | | | | | | | | |
| LYS | 2 | | | | | | | | | |
| PRO | 3 | | | | | | | | | |
| VAL | 4 | | | | | | | | | |
| THR | 5 | 40 | 2 | FALSE | 24 | 2 | FALSE | 39 | 2 | FALSE |
| LEU | 6 | 39 | 2 | FALSE | 23 | 2 | FALSE | 38 | 2 | FALSE |
| TYR | 7 | 38 | 2 | FALSE | 23 | 2 | FALSE | 37 | 2 | FALSE |
| ASP | 8 | 42 | 2 | FALSE | 26 | 2 | FALSE | 41 | 2 | FALSE |
| VAL | 9 | 27 | 3 | FALSE | 50 | 3 | FALSE | 12 | 1 | FALSE |
| ALA | 10 | 31 | 3 | FALSE | 54 | 3 | FALSE | 16 | 1 | FALSE |
| GLU | 11 | 32 | 3 | FALSE | 54 | 3 | FALSE | 16 | 1 | FALSE |
| TYR | 12 | 16 | 6 | TRUE | 32 | 0 | TRUE | 14 | 0 | TRUE |
| ALA | 13 | | | | | | | | | |
| GLY | 14 | 11 | 1 | FALSE | 19 | 0 | FALSE | 15 | 1 | FALSE |
| VAL | 15 | 8 | 1 | FALSE | 16 | 0 | FALSE | 12 | 1 | FALSE |
| SER | 16 | 12 | 1 | FALSE | 20 | 0 | FALSE | 16 | 1 | FALSE |
| TYR | 17 | 11 | 6 | TRUE | 26 | 0 | TRUE | 9 | 5 | TRUE |
| GLN | 18 | 15 | 5 | TRUE | 30 | 4 | TRUE | 9 | 5 | TRUE |
| THR | 19 | 20 | 0 | TRUE | 19 | 1 | TRUE | 18 | 1 | TRUE |
| VAL | 20 | 7 | 4 | TRUE | 14 | 7 | TRUE | 9 | 5 | TRUE |
| SER | 21 | 26 | 2 | FALSE | 36 | 0 | FALSE | 28 | 1 | FALSE |
| ARG | 22 | 27 | 2 | FALSE | 37 | 0 | FALSE | 29 | 1 | FALSE |
| VAL | 23 | 6 | 3 | TRUE | 26 | 0 | TRUE | 7 | 4 | TRUE |
| VAL | 24 | 5 | 2 | TRUE | 26 | 0 | TRUE | 5 | 2 | TRUE |
| ASN | 25 | 9 | 1 | FALSE | 18 | 1 | FALSE | 9 | 1 | FALSE |
| GLN | 26 | 9 | 1 | FALSE | 19 | 1 | FALSE | 9 | 1 | FALSE |
| ALA | 27 | 8 | 1 | FALSE | 18 | 1 | FALSE | 8 | 1 | FALSE |
| SER | 28 | 8 | 1 | FALSE | 17 | 1 | FALSE | 10 | 1 | FALSE |
| HIS | 29 | 8 | 1 | FALSE | 17 | 1 | FALSE | 10 | 1 | FALSE |

| | | | | | | | | | | |
|---|---|---|---|---|---|---|---|---|---|---|
| VAL | 30 | 4 | 1 | FALSE | 12 | 1 | FALSE | 5 | 1 | FALSE |
| SER | 31 | 7 | 1 | FALSE | 16 | 1 | FALSE | 9 | 1 | FALSE |
| ALA | 32 | 32 | 1 | TRUE | 34 | 0 | TRUE | 31 | 2 | TRUE |
| LYS | 33 | 53 | 20 | TRUE | 66 | 8 | TRUE | 62 | 11 | TRUE |
| THR | 34 | 53 | 19 | TRUE | 64 | 11 | TRUE | 55 | 19 | TRUE |
| ARG | 35 | 68 | 6 | TRUE | 70 | 6 | TRUE | 67 | 6 | TRUE |
| GLU | 36 | 38 | 4 | FALSE | 50 | 2 | FALSE | 55 | 4 | FALSE |
| LYS | 37 | 36 | 4 | FALSE | 48 | 2 | FALSE | 53 | 4 | FALSE |
| VAL | 38 | 34 | 4 | FALSE | 46 | 2 | FALSE | 51 | 4 | FALSE |
| GLU | 39 | 16 | 0 | TRUE | 29 | 1 | TRUE | 11 | 7 | TRUE |
| ALA | 40 | 24 | 1 | FALSE | 32 | 0 | FALSE | 27 | 0 | FALSE |
| ALA | 41 | 24 | 1 | FALSE | 33 | 0 | FALSE | 27 | 0 | FALSE |
| MET | 42 | 10 | 6 | TRUE | 24 | 0 | TRUE | 9 | 5 | TRUE |
| ALA | 43 | 25 | 4 | TRUE | 10 | 6 | TRUE | 26 | 7 | TRUE |
| GLU | 44 | 8 | 4 | TRUE | 7 | 3 | TRUE | 12 | 5 | TRUE |
| LEU | 45 | 14 | 3 | TRUE | 18 | 0 | TRUE | 14 | 1 | TRUE |
| ASN | 46 | 8 | 4 | TRUE | 12 | 7 | TRUE | 9 | 5 | TRUE |
| TYR | 47 | 16 | 10 | TRUE | 28 | 0 | TRUE | 13 | 10 | TRUE |
| ILE | 48 | 19 | 2 | FALSE | 24 | 1 | FALSE | 20 | 3 | FALSE |
| PRO | 49 | | | | | | | | | |
| ASN | 50 | 10 | 7 | TRUE | 24 | 3 | TRUE | 8 | 4 | TRUE |
| ARG | 51 | 8 | 4 | TRUE | 20 | 1 | TRUE | 8 | 4 | TRUE |
| VAL | 52 | 6 | 1 | FALSE | 15 | 0 | FALSE | 5 | 1 | FALSE |
| ALA | 53 | 8 | 1 | FALSE | 17 | 0 | FALSE | 7 | 1 | FALSE |
| GLN | 54 | 8 | 4 | TRUE | 26 | 0 | TRUE | 7 | 4 | TRUE |
| GLN | 55 | 8 | 4 | TRUE | 9 | 4 | TRUE | 8 | 4 | TRUE |
| LEU | 56 | 6 | 3 | TRUE | 6 | 3 | TRUE | 6 | 3 | TRUE |
| ALA | 57 | 7 | 3 | TRUE | 8 | 4 | TRUE | 7 | 3 | TRUE |
| GLY | 58 | 13 | 9 | TRUE | 26 | 0 | TRUE | 9 | 5 | TRUE |
| LYS | 59 | 8 | 4 | TRUE | 20 | 1 | TRUE | 7 | 4 | TRUE |
| GLN | 60 | 8 | 4 | TRUE | 14 | 5 | TRUE | 8 | 4 | TRUE |

| | | | | | | | | | | |
|---|---|---|---|---|---|---|---|---|---|---|
| SER | 61 | 9 | 2 | FALSE | 24 | 3 | FALSE | 19 | 3 | FALSE |
| LEU | 62 | 5 | 2 | FALSE | 19 | 3 | FALSE | 15 | 3 | FALSE |
| LEU | 63 | 14 | 10 | TRUE | 5 | 2 | TRUE | 12 | 9 | TRUE |
| ILE | 64 | 11 | 8 | TRUE | 24 | 0 | TRUE | 10 | 6 | TRUE |
| GLY | 65 | 40 | 0 | TRUE | 38 | 0 | TRUE | 68 | 7 | TRUE |
| VAL | 66 | 60 | 4 | FALSE | 66 | 3 | FALSE | 66 | 2 | FALSE |
| ALA | 67 | 62 | 4 | FALSE | 68 | 3 | FALSE | 68 | 2 | FALSE |
| THR | 68 | 10 | 2 | FALSE | 16 | 0 | FALSE | 11 | 2 | FALSE |
| SER | 69 | 14 | 2 | FALSE | 20 | 0 | FALSE | 14 | 2 | FALSE |
| SER | 70 | 21 | 1 | TRUE | 23 | 1 | TRUE | 18 | 7 | TRUE |
| LEU | 71 | 36 | 0 | TRUE | 36 | 0 | TRUE | 44 | 11 | TRUE |
| ALA | 72 | 14 | 3 | TRUE | 25 | 1 | TRUE | 10 | 5 | TRUE |
| LEU | 73 | 6 | 3 | TRUE | 9 | 5 | TRUE | 16 | 0 | TRUE |
| HIS | 74 | 28 | 2 | TRUE | 31 | 1 | TRUE | 30 | 1 | TRUE |
| ALA | 75 | 13 | 6 | TRUE | 10 | 6 | TRUE | 22 | 0 | TRUE |
| PRO | 76 | | | | | | | | | |
| SER | 77 | 23 | 3 | TRUE | 32 | 0 | TRUE | 29 | 1 | TRUE |
| GLN | 78 | 23 | 3 | TRUE | 22 | 0 | TRUE | 28 | 0 | TRUE |
| ILE | 79 | 22 | 1 | FALSE | 32 | 1 | FALSE | 51 | 4 | FALSE |
| VAL | 80 | 20 | 1 | FALSE | 30 | 1 | FALSE | 49 | 4 | FALSE |
| ALA | 81 | 35 | 1 | TRUE | 55 | 19 | TRUE | 67 | 7 | TRUE |
| ALA | 82 | 30 | 0 | TRUE | 35 | 3 | TRUE | 33 | 1 | TRUE |
| ILE | 83 | 23 | 4 | TRUE | 25 | 4 | TRUE | 16 | 0 | TRUE |
| LYS | 84 | 34 | 3 | TRUE | 52 | 15 | TRUE | 30 | 2 | TRUE |
| SER | 85 | 21 | 3 | FALSE | 34 | 1 | FALSE | 17 | 3 | FALSE |
| ARG | 86 | 20 | 3 | FALSE | 33 | 1 | FALSE | 17 | 3 | FALSE |
| ALA | 87 | 26 | 0 | FALSE | 21 | 2 | FALSE | 29 | 1 | FALSE |
| ASP | 88 | 25 | 0 | FALSE | 20 | 2 | FALSE | 28 | 1 | FALSE |
| GLN | 89 | 24 | 0 | FALSE | 19 | 2 | FALSE | 27 | 1 | FALSE |
| LEU | 90 | 29 | 1 | TRUE | 31 | 1 | TRUE | 30 | 3 | TRUE |
| GLY | 91 | 27 | 2 | TRUE | 31 | 2 | TRUE | 26 | 6 | TRUE |

| | | | | | | | | | | |
|---|---|---|---|---|---|---|---|---|---|---|
| ALA | 92 | 32 | 1 | TRUE | 46 | 13 | TRUE | 37 | 1 | TRUE |
| SER | 93 | 30 | 0 | TRUE | 28 | 2 | TRUE | 30 | 2 | TRUE |
| VAL | 94 | 6 | 3 | TRUE | 10 | 5 | TRUE | 6 | 3 | TRUE |
| VAL | 95 | 29 | 1 | TRUE | 30 | 0 | TRUE | 28 | 4 | TRUE |
| VAL | 96 | 37 | 3 | FALSE | 29 | 4 | FALSE | 31 | 4 | FALSE |
| SER | 97 | 43 | 3 | FALSE | 35 | 4 | FALSE | 37 | 4 | FALSE |
| MET | 98 | 43 | 3 | FALSE | 35 | 4 | FALSE | 38 | 4 | FALSE |
| VAL | 99 | | | | | | | | | |
| GLU | 100 | 6 | 3 | TRUE | 7 | 3 | TRUE | 7 | 3 | TRUE |
| ARG | 101 | 7 | 1 | FALSE | 7 | 1 | FALSE | 5 | 2 | FALSE |
| SER | 102 | 10 | 1 | FALSE | 10 | 1 | FALSE | 9 | 2 | FALSE |
| GLY | 103 | 8 | 1 | FALSE | 12 | 1 | FALSE | 9 | 1 | FALSE |
| VAL | 104 | 3 | 1 | FALSE | 8 | 1 | FALSE | 5 | 1 | FALSE |
| GLU | 105 | 5 | 1 | FALSE | 9 | 1 | FALSE | 7 | 1 | FALSE |
| ALA | 106 | 8 | 4 | TRUE | 13 | 5 | TRUE | 10 | 5 | TRUE |
| CYS | 107 | 11 | 5 | TRUE | 10 | 5 | TRUE | 12 | 7 | TRUE |
| LYS | 108 | 30 | 0 | TRUE | 30 | 0 | TRUE | 28 | 3 | TRUE |
| ALA | 109 | 26 | 0 | TRUE | 22 | 0 | TRUE | 23 | 3 | TRUE |
| ALA | 110 | 34 | 2 | TRUE | 33 | 4 | TRUE | 31 | 1 | TRUE |
| VAL | 111 | 27 | 1 | TRUE | 28 | 4 | TRUE | 25 | 1 | TRUE |
| HIS | 112 | 20 | 0 | TRUE | 26 | 0 | TRUE | 22 | 3 | TRUE |
| ASN | 113 | 34 | 1 | TRUE | 41 | 1 | TRUE | 33 | 1 | TRUE |
| LEU | 114 | 67 | 6 | TRUE | 67 | 7 | TRUE | 37 | 2 | TRUE |
| LEU | 115 | 51 | 18 | TRUE | 56 | 18 | TRUE | 34 | 1 | TRUE |
| ALA | 116 | 25 | 1 | TRUE | 29 | 1 | TRUE | 22 | 4 | TRUE |
| GLN | 117 | 27 | 1 | TRUE | 34 | 0 | TRUE | 28 | 1 | TRUE |
| ARG | 118 | 19 | 1 | TRUE | 22 | 0 | TRUE | 18 | 1 | TRUE |
| VAL | 119 | 22 | 3 | FALSE | 29 | 2 | FALSE | 44 | 3 | FALSE |
| SER | 120 | 26 | 3 | FALSE | 33 | 2 | FALSE | 48 | 3 | FALSE |
| GLY | 121 | 33 | 6 | TRUE | 38 | 3 | TRUE | 68 | 6 | TRUE |
| LEU | 122 | 44 | 16 | TRUE | 34 | 4 | TRUE | 29 | 3 | TRUE |

| | | | | | | | | | | |
|---|---|---|---|---|---|---|---|---|---|---|
| ILE | 123 | 19 | 2 | TRUE | 26 | 0 | TRUE | 15 | 1 | TRUE |
| ILE | 124 | 45 | 21 | TRUE | 54 | 21 | TRUE | 30 | 1 | TRUE |
| ASN | 125 | 20 | 1 | TRUE | 23 | 2 | TRUE | 28 | 0 | TRUE |
| TYR | 126 | 66 | 5 | FALSE | 69 | 2 | FALSE | 71 | 2 | FALSE |
| PRO | 127 | | | | | | | | | |
| LEU | 128 | 68 | 6 | TRUE | 69 | 6 | TRUE | 69 | 6 | TRUE |
| ASP | 129 | 70 | 5 | TRUE | 70 | 5 | TRUE | 69 | 5 | TRUE |
| ASP | 130 | 34 | 6 | TRUE | 32 | 1 | TRUE | 68 | 7 | TRUE |
| GLN | 131 | 17 | 1 | TRUE | 14 | 3 | TRUE | 16 | 0 | TRUE |
| ASP | 132 | 20 | 0 | TRUE | 20 | 0 | TRUE | 20 | 0 | TRUE |
| ALA | 133 | 34 | 4 | TRUE | 33 | 2 | TRUE | 32 | 1 | TRUE |
| ILE | 134 | 49 | 20 | TRUE | 52 | 18 | TRUE | 35 | 4 | TRUE |
| ALA | 135 | 38 | 5 | TRUE | 57 | 21 | TRUE | 37 | 2 | TRUE |
| VAL | 136 | 25 | 1 | FALSE | 25 | 0 | FALSE | 25 | 0 | FALSE |
| GLU | 137 | 27 | 1 | FALSE | 28 | 0 | FALSE | 27 | 0 | FALSE |
| ALA | 138 | 25 | 2 | TRUE | 27 | 1 | TRUE | 26 | 0 | TRUE |
| ALA | 139 | 56 | 22 | TRUE | 62 | 13 | TRUE | 60 | 14 | TRUE |
| CYS | 140 | 22 | 2 | TRUE | 22 | 0 | TRUE | 23 | 2 | TRUE |
| THR | 141 | 12 | 7 | TRUE | 33 | 11 | TRUE | 10 | 5 | TRUE |
| ASN | 142 | 49 | 30 | TRUE | 11 | 5 | TRUE | 48 | 27 | TRUE |
| VAL | 143 | 12 | 4 | TRUE | 16 | 0 | TRUE | 10 | 5 | TRUE |
| PRO | 144 | | | | | | | | | |
| ALA | 145 | 48 | 3 | FALSE | 44 | 6 | FALSE | 48 | 6 | FALSE |
| LEU | 146 | 46 | 3 | FALSE | 42 | 6 | FALSE | 46 | 6 | FALSE |
| PHE | 147 | 67 | 7 | TRUE | 31 | 4 | TRUE | 30 | 3 | TRUE |
| LEU | 148 | 56 | 21 | TRUE | 67 | 6 | TRUE | 68 | 6 | TRUE |
| ASP | 149 | 28 | 0 | TRUE | 31 | 1 | TRUE | 33 | 1 | TRUE |
| VAL | 150 | 33 | 1 | TRUE | 62 | 12 | TRUE | 63 | 11 | TRUE |
| SER | 151 | 8 | 4 | TRUE | 12 | 6 | TRUE | 14 | 6 | TRUE |
| ASP | 152 | 16 | 8 | TRUE | 20 | 7 | TRUE | 22 | 8 | TRUE |
| GLN | 153 | 22 | 8 | TRUE | 22 | 8 | TRUE | 22 | 5 | TRUE |

| | | | | | | | | | | |
|---|---|---|---|---|---|---|---|---|---|---|
| THR | 154 | 20 | 0 | TRUE | 20 | 0 | TRUE | 20 | 2 | TRUE |
| PRO | 155 | | | | | | | | | |
| ILE | 156 | 15 | 3 | FALSE | 20 | 2 | FALSE | 21 | 2 | FALSE |
| ASN | 157 | 22 | 3 | FALSE | 27 | 2 | FALSE | 27 | 2 | FALSE |
| SER | 158 | 33 | 2 | TRUE | 32 | 1 | TRUE | 33 | 1 | TRUE |
| ILE | 159 | 30 | 0 | TRUE | 40 | 2 | TRUE | 67 | 7 | TRUE |
| ILE | 160 | 25 | 1 | TRUE | 31 | 1 | TRUE | 33 | 1 | TRUE |
| PHE | 161 | 30 | 0 | TRUE | 29 | 1 | TRUE | 59 | 12 | TRUE |
| SER | 162 | 49 | 5 | FALSE | 52 | 5 | FALSE | 33 | 3 | FALSE |
| HIS | 163 | 49 | 5 | FALSE | 51 | 5 | FALSE | 33 | 3 | FALSE |
| GLU | 164 | 35 | 1 | TRUE | 53 | 18 | TRUE | 51 | 23 | TRUE |
| ASP | 165 | 18 | 0 | TRUE | 18 | 0 | TRUE | 21 | 7 | TRUE |
| GLY | 166 | 18 | 0 | TRUE | 17 | 1 | TRUE | 33 | 22 | TRUE |
| THR | 167 | 30 | 1 | TRUE | 44 | 16 | TRUE | 65 | 7 | TRUE |
| ARG | 168 | 37 | 0 | FALSE | 49 | 8 | FALSE | 69 | 2 | FALSE |
| LEU | 169 | 34 | 0 | FALSE | 45 | 8 | FALSE | 66 | 2 | FALSE |
| GLY | 170 | 62 | 12 | TRUE | 63 | 9 | TRUE | 65 | 8 | TRUE |
| VAL | 171 | 66 | 8 | TRUE | 66 | 8 | TRUE | 66 | 7 | TRUE |
| GLU | 172 | 29 | 0 | FALSE | 31 | 1 | FALSE | 31 | 1 | FALSE |
| HIS | 173 | 30 | 0 | FALSE | 32 | 1 | FALSE | 31 | 1 | FALSE |
| LEU | 174 | 67 | 7 | TRUE | 66 | 7 | TRUE | 67 | 7 | TRUE |
| VAL | 175 | 61 | 12 | TRUE | 49 | 16 | TRUE | 49 | 20 | TRUE |
| ALA | 176 | 60 | 12 | TRUE | 54 | 14 | TRUE | 40 | 1 | TRUE |
| LEU | 177 | 16 | 0 | TRUE | 16 | 0 | TRUE | 16 | 0 | TRUE |
| GLY | 178 | 32 | 0 | TRUE | 33 | 2 | TRUE | 34 | 0 | TRUE |
| HIS | 179 | 39 | 1 | TRUE | 35 | 4 | TRUE | 38 | 0 | TRUE |
| GLN | 180 | 30 | 0 | TRUE | 52 | 22 | TRUE | 30 | 0 | TRUE |
| GLN | 181 | 8 | 4 | TRUE | 45 | 19 | TRUE | 10 | 5 | TRUE |
| ILE | 182 | 47 | 28 | TRUE | 6 | 3 | TRUE | 67 | 7 | TRUE |
| ALA | 183 | 66 | 6 | TRUE | 56 | 22 | TRUE | 68 | 6 | TRUE |
| LEU | 184 | 66 | 7 | TRUE | 61 | 11 | TRUE | 61 | 11 | TRUE |

| | | | | | | | | | | |
|---|---|---|---|---|---|---|---|---|---|---|
| LEU | 185 | 34 | 0 | TRUE | 48 | 16 | TRUE | 36 | 1 | TRUE |
| ALA | 186 | 31 | 2 | TRUE | 33 | 2 | TRUE | 36 | 2 | TRUE |
| GLY | 187 | 28 | 5 | TRUE | 30 | 7 | TRUE | 30 | 5 | TRUE |
| PRO | 188 | | | | | | | | | |
| LEU | 189 | 17 | 6 | TRUE | 17 | 3 | TRUE | 18 | 6 | TRUE |
| SER | 190 | 23 | 3 | FALSE | 19 | 4 | FALSE | 19 | 3 | FALSE |
| SER | 191 | 26 | 3 | FALSE | 22 | 4 | FALSE | 22 | 3 | FALSE |
| VAL | 192 | 15 | 1 | TRUE | 16 | 1 | TRUE | 20 | 10 | TRUE |
| SER | 193 | 9 | 5 | TRUE | 12 | 5 | TRUE | 29 | 1 | TRUE |
| ALA | 194 | 20 | 0 | TRUE | 20 | 0 | TRUE | 26 | 3 | TRUE |
| ARG | 195 | 32 | 1 | TRUE | 32 | 1 | TRUE | 50 | 18 | TRUE |
| LEU | 196 | 29 | 1 | TRUE | 32 | 1 | TRUE | 35 | 1 | TRUE |
| ARG | 197 | 54 | 17 | TRUE | 43 | 8 | TRUE | 67 | 7 | TRUE |
| LEU | 198 | 30 | 4 | TRUE | 28 | 4 | TRUE | 32 | 6 | TRUE |
| ALA | 199 | 36 | 19 | TRUE | 66 | 7 | TRUE | 66 | 7 | TRUE |
| GLY | 200 | 50 | 21 | TRUE | 53 | 22 | TRUE | 54 | 20 | TRUE |
| TRP | 201 | 64 | 3 | FALSE | 64 | 3 | FALSE | 57 | 5 | FALSE |
| HIS | 202 | 65 | 3 | FALSE | 65 | 3 | FALSE | 58 | 5 | FALSE |
| LYS | 203 | 34 | 2 | TRUE | 33 | 1 | TRUE | 36 | 0 | TRUE |
| TYR | 204 | 27 | 10 | TRUE | 37 | 1 | TRUE | 36 | 18 | TRUE |
| LEU | 205 | 26 | 8 | TRUE | 21 | 8 | TRUE | 34 | 23 | TRUE |
| THR | 206 | 29 | 2 | TRUE | 29 | 1 | TRUE | 29 | 3 | TRUE |
| ARG | 207 | 15 | 8 | TRUE | 18 | 1 | TRUE | 23 | 6 | TRUE |
| ASN | 208 | 29 | 5 | TRUE | 29 | 5 | TRUE | 21 | 1 | TRUE |
| GLN | 209 | 26 | 3 | TRUE | 28 | 3 | TRUE | 26 | 4 | TRUE |
| ILE | 210 | 32 | 0 | TRUE | 35 | 1 | TRUE | 32 | 0 | TRUE |
| GLN | 211 | 24 | 0 | FALSE | 24 | 0 | FALSE | 25 | 0 | FALSE |
| PRO | 212 | | | | | | | | | |
| ILE | 213 | 19 | 0 | FALSE | 19 | 0 | FALSE | 20 | 0 | FALSE |
| ALA | 214 | 23 | 0 | FALSE | 23 | 0 | FALSE | 24 | 0 | FALSE |
| GLU | 215 | 13 | 5 | TRUE | 16 | 0 | TRUE | 10 | 5 | TRUE |

| | | | | | | | | | | |
|---|---|---|---|---|---|---|---|---|---|---|
| ARG | 216 | 30 | 0 | TRUE | 31 | 1 | TRUE | 30 | 0 | TRUE |
| GLU | 217 | 14 | 9 | TRUE | 8 | 4 | TRUE | 8 | 4 | TRUE |
| GLY | 218 | 45 | 6 | FALSE | 31 | 3 | FALSE | 27 | 1 | FALSE |
| ASP | 219 | 47 | 6 | FALSE | 34 | 3 | FALSE | 30 | 1 | FALSE |
| TRP | 220 | 39 | 18 | TRUE | 31 | 3 | TRUE | 34 | 2 | TRUE |
| SER | 221 | 21 | 14 | TRUE | 24 | 8 | TRUE | 14 | 6 | TRUE |
| ALA | 222 | 25 | 6 | TRUE | 39 | 4 | TRUE | 68 | 6 | TRUE |
| MET | 223 | 20 | 4 | TRUE | 20 | 0 | TRUE | 19 | 4 | TRUE |
| SER | 224 | 26 | 3 | TRUE | 28 | 0 | TRUE | 24 | 4 | TRUE |
| GLY | 225 | 42 | 3 | TRUE | 38 | 0 | TRUE | 40 | 1 | TRUE |
| PHE | 226 | 51 | 15 | TRUE | 51 | 13 | TRUE | 38 | 3 | TRUE |
| GLN | 227 | 39 | 1 | TRUE | 40 | 2 | TRUE | 53 | 18 | TRUE |
| GLN | 228 | 36 | 1 | TRUE | 36 | 1 | TRUE | 40 | 3 | TRUE |
| THR | 229 | 69 | 6 | TRUE | 69 | 6 | TRUE | 69 | 5 | TRUE |
| MET | 230 | 41 | 4 | TRUE | 54 | 14 | TRUE | 36 | 3 | TRUE |
| GLN | 231 | 32 | 0 | TRUE | 32 | 0 | TRUE | 33 | 2 | TRUE |
| MET | 232 | 38 | 0 | TRUE | 36 | 0 | TRUE | 36 | 1 | TRUE |
| LEU | 233 | 33 | 1 | TRUE | 34 | 0 | TRUE | 31 | 1 | TRUE |
| ASN | 234 | 37 | 1 | TRUE | 38 | 1 | TRUE | 41 | 2 | TRUE |
| GLU | 235 | 28 | 0 | TRUE | 28 | 0 | TRUE | 28 | 0 | TRUE |
| GLY | 236 | 18 | 0 | TRUE | 16 | 0 | TRUE | 18 | 0 | TRUE |
| ILE | 237 | 20 | 0 | TRUE | 20 | 0 | TRUE | 20 | 0 | TRUE |
| VAL | 238 | 64 | 7 | TRUE | 57 | 22 | TRUE | 56 | 20 | TRUE |
| PRO | 239 | | | | | | | | | |
| THR | 240 | 69 | 6 | TRUE | 69 | 6 | TRUE | 69 | 6 | TRUE |
| ALA | 241 | 22 | 0 | TRUE | 22 | 0 | TRUE | 21 | 1 | TRUE |
| MET | 242 | 35 | 2 | TRUE | 35 | 2 | TRUE | 34 | 0 | TRUE |
| LEU | 243 | 34 | 3 | TRUE | 34 | 2 | TRUE | 36 | 2 | TRUE |
| VAL | 244 | 48 | 18 | TRUE | 60 | 13 | TRUE | 32 | 1 | TRUE |
| ALA | 245 | 70 | 6 | TRUE | 58 | 19 | TRUE | 42 | 1 | TRUE |
| ASN | 246 | 30 | 5 | TRUE | 36 | 1 | TRUE | 35 | 6 | TRUE |

| | | | | | | | | | | |
|---|---|---|---|---|---|---|---|---|---|---|
| ASP | 247 | 26 | 4 | TRUE | 28 | 0 | TRUE | 33 | 5 | TRUE |
| GLN | 248 | 53 | 24 | TRUE | 71 | 5 | TRUE | 49 | 19 | TRUE |
| MET | 249 | 71 | 5 | TRUE | 56 | 24 | TRUE | 58 | 13 | TRUE |
| ALA | 250 | 71 | 5 | TRUE | 71 | 5 | TRUE | 69 | 5 | TRUE |
| LEU | 251 | 68 | 5 | TRUE | 69 | 6 | TRUE | 67 | 6 | TRUE |
| GLY | 252 | 69 | 6 | TRUE | 69 | 6 | TRUE | 68 | 6 | TRUE |
| ALA | 253 | 44 | 3 | TRUE | 57 | 21 | TRUE | 41 | 2 | TRUE |
| MET | 254 | 48 | 12 | TRUE | 57 | 20 | TRUE | 69 | 6 | TRUE |
| ARG | 255 | 48 | 18 | TRUE | 51 | 20 | TRUE | 48 | 19 | TRUE |
| ALA | 256 | 37 | 1 | TRUE | 36 | 1 | TRUE | 45 | 16 | TRUE |
| ILE | 257 | 57 | 13 | TRUE | 50 | 17 | TRUE | 52 | 14 | TRUE |
| THR | 258 | 59 | 13 | TRUE | 62 | 12 | TRUE | 68 | 7 | TRUE |
| GLU | 259 | 29 | 1 | TRUE | 28 | 0 | TRUE | 28 | 0 | TRUE |
| SER | 260 | 8 | 4 | TRUE | 8 | 4 | TRUE | 10 | 5 | TRUE |
| GLY | 261 | 56 | 13 | TRUE | 49 | 14 | TRUE | 39 | 2 | TRUE |
| LEU | 262 | 33 | 1 | TRUE | 34 | 1 | TRUE | 32 | 0 | TRUE |
| ARG | 263 | 33 | 2 | TRUE | 35 | 1 | TRUE | 34 | 1 | TRUE |
| VAL | 264 | 29 | 1 | TRUE | 30 | 2 | TRUE | 28 | 2 | TRUE |
| GLY | 265 | 14 | 1 | FALSE | 16 | 1 | FALSE | 16 | 1 | FALSE |
| ALA | 266 | 16 | 1 | FALSE | 17 | 1 | FALSE | 17 | 1 | FALSE |
| ASP | 267 | 16 | 1 | FALSE | 17 | 1 | FALSE | 17 | 1 | FALSE |
| ILE | 268 | 26 | 0 | TRUE | 26 | 2 | TRUE | 25 | 1 | TRUE |
| SER | 269 | 28 | 5 | TRUE | 30 | 0 | TRUE | 29 | 2 | TRUE |
| VAL | 270 | 67 | 1 | FALSE | 66 | 2 | FALSE | 63 | 2 | FALSE |
| VAL | 271 | 65 | 1 | FALSE | 63 | 2 | FALSE | 61 | 2 | FALSE |
| GLY | 272 | 68 | 1 | FALSE | 67 | 2 | FALSE | 65 | 2 | FALSE |
| TYR | 273 | 35 | 1 | TRUE | 38 | 0 | TRUE | 38 | 1 | TRUE |
| ASP | 274 | 47 | 15 | TRUE | 62 | 11 | TRUE | 69 | 6 | TRUE |
| ASP | 275 | 31 | 4 | TRUE | 18 | 1 | TRUE | 46 | 23 | TRUE |
| THR | 276 | 15 | 1 | TRUE | 69 | 6 | TRUE | 39 | 28 | TRUE |
| GLU | 277 | 17 | 1 | TRUE | 9 | 5 | TRUE | 12 | 6 | TRUE |

| | | | | | | | | | | |
|---|---|---|---|---|---|---|---|---|---|---|
| ASP | 278 | 33 | 3 | TRUE | 32 | 1 | TRUE | 23 | 6 | TRUE |
| SER | 279 | 69 | 6 | TRUE | 69 | 5 | TRUE | 59 | 13 | TRUE |
| SER | 280 | 27 | 1 | TRUE | 27 | 1 | TRUE | 26 | 3 | TRUE |
| CYS | 281 | 45 | 14 | TRUE | 68 | 8 | TRUE | 42 | 18 | TRUE |
| TYR | 282 | 30 | 1 | TRUE | 68 | 5 | TRUE | 43 | 21 | TRUE |
| ILE | 283 | 32 | 6 | TRUE | 68 | 6 | TRUE | 41 | 19 | TRUE |
| PRO | 284 | | | | | | | | | |
| PRO | 285 | | | | | | | | | |
| LEU | 286 | 29 | 0 | FALSE | 40 | 2 | FALSE | 38 | 3 | FALSE |
| THR | 287 | 32 | 0 | FALSE | 43 | 2 | FALSE | 41 | 3 | FALSE |
| THR | 288 | 35 | 0 | FALSE | 46 | 2 | FALSE | 43 | 3 | FALSE |
| ILE | 289 | 35 | 1 | FALSE | 66 | 2 | FALSE | 65 | 4 | FALSE |
| LYS | 290 | 37 | 1 | FALSE | 67 | 2 | FALSE | 66 | 4 | FALSE |
| GLN | 291 | 29 | 1 | TRUE | 34 | 0 | TRUE | 38 | 0 | TRUE |
| ASP | 292 | 20 | 1 | TRUE | 21 | 2 | TRUE | 21 | 4 | TRUE |
| PHE | 293 | 17 | 1 | TRUE | 18 | 3 | TRUE | 18 | 4 | TRUE |
| ARG | 294 | 20 | 0 | TRUE | 20 | 0 | TRUE | 20 | 0 | TRUE |
| LEU | 295 | 26 | 0 | TRUE | 26 | 0 | TRUE | 24 | 0 | TRUE |
| LEU | 296 | 31 | 1 | TRUE | 34 | 0 | TRUE | 34 | 0 | TRUE |
| GLY | 297 | 47 | 14 | TRUE | 54 | 13 | TRUE | 49 | 12 | TRUE |
| GLN | 298 | 36 | 1 | TRUE | 38 | 3 | TRUE | 39 | 1 | TRUE |
| THR | 299 | 28 | 0 | TRUE | 45 | 18 | TRUE | 43 | 18 | TRUE |
| SER | 300 | 44 | 2 | TRUE | 45 | 17 | TRUE | 69 | 7 | TRUE |
| VAL | 301 | 38 | 3 | TRUE | 47 | 15 | TRUE | 31 | 7 | TRUE |
| ASP | 302 | 32 | 0 | TRUE | 30 | 3 | TRUE | 36 | 1 | TRUE |
| ARG | 303 | 69 | 6 | TRUE | 69 | 6 | TRUE | 68 | 6 | TRUE |
| LEU | 304 | 70 | 5 | TRUE | 69 | 5 | TRUE | 69 | 6 | TRUE |
| LEU | 305 | 69 | 6 | TRUE | 68 | 6 | TRUE | 68 | 7 | TRUE |
| GLN | 306 | 32 | 0 | TRUE | 33 | 1 | TRUE | 34 | 1 | TRUE |
| LEU | 307 | 16 | 0 | TRUE | 17 | 1 | TRUE | 16 | 1 | TRUE |
| SER | 308 | 9 | 5 | TRUE | 9 | 6 | TRUE | 8 | 4 | TRUE |

| | | | | | | | | | | |
|---|---|---|---|---|---|---|---|---|---|---|
| GLN | 309 | 9 | 5 | TRUE | 17 | 1 | TRUE | 12 | 8 | TRUE |
| GLY | 310 | 18 | 2 | FALSE | 23 | 2 | FALSE | 24 | 2 | FALSE |
| GLN | 311 | 18 | 2 | FALSE | 24 | 2 | FALSE | 25 | 2 | FALSE |
| ALA | 312 | 9 | 4 | TRUE | 8 | 4 | TRUE | 7 | 4 | TRUE |
| VAL | 313 | 3 | 1 | FALSE | 4 | 1 | FALSE | 3 | 1 | FALSE |
| LYS | 314 | 6 | 1 | FALSE | 7 | 1 | FALSE | 6 | 1 | FALSE |
| GLY | 315 | 7 | 1 | FALSE | 9 | 1 | FALSE | 8 | 1 | FALSE |
| ASN | 316 | 9 | 4 | TRUE | 9 | 5 | TRUE | 10 | 4 | TRUE |
| GLN | 317 | 8 | 4 | TRUE | 7 | 4 | TRUE | 8 | 4 | TRUE |
| LEU | 318 | 6 | 3 | TRUE | 7 | 4 | TRUE | 7 | 3 | TRUE |
| LEU | 319 | 5 | 2 | TRUE | 14 | 0 | TRUE | 14 | 0 | TRUE |
| PRO | 320 | | | | | | | | | |
| VAL | 321 | 25 | 0 | FALSE | 27 | 1 | FALSE | 38 | 2 | FALSE |
| SER | 322 | 31 | 0 | FALSE | 33 | 1 | FALSE | 45 | 2 | FALSE |
| LEU | 323 | 28 | 0 | FALSE | 31 | 1 | FALSE | 42 | 2 | FALSE |
| VAL | 324 | | | | | | | | | |
| LYS | 325 | | | | | | | | | |
| ARG | 326 | 13 | 0 | FALSE | 13 | 0 | FALSE | 16 | 0 | FALSE |
| LYS | 327 | 13 | 0 | FALSE | 13 | 0 | FALSE | 15 | 0 | FALSE |
| THR | 328 | 12 | 0 | FALSE | 12 | 0 | FALSE | 15 | 0 | FALSE |
| THR | 329 | 13 | 0 | FALSE | 13 | 0 | FALSE | 15 | 0 | FALSE |
| LEU | 330 | 10 | 0 | FALSE | 10 | 0 | FALSE | 12 | 0 | FALSE |
| ALA | 331 | 11 | 0 | FALSE | 11 | 0 | FALSE | 13 | 0 | FALSE |
| PRO | 332 | | | | | | | | | |
| ASN | 333 | | | | | | | | | |
| THR | 334 | | | | | | | | | |
| GLN | 335 | | | | | | | | | |
| THR | 336 | | | | | | | | | |
| ALA | 337 | | | | | | | | | |
| SER | 338 | | | | | | | | | |
| PRO | 339 | | | | | | | | | |

| | | | | | | | | | | |
|---|---|---|---|---|---|---|---|---|---|---|
| ARG | 340 | | | | | | | | | |
| ALA | 341 | | | | | | | | | |
| LEU | 342 | | | | | | | | | |
| ALA | 343 | | | | | | | | | |
| ASP | 344 | 37 | 4 | FALSE | 37 | 3 | FALSE | 17 | 1 | FALSE |
| SER | 345 | 37 | 4 | FALSE | 37 | 3 | FALSE | 17 | 1 | FALSE |
| LEU | 346 | 35 | 4 | FALSE | 35 | 3 | FALSE | 15 | 1 | FALSE |
| MET | 347 | 14 | 2 | FALSE | 18 | 2 | FALSE | 22 | 2 | FALSE |
| GLN | 348 | 16 | 2 | FALSE | 20 | 2 | FALSE | 25 | 2 | FALSE |
| LEU | 349 | 13 | 2 | FALSE | 17 | 2 | FALSE | 21 | 2 | FALSE |
| ALA | 350 | 36 | 7 | FALSE | 29 | 8 | FALSE | 47 | 6 | FALSE |
| ARG | 351 | 37 | 7 | FALSE | 30 | 8 | FALSE | 48 | 6 | FALSE |
| GLN | 352 | 63 | 12 | TRUE | 52 | 20 | TRUE | 49 | 14 | TRUE |
| VAL | 353 | 16 | 1 | TRUE | 26 | 22 | TRUE | 25 | 18 | TRUE |
| SER | 354 | 53 | 22 | TRUE | 44 | 22 | TRUE | 54 | 20 | TRUE |
| ARG | 355 | 41 | 1 | FALSE | 40 | 2 | FALSE | 41 | 1 | FALSE |
| LEU | 356 | 37 | 1 | FALSE | 35 | 2 | FALSE | 37 | 1 | FALSE |
| GLU | 357 | 37 | 1 | FALSE | 36 | 2 | FALSE | 37 | 1 | FALSE |
| SER | 358 | 40 | 1 | FALSE | 39 | 2 | FALSE | 40 | 1 | FALSE |
| GLY | 359 | 41 | 1 | FALSE | 39 | 2 | FALSE | 40 | 1 | FALSE |
| GLN | 360 | 25 | 2 | TRUE | 23 | 2 | TRUE | 33 | 22 | TRUE |

#### Table S13. Plasmid sequences used in this study

| Protein | DNA sequence |
|---|---|
| *E. coli* DHFR with N. term 6X-His-TEV site tag | TTCTTAGAAAAACTCATCGAGCATCAAATGAAACTGCAATTTATTCATATCAGGATTATCAATA<br>CCATATTTTTGAAAAAGCCGTTTCTGTAATGAAGGAGAAAACTCACCGAGGCAGTTCCATAGGA<br>TGGCAAGATCCTGGTATCGGTCTGCGATTCCGACTCGTCCAACATCAATACAACCTATTAATTT<br>CCCCTCGTCAAAAATAAGGTTATCAAGTGAGAAATCACCATGAGTGACGACTGAATCCGGTGAG<br>AATGGCAAAAGCTTATGCATTTCTTTCCAGACTTGTTCAACAGGCCAGCCATTACGCTCGTCAT<br>CAAAATCACTCGCATCAACCAAACCGTTATTCATTCGTGATTGCGCCTGAGCGAGACGAAATAC<br>GCGATCGCTGTTAAAAGGACAATTACAAACAGGAATCGAATGCAACCGGCGCAGGAACACTGCC<br>AGCGCATCAACAATATTTTCACCTGAATCAGGATATTCTTCTAATACCTGGAATGCTGTTTTCC<br>CGGGGATCGCAGTGGTGAGTAACCATGCATCATCAGGAGTACGGATAAAATGCTTGATGGTCGG<br>AAGAGGCATAAATTCCGTCAGCCAGTTTAGTCTGACCATCTCATCTGTAACATCATTGGCAACG<br>CTACCTTTGCCATGTTTCAGAAACAACTCTGGCGCATCGGGCTTCCCATACAATCGATAGATTG<br>TCGCACCTGATTGCCCGACATTATCGCGAGCCCATTTATACCCATATAAATCAGCATCCATGTT<br>GGAATTTAATCGCGGCCTCGAGCAAGACGTTTCCCGTTGAATATGGCTCATAACACCCCTTGTA<br>TTACTGTTTATGTAAGCAGACAGTTTTATTGTTCATGACCAAAATCCCTTAACGTGAGTTTTCG<br>TTCCACTGAGCGTCAGACCCCGTAGAAAAGATCAAAGGATCTTCTTGAGATCCTTTTTTTCTGC<br>GCGTAATCTGCTGCTTGCAAACAAAAAAACCACCGCTACCAGCGGTGGTTTGTTTGCCGGATCA<br>AGAGCTACCAACTCTTTTTCCGAAGGTAACTGGCTTCAGCAGAGCGCAGATACCAAATACTGTC<br>CTTCTAGTGTAGCCGTAGTTAGGCCACCACTTCAAGAACTCTGTAGCACCGCCTACATACCTCG<br>CTCTGCTAATCCTGTTACCAGTGGCTGCTGCCAGTGGCGATAAGTCGTGTCTTACCGGGTTGGA<br>CTCAAGACGATAGTTACCGGATAAGGCGCAGCGGTCGGGCTGAACGGGGGGTTCGTGCACACAG<br>CCCAGCTTGGAGCGAACGACCTACACCGAACTGAGATACCTACAGCGTGAGCTATGAGAAAGCG<br>CCACGCTTCCCGAAGGGAGAAAGGCGGACAGGTATCCGGTAAGCGGCAGGGTCGGAACAGGAGA<br>GCGCACGAGGGAGCTTCCAGGGGGAAACGCCTGGTATCTTTATAGTCCTGTCGGGTTTCGCCAC<br>CTCTGACTTGAGCGTCGATTTTTGTGATGCTCGTCAGGGGGGCGGAGCCTATGGAAAAACGCCA<br>GCAACGCGGCCTTTTTACGGTTCCTGGCCTTTTGCTGGCCTTTTGCTCACATGTTCTTTCCTGC<br>GTTATCCCCTGATTCTGTGGATAACCGTATTACCGCCTTTGAGTGAGCTGATACCGCTCGCCGC<br>AGCCGAACGACCGAGCGCAGCGAGTCAGTGAGCGAGGAAGCGGAAGAGCGCCTGATGCGGTATT<br>TTCTCCTTACGCATCTGTGCGGTATTTCACACCGCATATATGGTGCACTCTCAGTACAATCTGC<br>TCTGATGCCGCATAGTTAAGCCAGTATACACTCCGCTATCGCTACGTGACTGGGTCATGGCTGC<br>GCCCCGACACCCGCCAACACCCGCTGACGCGCCCTGACGGGCTTGTCTGCTCCCGGCATCCGCT<br>TACAGACAAGCTGTGACCGTCTCCGGGAGCTGCATGTGTCAGAGGTTTTCACCGTCATCACCGA<br>AACGCGCGAGGCAGCTGCGGTAAAGCTCATCAGCGTGGTCGTGAAGCGATTCACAGATGTCTGC<br>CTGTTCATCCGCGTCCAGCTCGTTGAGTTTCTCCAGAAGCGTTAATGTCTGGCTTCTGATAAAG<br>CGGGCCATGTTAAGGGCGGTTTTTTCCTGTTTGGTCACTGATGCCTCCGTGTAAGGGGGATTTC<br>TGTTCATGGGGGTAATGATACCGATGAAACGAGAGAGGATGCTCACGATACGGGTTACTGATGA<br>TGAACATGCCCGGTTACTGGAACGTTGTGAGGGTAAACAACTGGCGGTATGGATGCGGCGGGAC<br>CAGAGAAAAATCACTCAGGGTCAATGCCAGCGCTTCGTTAATACAGATGTAGGTGTTCCACAGG<br>GTAGCCAGCAGCATCCTGCGATGCAGATCCGGAACATAATGGTGCAGGGCGCTGACTTCCGCGT<br>TTCCAGACTTTACGAAACACGGAAACCGAAGACCATTCATGTTGTTGCTCAGGTCGCAGACGTT<br>TTGCAGCAGCAGTCGCTTCACGTTCGCTCGCGTATCGGTGATTCATTCTGCTAACCAGTAAGGC<br>AACCCCGCCAGCCTAGCCGGGTCCTCAACGACAGGAGCACGATCATGCGCACCCGTGGCCAGGA<br>CCCAACGCTGCCCGAGATGCGCCGCGTGCGGCTGCTGGAGATGGCGGACGCGATGGATATGTTC<br>TGCCAAGGGTTGGTTTGCGCATTCACAGTTCTCCGCAAGAATTGATTGGCTCCAATTCTTGGAG<br>TGGTGAATCCGTTAGCGAGGTGCCGCCGGCTTCCATTCAGGTCGAGGTGGCCCGGCTCCATGCA<br>CCGCGACGCAACGCGGGGAGGCAGACAAGGTATAGGGCGGCGCCTACAATCCATGCCAACCCGT<br>TCCATGTGCTCGCCGAGGCGGCATAAATCGCCGTGACGATCAGCGGTCCAGTGATCGAAGTTAG<br>GCTGGTAAGAGCCGCGAGCGATCCTTGAAGCTGTCCCTGATGGTCGTCATCTACCTGCCTGGAC<br>AGCATGGCCTGCAACGCGGGCATCCCGATGCCGCCGGAAGCGAGAAGAATCATAATGGGGAAGG<br>CCATCCAGCCTCGCGTCGCGAACGCCAGCAAGACGTAGCCCAGCGCGTCGGCCGCCATGCCGGC<br>GATAATGGCCTGCTTCTCGCCGAAACGTTTGGTGGCGGGACCAGTGACGAAGGCTTGAGCGAGG |

| | |
|---|---|
| | GCGTGCAAGATTCCGAATACCGCAAGCGACAGGCCGATCATCGTCGCGCTCCAGCGAAAGCGGT<br>CCTCGCCGAAAATGACCCAGAGCGCTGCCGGCACCTGTCCTACGAGTTGCATGATAAAGAAGAC<br>AGTCATAAGTGCGGCGACGATAGTCATGCCCCGCGCCCACCGGAAGGAGCTGACTGGGTTGAAG<br>GCTCTCAAGGGCATCGGTCGACGCTCTCCCTTATGCGACTCCTGCATTAGGAAGCAGCCCAGTA<br>GTAGGTTGAGGCCGTTGAGCACCGCCGCCGCAAGGAATGGTGCATGCAAGGAGATGGCGCCCAA<br>CAGTCCCCCGGCCACGGGGCCTGCCACCATACCCACGCCGAAACAAGCGCTCATGAGCCCGAAG<br>TGGCGAGCCCGATCTTCCCCATCGGTGATGTCGGCGATATAGGCGCCAGCAACCGCACCTGTGG<br>CGCCGGTGATGCCGGCCACGATGCGTCCGGCGTAGAGGATCGAGATCTCGATCCCGCGAAATTA<br>ATACGACTCACTATAGGCCCCTCTAGAAATAATTTTGTTTAACTTTAAGAAGGAGATATACCAT<br>GACCGGTCATCACCATCACCATCACGAAAACTTATATTTCCAATCTATCAGCCTGATTGCTGCA<br>TTAGCCGTGGACCGTGTGATTGGCATGGAGAATGCAATGCCGTGGAACTTACCGGCGGACCTTG<br>CATGGTTCAAGCGCAACACACTGGATAAGCCGGTAATCATGGGTCGCCATACTTGGGAATCCAT<br>CGGCCGTCCGTTGCCCGGCCGTAAGAACATTATCTTGTCTTCTCAGCCTGGGACTGACGACCGT<br>GTAACATGGGTCAAAAGTGTAGATGAAGCTATCGCGGCTTGTGGGGATGTTCCTGAGATCATGG<br>TAATCGGCGGGGGCCGTGTGTACGAGCAGTTCCTGCCCAAAGCGCAGAAACTTTATTTGACTCA<br>CATTGATGCAGAGGTGGAAGGGGATACGCACTTTCCAGATTACGAGCCAGACGACTGGGAATCA<br>GTCTTCTCCGAATTTCACGATGCGGACGCCCAAAATAGCCACTCTTATTGCTTTGAGATTTTGG<br>AACGCCGTTAAGAGCTCCGTCGACAAGCTTGCGGCCGCACTCGAGCACCACCACCACCACCACT<br>GAGATCCGGCTGCTAACAAAGCCCGAAAGGAAGCTGAGTTGGCTGCTGCCACCGCTGAGCAATA<br>ACTAGCATAACCCCTTGGGGCCTCTAAACGGGTCTTGAGGGGTTTTTTGCTGAAAGGAGGAACT<br>ATATCCGGATATCCACAGGACGGGTGTGGTCGCCATGATCGCGTAGTCGATAGTGGCTCCAAGT<br>AGCGAAGCGAGCAGGACTGGGCGGCGGCCAAAGCGGTCGGACAGTGCTCCGAGAACGGGTGCGC<br>ATAGAAATTGCATCAACGCATATAGCGCTAGCAGCACGCCATAGTGACTGGCGATGCTGTCGGA<br>ATGGACGATATCCCGCAAGAGGCCCGGCAGTACCGGCATAACCAAGCCTATGCCTACAGCATCC<br>AGGGTGACGGTGCCGAGGATGACGATGAGCGCATTGTTAGATTTCATACACGGTGCCTGACTGC<br>GTTAGCAATTTAACTGTGATAAACTACCGCATTAAAGCTTATCGATGATAAGCTGTCAAACATG<br>AGAA |
| *E. coli lac* repressor with N-term 6X-His-TEV site tag | TTCTTAGAAAAACTCATCGAGCATCAAATGAAACTGCAATTTATTCATATCAGGATTATCAATA<br>CCATATTTTTGAAAAAGCCGTTTCTGTAATGAAGGAGAAAACTCACCGAGGCAGTTCCATAGGA<br>TGGCAAGATCCTGGTATCGGTCTGCGATTCCGACTCGTCCAACATCAATACAACCTATTAATTT<br>CCCCTCGTCAAAAATAAGGTTATCAAGTGAGAAATCACCATGAGTGACGACTGAATCCGGTGAG<br>AATGGCAAAAGCTTATGCATTTCTTTCCAGACTTGTTCAACAGGCCAGCCATTACGCTCGTCAT<br>CAAAATCACTCGCATCAACCAAACCGTTATTCATTCGTGATTGCGCCTGAGCGAGACGAAATAC<br>GCGATCGCTGTTAAAAGGACAATTACAAACAGGAATCGAATGCAACCGGCGCAGGAACACTGCC<br>AGCGCATCAACAATATTTTCACCTGAATCAGGATATTCTTCTAATACCTGGAATGCTGTTTTCC<br>CGGGGATCGCAGTGGTGAGTAACCATGCATCATCAGGAGTACGGATAAAATGCTTGATGGTCGG<br>AAGAGGCATAAATTCCGTCAGCCAGTTTAGTCTGACCATCTCATCTGTAACATCATTGGCAACG<br>CTACCTTTGCCATGTTTCAGAAACAACTCTGGCGCATCGGGCTTCCCATACAATCGATAGATTG<br>TCGCACCTGATTGCCCGACATTATCGCGAGCCCATTTATACCCATATAAATCAGCATCCATGTT<br>GGAATTTAATCGCGGCCTCGAGCAAGACGTTTCCCGTTGAATATGGCTCATAACACCCCTTGTA<br>TTACTGTTTATGTAAGCAGACAGTTTTATTGTTCATGACCAAAATCCCTTAACGTGAGTTTTCG<br>TTCCACTGAGCGTCAGACCCCGTAGAAAAGATCAAAGGATCTTCTTGAGATCCTTTTTTTCTGC<br>GCGTAATCTGCTGCTTGCAAACAAAAAAACCACCGCTACCAGCGGTGGTTTGTTTGCCGGATCA<br>AGAGCTACCAACTCTTTTTCCGAAGGTAACTGGCTTCAGCAGAGCGCAGATACCAAATACTGTC<br>CTTCTAGTGTAGCCGTAGTTAGGCCACCACTTCAAGAACTCTGTAGCACCGCCTACATACCTCG<br>CTCTGCTAATCCTGTTACCAGTGGCTGCTGCCAGTGGCGATAAGTCGTGTCTTACCGGGTTGGA<br>CTCAAGACGATAGTTACCGGATAAGGCGCAGCGGTCGGGCTGAACGGGGGGTTCGTGCACACAG<br>CCCAGCTTGGAGCGAACGACCTACACCGAACTGAGATACCTACAGCGTGAGCTATGAGAAAGCG<br>CCACGCTTCCCGAAGGGAGAAAGGCGGACAGGTATCCGGTAAGCGGCAGGGTCGGAACAGGAGA<br>GCGCACGAGGGAGCTTCCAGGGGGAAACGCCTGGTATCTTTATAGTCCTGTCGGGTTTCGCCAC<br>CTCTGACTTGAGCGTCGATTTTTGTGATGCTCGTCAGGGGGGCGGAGCCTATGGAAAAACGCCA<br>GCAACGCGGCCTTTTTACGGTTCCTGGCCTTTTGCTGGCCTTTTGCTCACATGTTCTTTCCTGC<br>GTTATCCCCTGATTCTGTGGATAACCGTATTACCGCCTTTGAGTGAGCTGATACCGCTCGCCGC<br>AGCCGAACGACCGAGCGCAGCGAGTCAGTGAGCGAGGAAGCGGAAGAGCGCCTGATGCGGTATT<br>TTCTCCTTACGCATCTGTGCGGTATTTCACACCGCATATATGGTGCACTCTCAGTACAATCTGC |

| | |
|---|---|
| | TCTGATGCCGCATAGTTAAGCCAGTATACACTCCGCTATCGCTACGTGACTGGGTCATGGCTGC<br>GCCCCGACACCCGCCAACACCCGCTGACGCGCCCTGACGGGCTTGTCTGCTCCCGGCATCCGCT<br>TACAGACAAGCTGTGACCGTCTCCGGGAGCTGCATGTGTCAGAGGTTTTCACCGTCATCACCGA<br>AACGCGCGAGGCAGCTGCGGTAAAGCTCATCAGCGTGGTCGTGAAGCGATTCACAGATGTCTGC<br>CTGTTCATCCGCGTCCAGCTCGTTGAGTTTCTCCAGAAGCGTTAATGTCTGGCTTCTGATAAAG<br>CGGGCCATGTTAAGGGCGGTTTTTTCCTGTTTGGTCACTGATGCCTCCGTGTAAGGGGGATTTC<br>TGTTCATGGGGGTAATGATACCGATGAAACGAGAGAGGATGCTCACGATACGGGTTACTGATGA<br>TGAACATGCCCGGTTACTGGAACGTTGTGAGGGTAAACAACTGGCGGTATGGATGCGGCGGGAC<br>CAGAGAAAAATCACTCAGGGTCAATGCCAGCGCTTCGTTAATACAGATGTAGGTGTTCCACAGG<br>GTAGCCAGCAGCATCCTGCGATGCAGATCCGGAACATAATGGTGCAGGGCGCTGACTTCCGCGT<br>TTCCAGACTTTACGAAACACGGAAACCGAAGACCATTCATGTTGTTGCTCAGGTCGCAGACGTT<br>TTGCAGCAGCAGTCGCTTCACGTTCGCTCGCGTATCGGTGATTCATTCTGCTAACCAGTAAGGC<br>AACCCCGCCAGCCTAGCCGGGTCCTCAACGACAGGAGCACGATCATGCGCACCCGTGGCCAGGA<br>CCCAACGCTGCCCGAGATGCGCCGCGTGCGGCTGCTGGAGATGGCGGACGCGATGGATATGTTC<br>TGCCAAGGGTTGGTTTGCGCATTCACAGTTCTCCGCAAGAATTGATTGGCTCCAATTCTTGGAG<br>TGGTGAATCCGTTAGCGAGGTGCCGCCGGCTTCCATTCAGGTCGAGGTGGCCCGGCTCCATGCA<br>CCGCGACGCAACGCGGGGAGGCAGACAAGGTATAGGGCGGCGCCTACAATCCATGCCAACCCGT<br>TCCATGTGCTCGCCGAGGCGGCATAAATCGCCGTGACGATCAGCGGTCCAGTGATCGAAGTTAG<br>GCTGGTAAGAGCCGCGAGCGATCCTTGAAGCTGTCCCTGATGGTCGTCATCTACCTGCCTGGAC<br>AGCATGGCCTGCAACGCGGGCATCCCGATGCCGCCGGAAGCGAGAAGAATCATAATGGGGAAGG<br>CCATCCAGCCTCGCGTCGCGAACGCCAGCAAGACGTAGCCCAGCGCGTCGGCCGCCATGCCGGC<br>GATAATGGCCTGCTTCTCGCCGAAACGTTTGGTGGCGGGACCAGTGACGAAGGCTTGAGCGAGG<br>GCGTGCAAGATTCCGAATACCGCAAGCGACAGGCCGATCATCGTCGCGCTCCAGCGAAAGCGGT<br>CCTCGCCGAAAATGACCCAGAGCGCTGCCGGCACCTGTCCTACGAGTTGCATGATAAAGAAGAC<br>AGTCATAAGTGCGGCGACGATAGTCATGCCCCGCGCCCACCGGAAGGAGCTGACTGGGTTGAAG<br>GCTCTCAAGGGCATCGGTCGACGCTCTCCCTTATGCGACTCCTGCATTAGGAAGCAGCCCAGTA<br>GTAGGTTGAGGCCGTTGAGCACCGCCGCCGCAAGGAATGGTGCATGCAAGGAGATGGCGCCCAA<br>CAGTCCCCCGGCCACGGGGCCTGCCACCATACCCACGCCGAAACAAGCGCTCATGAGCCCGAAG<br>TGGCGAGCCCGATCTTCCCCATCGGTGATGTCGGCGATATAGGCGCCAGCAACCGCACCTGTGG<br>CGCCGGTGATGCCGGCCACGATGCGTCCGGCGTAGAGGATCGAGATCTCGATCCCGCGAAATTA<br>ATACGACTCACTATAGGCCCCTCTAGAAATAATTTTGTTTAACTTTAAGAAGGAGATATACCAT<br>GCATCACCATCACCATCACGAAAACTTATATTTCCAATCTAAACCAGTAACGTTATACGATGTC<br>GCAGAGTATGCCGGTGTCTCTTATCAGACCGTTTCCCGCGTGGTGAACCAGGCCAGCCACGTTT<br>CTGCGAAAACGCGGGAAAAAGTGGAAGCGGCGATGGCGGAGCTGAATTACATTCCCAACCGCGT<br>GGCACAACAACTGGCGGGCAAACAGTCGTTGCTGATTGGCGTTGCCACCTCCAGTCTGGCCCTG<br>CACGCGCCGTCGCAAATTGTCGCGGCGATTAAATCTCGCGCCGATCAACTGGGTGCCAGCGTGG<br>TGGTGTCGATGGTAGAACGAAGCGGCGTCGAAGCCTGTAAAGCGGCGGTGCACAATCTTCTCGC<br>GCAACGCGTCAGTGGGCTGATCATTAACTATCCGCTGGATGACCAGGATGCCATTGCTGTGGAA<br>GCTGCCTGCACTAATGTTCCGGCGTTATTTCTTGATGTCTCTGACCAGACACCCATCAACAGTA<br>TTATTTTCTCCCATGAAGACGGTACGCGACTGGGCGTGGAGCATCTGGTCGCATTGGGTCACCA<br>GCAAATCGCGCTGTTAGCGGGCCCATTAAGTTCTGTCTCGGCGCGTCTGCGTCTGGCTGGCTGG<br>CATAAATATCTCACTCGCAATCAAATTCAGCCGATAGCGGAACGGGAAGGCGACTGGAGTGCCA<br>TGTCCGGTTTTCAACAAACCATGCAAATGCTGAATGAGGGCATCGTTCCCACTGCGATGCTGGT<br>TGCCAACGATCAGATGGCGCTGGGCGCAATGCGCGCCATTACCGAGTCCGGGCTGCGCGTTGGT<br>GCGGATATCTCGGTAGTGGGATACGACGATACCGAAGACAGCTCATGTTATATCCCGCCGTCAA<br>CCACCATCAAACAGGATTTTCGCCTGCTGGGGCAAACCAGCGTGGACCGCTTGCTGCAACTCTC<br>TCAGGGCCAGGCGGTGAAGGGCAATCAGCTGTTGCCCGTCTCACTGGTGAAAAGAAAAACCACC<br>CTGGCGCCCTAAGAGCTCCGTCGACAAGCTTGCGGCCGCACTCGAGCACCACCACCACCACCAC<br>TGAGATCCGGCTGCTAACAAAGCCCGAAAGGAAGCTGAGTTGGCTGCTGCCACCGCTGAGCAAT<br>AACTAGCATAACCCCTTGGGGCCTCTAAACGGGTCTTGAGGGGTTTTTTGCTGAAAGGAGGAAC<br>TATATCCGGATATCCACAGGACGGGTGTGGTCGCCATGATCGCGTAGTCGATAGTGGCTCCAAG<br>TAGCGAAGCGAGCAGGACTGGGCGGCGGCCAAAGCGGTCGGACAGTGCTCCGAGAACGGGTGCG<br>CATAGAAATTGCATCAACGCATATAGCGCTAGCAGCACGCCATAGTGACTGGCGATGCTGTCGG<br>AATGGACGATATCCCGCAAGAGGCCCGGCAGTACCGGCATAACCAAGCCTATGCCTACAGCATC<br>CAGGGTGACGGTGCCGAGGATGACGATGAGCGCATTGTTAGATTTCATACACGGTGCCTGACTG<br>CGTTAGCAATTTAACTGTGATAAACTACCGCATTAAAGCTTATCGATGATAAGCTGTCAAACAT |

| | |
|---|---|
| | GAGAA |
| *Homo sapiens* DHFR with N-term 6X-His-TEV site tag | GCGAACGCCAGCAAGACGTAGCCCAGCGCGTCGGCCGCCATGCCGGCGATAATGGCCTGCTTCT<br>CGCCGAAACGTTTGGTGGCGGGACCAGTGACGAAGGCTTGAGCGAGGGCGTGCAAGATTCCGAA<br>TACCGCAAGCGACAGGCCGATCATCGTCGCGCTCCAGCGAAAGCGGTCCTCGCCGAAAATGACC<br>CAGAGCGCTGCCGGCACCTGTCCTACGAGTTGCATGATAAAGAAGACAGTCATAAGTGCGGCGA<br>CGATAGTCATGCCCCGCGCCCACCGGAAGGAGCTGACTGGGTTGAAGGCTCTCAAGGGCATCGG<br>TCGACGCTCTCCCTTATGCGACTCCTGCATTAGGAAGCAGCCCAGTAGTAGGTTGAGGCCGTTG<br>AGCACCGCCGCCGCAAGGAATGGTGCATGCAAGGAGATGGCGCCCAACAGTCCCCCGGCCACGG<br>GGCCTGCCACCATACCCACGCCGAAACAAGCGCTCATGAGCCCGAAGTGGCGAGCCCGATCTTC<br>CCCATCGGTGATGTCGGCGATATAGGCGCCAGCAACCGCACCTGTGGCGCCGGTGATGCCGGCC<br>ACGATGCGTCCGGCGTAGAGGATCGAGATCTCGATCCCGCGAAATTAATACGACTCACTATAGG<br>CCCCTCTAGAAATAATTTTGTTTAACTTTAAGAAGGAGATATACCATGGGTTCGTCACACCACC<br>ATCATCACCACTCTTCTGGACTTGTACCACGTGGAAGTCATATGGTAGGATCTCTGAATTGTAT<br>TGTTGCAGTGTCTCAGAATATGGGCATCGGAAAGAACGGGGACTTGCCGTGGCCTCCATTACGC<br>AACGAATTCCGCTACTTTCAACGTATGACGACCACCTCTAGTGTGGAGGGGAAGCAAAATTTGG<br>TGATTATGGGTAAAAAAACCTGGTTCAGTATTCCAGAAAAAAACCGCCCTTTAAAGGGCCGCAT<br>CAACCTTGTGCTTTCACGTGAACTTAAGGAGCCACCTCAGGGCGCGCATTTTCTTTCTCGCTCC<br>CTTGACGACGCACTGAAGTTGACTGAACAGCCAGAACTGGCTAATAAAGTTGACATGGTTTGGA<br>TCGTCGGGGGGTCGTCAGTGTATAAGGAGGCCATGAACCATCCAGGACATCTGAAATTGTTTGT<br>TACTCGCATTATGCAGGATTTCGAGAGCGATACATTTTTTCCTGAGATCGACCTGGAAAAATAT<br>AAGCTGCTGCCGGAATACCCAGGGGTATTAAGTGACGTTCAAGAGGAGAAAGGCATCAAATACA<br>AATTTGAAGTCTATGAGAAGAATGATTAAGAGCTCCGTCGACAAGCTTGCGGCCGCACTCGAGC<br>ACCACCACCACCACCACTGAGATCCGGCTGCTAACAAAGCCCGAAAGGAAGCTGAGTTGGCTGC<br>TGCCACCGCTGAGCAATAACTAGCATAACCCCTTGGGGCCTCTAAACGGGTCTTGAGGGGTTTT<br>TTGCTGAAAGGAGGAACTATATCCGGATATCCACAGGACGGGTGTGGTCGCCATGATCGCGTAG<br>TCGATAGTGGCTCCAAGTAGCGAAGCGAGCAGGACTGGGCGGCGGCCAAAGCGGTCGGACAGTG<br>CTCCGAGAACGGGTGCGCATAGAAATTGCATCAACGCATATAGCGCTAGCAGCACGCCATAGTG<br>ACTGGCGATGCTGTCGGAATGGACGATATCCCGCAAGAGGCCCGGCAGTACCGGCATAACCAAG<br>CCTATGCCTACAGCATCCAGGGTGACGGTGCCGAGGATGACGATGAGCGCATTGTTAGATTTCA<br>TACACGGTGCCTGACTGCGTTAGCAATTTAACTGTGATAAACTACCGCATTAAAGCTTATCGAT<br>GATAAGCTGTCAAACATGAGAATTCTTAGAAAAACTCATCGAGCATCAAATGAAACTGCAATTT<br>ATTCATATCAGGATTATCAATACCATATTTTTGAAAAAGCCGTTTCTGTAATGAAGGAGAAAAC<br>TCACCGAGGCAGTTCCATAGGATGGCAAGATCCTGGTATCGGTCTGCGATTCCGACTCGTCCAA<br>CATCAATACAACCTATTAATTTCCCCTCGTCAAAAATAAGGTTATCAAGTGAGAAATCACCATG<br>AGTGACGACTGAATCCGGTGAGAATGGCAAAAGCTTATGCATTTCTTTCCAGACTTGTTCAACA<br>GGCCAGCCATTACGCTCGTCATCAAAATCACTCGCATCAACCAAACCGTTATTCATTCGTGATT<br>GCGCCTGAGCGAGACGAAATACGCGATCGCTGTTAAAAGGACAATTACAAACAGGAATCGAATG<br>CAACCGGCGCAGGAACACTGCCAGCGCATCAACAATATTTTCACCTGAATCAGGATATTCTTCT<br>AATACCTGGAATGCTGTTTTCCCGGGGATCGCAGTGGTGAGTAACCATGCATCATCAGGAGTAC<br>GGATAAAATGCTTGATGGTCGGAAGAGGCATAAATTCCGTCAGCCAGTTTAGTCTGACCATCTC<br>ATCTGTAACATCATTGGCAACGCTACCTTTGCCATGTTTCAGAAACAACTCTGGCGCATCGGGC<br>TTCCCATACAATCGATAGATTGTCGCACCTGATTGCCCGACATTATCGCGAGCCCATTTATACC<br>CATATAAATCAGCATCCATGTTGGAATTTAATCGCGGCCTCGAGCAAGACGTTTCCCGTTGAAT<br>ATGGCTCATAACACCCCTTGTATTACTGTTTATGTAAGCAGACAGTTTTATTGTTCATGACCAA<br>AATCCCTTAACGTGAGTTTTCGTTCCACTGAGCGTCAGACCCCGTAGAAAAGATCAAAGGATCT<br>TCTTGAGATCCTTTTTTTCTGCGCGTAATCTGCTGCTTGCAAACAAAAAAACCACCGCTACCAG<br>CGGTGGTTTGTTTGCCGGATCAAGAGCTACCAACTCTTTTTCCGAAGGTAACTGGCTTCAGCAG<br>AGCGCAGATACCAAATACTGTCCTTCTAGTGTAGCCGTAGTTAGGCCACCACTTCAAGAACTCT<br>GTAGCACCGCCTACATACCTCGCTCTGCTAATCCTGTTACCAGTGGCTGCTGCCAGTGGCGATA<br>AGTCGTGTCTTACCGGGTTGGACTCAAGACGATAGTTACCGGATAAGGCGCAGCGGTCGGGCTG<br>AACGGGGGGTTCGTGCACACAGCCCAGCTTGGAGCGAACGACCTACACCGAACTGAGATACCTA<br>CAGCGTGAGCTATGAGAAAGCGCCACGCTTCCCGAAGGGAGAAAGGCGGACAGGTATCCGGTAA<br>GCGGCAGGGTCGGAACAGGAGAGCGCACGAGGGAGCTTCCAGGGGGAAACGCCTGGTATCTTTA<br>TAGTCCTGTCGGGTTTCGCCACCTCTGACTTGAGCGTCGATTTTTGTGATGCTCGTCAGGGGGG<br>CGGAGCCTATGGAAAAACGCCAGCAACGCGGCCTTTTTACGGTTCCTGGCCTTTTGCTGGCCTT |

| | |
|---|---|
| | TTGCTCACATGTTCTTTCCTGCGTTATCCCCTGATTCTGTGGATAACCGTATTACCGCCTTTGA<br>GTGAGCTGATACCGCTCGCCGCAGCCGAACGACCGAGCGCAGCGAGTCAGTGAGCGAGGAAGCG<br>GAAGAGCGCCTGATGCGGTATTTTCTCCTTACGCATCTGTGCGGTATTTCACACCGCATATATG<br>GTGCACTCTCAGTACAATCTGCTCTGATGCCGCATAGTTAAGCCAGTATACACTCCGCTATCGC<br>TACGTGACTGGGTCATGGCTGCGCCCCGACACCCGCCAACACCCGCTGACGCGCCCTGACGGGC<br>TTGTCTGCTCCCGGCATCCGCTTACAGACAAGCTGTGACCGTCTCCGGGAGCTGCATGTGTCAG<br>AGGTTTTCACCGTCATCACCGAAACGCGCGAGGCAGCTGCGGTAAAGCTCATCAGCGTGGTCGT<br>GAAGCGATTCACAGATGTCTGCCTGTTCATCCGCGTCCAGCTCGTTGAGTTTCTCCAGAAGCGT<br>TAATGTCTGGCTTCTGATAAAGCGGGCCATGTTAAGGGCGGTTTTTTCCTGTTTGGTCACTGAT<br>GCCTCCGTGTAAGGGGGATTTCTGTTCATGGGGGTAATGATACCGATGAAACGAGAGAGGATGC<br>TCACGATACGGGTTACTGATGATGAACATGCCCGGTTACTGGAACGTTGTGAGGGTAAACAACT<br>GGCGGTATGGATGCGGCGGGACCAGAGAAAAATCACTCAGGGTCAATGCCAGCGCTTCGTTAAT<br>ACAGATGTAGGTGTTCCACAGGGTAGCCAGCAGCATCCTGCGATGCAGATCCGGAACATAATGG<br>TGCAGGGCGCTGACTTCCGCGTTTCCAGACTTTACGAAACACGGAAACCGAAGACCATTCATGT<br>TGTTGCTCAGGTCGCAGACGTTTTGCAGCAGCAGTCGCTTCACGTTCGCTCGCGTATCGGTGAT<br>TCATTCTGCTAACCAGTAAGGCAACCCCGCCAGCCTAGCCGGGTCCTCAACGACAGGAGCACGA<br>TCATGCGCACCCGTGGCCAGGACCCAACGCTGCCCGAGATGCGCCGCGTGCGGCTGCTGGAGAT<br>GGCGGACGCGATGGATATGTTCTGCCAAGGGTTGGTTTGCGCATTCACAGTTCTCCGCAAGAAT<br>TGATTGGCTCCAATTCTTGGAGTGGTGAATCCGTTAGCGAGGTGCCGCCGGCTTCCATTCAGGT<br>CGAGGTGGCCCGGCTCCATGCACCGCGACGCAACGCGGGGAGGCAGACAAGGTATAGGGCGGCG<br>CCTACAATCCATGCCAACCCGTTCCATGTGCTCGCCGAGGCGGCATAAATCGCCGTGACGATCA<br>GCGGTCCAGTGATCGAAGTTAGGCTGGTAAGAGCCGCGAGCGATCCTTGAAGCTGTCCCTGATG<br>GTCGTCATCTACCTGCCTGGACAGCATGGCCTGCAACGCGGGCATCCCGATGCCGCCGGAAGCG<br>AGAAGAATCATAATGGGGAAGGCCATCCAGCCTCGCGTC |

### 5. Supplemental methods

**Protein expression and purification**

*ecDHFR*

We introduced an *E. coli* DHFR gene with an N-terminal 6xHis tag into a modified pET9a plasmid via Golden Gate cloning using a BsaI restriction endonuclease to produce the pET9a-6xHis-DHFR plasmid. We chemically transformed the pET9a-6xHis-DHFR plasmid into a BL21 LOBSTR strain. Cells were grown at 37 ºC overnight shaking at 200 rpm in LB media with 50 µg/ml kanamycin, then subcultured 1/100 into 0.5 L LB media with 50 µg/ml kanamycin and grown at 37 ºC until the optical density at 600 nm was measured between 0.2 and 0.4. Protein expression was induced by the addition of 1 mM IPTG and the cells were grown at 16 ºC for 12-16 hrs. The culture was centrifuged at 6000 × g for 20 minutes and the cell pellet was frozen at -20 ºC. After several weeks, the pellet was thawed in 20 ml lysis buffer: 50 mM MOPS, 200 mM NaCl, 0.5 mM TCEP, 5 mM MgCl2, 1 mM MnCl2, 100 µM CaCl2, with a protease inhibitor cocktail (Pierce), and 2000 units/ml DNase I. The mixture was vortexed at 20 ºC for 15 minutes, then sonicated in an ice bath at 30% amplitude for 10 minutes, with a four second break following every two seconds of sonication to prevent overheating. Next, the lysate was centrifuged at 27,000 × g for 30 minutes at 4 ºC. The soluble fraction was applied to 2 ml Ni-NTA resin in 2-5 ml sample buffer (50 mM MOPS, 200 mM NaCl, 0.5 mM TCEP, pH8.0), and the mixture left to nutate overnight at 4 ºC. The mixture was then added to a 20 ml benchtop gravity column, allowed to drain and settle, and washed with 50 mL wash buffer (50 mM MOPS, 250 mM NaCl, 0.5 mM TCEP, 20 mM imidazole, pH 8.0). The protein was eluted in 15 ml of elution buffer (50 mM MOPS, 150 mM NaCl, 200 mM imidazole, pH 8.0) and concentrated to 4 mg/mL using a 15 ml ultracentrifugal filter column with a 10 KDa MWCO (MilliporeSigma). This purified stock was split into aliquots, flash frozen in liquid nitrogen, and stored at -80 ºC.

*hDHFR*

For hDHFR Rep1 and Rep2, we purchased full-length recombinant human DHFR from Abnova (catalog # P3492). The protein was diluted threefold into 50 mM HEPES buffer (pH 7.0) containing 150 mM NaCl, 1 mM TCEP, and 7.5% glycerol. It was then flash-frozen in liquid nitrogen and stored at -80 °C. Following the same Golden Gate cloning procedure used for *E. coli* DHFR, we cloned the ET9a-6xHis-hDHFR construct with the exact same gene sequence as the purchased protein and transformed it into the BL21 LOBSTR strain. The hDHFR was purified using the HEPES buffer and the same purification procedure as for *E. coli* DHFR, and used for HX/MS hDHFR Rep3 and activity assays.

*LacI*

We introduced a 6xHis-LacI1-331 gene amplified from the *E. coli* DH10B genome into a modified pET9a plasmid via Golden Gate cloning using a BsaI restriction endonuclease (NEB) to produce the pET9a-6xHis-LacI1-331 plasmid. This construct encodes a dimeric LacI without the tetramerization domain and with an N-terminal 6×-histidine (6×his) tag. We chemically transformed the pET9a-6xHis-LacI1-331 plasmid into a BL21 LOBSTR strain. Cells were grown at 37 ºC overnight shaking at 200 rpm in LB media with 50 µg/ml kanamycin, then subcultured

1/100 into 1 L LB media with 50 µg/ml kanamycin and grown at 37 ºC until the optical density at 600 nm was measured between 0.2 and 0.4. Protein expression was induced by the addition of 1 mM isopropyl ß-D-1-thiogalactopyranoside (IPTG) and the cells were grown at 16 ºC for 12-16 hrs. The culture was centrifuged at 6000 × g for 20 minutes and the cell pellet was frozen at -20 ºC. After several weeks, the pellet was thawed in 30 ml lysis buffer: 50 mM Tris, 30 mM NaCl, 5 mM MgCl2, 1 mM MnCl2, 100 µM CaCl2, 2 mM EDTA, with a protease inhibitor cocktail (Pierce), 1 mg/ml lysozyme, and 2000 units/ml DNase I (Sigma-Aldrich). The mixture was lysed at 37 ºC for 15 minutes prior to centrifugation at 27,000 × g for 30 minutes at 4 ºC. The soluble fraction was filtered and applied to 2 ml Ni-NTA resin (Thermo Fisher) in 2-5 ml wash buffer (50 mM Tris, 150 mM NaCl, 20 mM imidazole, pH 8.0), and the mixture left to nutate overnight at 4 ºC. The mixture was then added to a 20 ml benchtop gravity column, allowed to drain and settle, and washed with 50 ml of wash buffer. The protein was eluted in 15 ml of elution buffer (50 mM Tris, 150 mM NaCl, 250 mM imidazole, pH 8.0) and concentrated to 1.5 ml using a 15 ml ultracentrifugal filter column with a 10 KDa MWCO (MilliporeSigma). To remove the 6×his tag, 10 µl TEV protease (MilliporeSigma) was added to the concentrated protein solution, and the reaction was incubated at 30 ºC for 1 hour. The mixture was injected on a Bio-Rad NGC™ medium-pressure chromatography system and applied to a Cytiva S100 size-exclusion chromatography column in sample buffer (50 mM Tris, 150 mM NaCl, pH 8.0). Fractions corresponding to an A280 peak of the correct size were collected, pooled, concentrated to 3.8 mg/ml, and flash-frozen in aliquots at -80 ºC.

##### HX/MS sample preparation

$H_2O$ exchange buffer (for LacI, 50mM Tris, 150mM NaCl, pH 8.0, and for ecDHFR, 50 mM potassium phosphate, 1 mM NaCl, 10 mM beta-mercaptoethanol, pH 7.0) was lyophilized in 5 mL aliquots using Labconco FreeZone 4.5 lyophilizer for at least 24 hours, and then re-hydrated in $D_2O$ to produce deuterated exchange buffer. For the LacI dimer, the protein solution was thawed on ice and stock solutions were prepared at 7.2 µM. IPTG stocks were prepared at 10 mM IPTG and DNA stocks at 14.4 µM double-stranded.

For ecDHFR and hDHFR, the protein stocks were thawed, diluted tenfold into the analysis buffer (50 mM potassium phosphate, 1 mM NaCl, 10 mM beta-mercaptoethanol, pH 7.0), and then concentrated to 30 µM and 15 µM, respectively, using a 15 ml ultracentrifugal filter column with a 10 KDa MWCO (MilliporeSigma). For ecDHFR and hDHFR, TMP and MTX were added to separate stock vials at concentrations of 40 µM and 35 µM, respectively. To increase TMP concentration during exchange without exceeding solubility limits, the deuterated buffer for the hDHFR experiment also contained 40 µM TMP.

##### Preparation of fully deuterated control samples

Lyophilized buffer salts (see *HX/MS sample preparation*) were combined with GuHCl and re-hydrated in D2O to a concentration of 3 M to produce a denaturing exchange buffer. This was permitted to exchange for at least 5 minutes and then re-lyophilized and re-hydrated in D2O. The protein solution was thawed on ice, and then 50 µl protein solution added to 450 µl of denaturing buffer. This combination was permitted to exchange for 24 hr at 37 ºC. This solution was then manually added in place of an equivalent volume of exchange reaction in the usual HX/MS workflow (see *HX/MS data collection*), substituting a quench buffer without GuHCl to preserve composition of the quenched exchange reaction.

**HX/MS data collection**

The HX liquid handling was performed by the LEAP system (Trajan) and scheduled for automated liquid handling steps and MS injections using the Chronos 5.8.3 software (Trajan). 75 µL of exchange buffer was added to 7.5 µL of sample and mixed by pipetting. At the specified time, 75 µL from the mix was added to 75 µL of quench solution chilled to 2° C and mixed. The quench solution consisted of 3 M guanidine hydrochloride (Sigma-Aldrich) in 3% acetonitrile (Fisher Scientific) and 1% formic acid (Fisher Scientific), filtered through 0.20 µM syringe filter (Corning). All solvents used were MS-grade. 140 µL of the chilled, quenched mixture was injected into the protease column, which was chilled to 7 °C. Solvent A (3% acetonitrile, and 0.15% formic acid, v/v, both from Fisher Scientific) was pumped (UltiMate 3000, ThermoFisher Scientific) at 150 µL/min through the protease column. Peptides eluted from the protease column were trapped on a 1x10 mm C18 column (Hypersil GOLD, 3 µM pore size, ThermoFisher Scientific). For LacI and ecDHFR, we used an immobilized protease type XIII/pepsin column (FP, 1:1 w/w, 2.1 × 30 mm, NovaBioAssays), and for ecDHFR we also used an alanyl aminopeptidase/pepsin and a nepenthesin 2/pepsin column (AP/pepsin and Nep/pepsin, both 2.1 x 20 mm, Affipro). For hDHFR, we used the FP/pepsin and the Nep/pepsin columns. Full experimental details are available for ecDHFR in Table S7, hDHFR in Table S9, and LacI in Table S11.

An auxiliary pump (MX-Class, Teledyne) flowed additional Solvent A over the trap C18 at 150 µL/min. After 200 s of digesting and desalting, the LEAP system valve changed configuration so that the binary pump (Infinity II, Agilent) was in-line with the trap C18, flowing in the opposite direction. The binary pump used a gradient of Solvent A and Solvent B (96.85% acetonitrile, 0.15% formic acid) to elute peptides from a 50 x 1 mm C18 analytical column (Hypersil GOLD, 1.9 µM pore size, ThermoFisher Scientific). The elution method consisted of isocratic flow at 10% B for 1 min, a linear gradient to 40% B from 1 to 15 min, a linear gradient to 95% B from 15 to 19 min, isocratic flow at 100% B from 19 to 24 min, a linear gradient to 10% B from 24 to 30 min, and then isocratic flow at 10% B from 30 to 45 min, at a flow rate of 0.04 mL/min. Between each injection, the protease column was washed with injections of 3 M guanidine hydrochloride in 3% acetonitrile and 1% formic acid.

Eluted peptides entered the mass spectrometer (Bruker Maxis II ETD) via electro-spray ionization, where full-scan mass spectra were collected in positive ion mode from 100-2250 m/z with a spectra rate of 1.0 Hz, 500 V end plate offset, 4200 V capillary, 1.7 bar nebulizer, 8 L/min dry gas, and 180° C dry temp. We also ran tandem mass spectrometry (MS/MS) experiments for each sample with the same full MS settings as described above with the exception of the spectra rate of 1.20 Hz and the 200° C dry temp. We used Bruker Compass Hystar 5.1 software to acquire the data.

#### HX/MS data analysis

We used Compass DataAnalysis 5.1 (Bruker) to identify and deconvolute peptides and produce .mgf format compound lists for PIGEON.

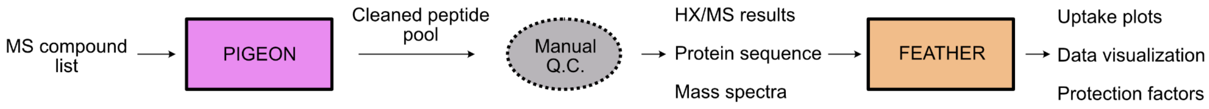

#### Peptide deconvolution by PIGEON

It is not required for the PIGEON-FEATHER protocol, but to make sure that all the peptides were correct, we used HDExaminer 3.3 (Trajan) to identify the mass spectrum for each time point and peptide in the experiment and then manually checked each mass spectrum to make sure the automatically selected retention times matched the same chromatographic peak for each time point. To do this, we first used the PIGEON-exported .csv peptide pool as the peptide source in HDExaminer and extracted actual exchange times from the LEAP run logs and corrected our recorded exchange times with these values using an in-house Python script. Following this manual quality control step, we exported the peptide pool results and the pool spectra for use in FEATHER. We also made a range list of peptides to either include or exclude in FEATHER. A range list example is included in the PIGEON-FEATHER GitHub repository.

#### Preparation of structural priors for FEATHER

For LacI, we used the experimental structures solved by Daber *et al.* (IPTG state, PDB ID 2P9H[23]), Bell and Lewis (DNA-bound state, PDB ID 1EFA), and Lewis *et al.* (APO state, PDB ID 1LBI[26]). These were relaxed using Rosetta with the REF2015 energy function and solvated using Rosetta-ECO. For the IPTG structure (solved at 2 Å resolution), several low-b factor water molecules were visible in the binding pocket. To prepare the structural prior, all crystallographic water molecules with b factors below 30 were retained, and we applied Rosetta-ECO to the partially solvated structure to model the remaining water molecules. Our script for preparing structural models for LacI is here:

```
import pyrosetta
import sys
import os
from subprocess import Popen, PIPE
from glob import glob

pyrosetta.init()

class FastRelaxer:
    def __init__(
        self, scorefxn=None, constrain_relax=False, task_factory=None,
**kwargs
    ):
        self.scorefxn = scorefxn
```

```
        self.kwargs = kwargs
        self.constrain_relax = constrain_relax
        self.task_factory = task_factory
    def configure_relax(self):
        if self.scorefxn is None:
            self.scorefxn = pyrosetta.get_fa_scorefxn()
        self.relax =
pyrosetta.rosetta.protocols.relax.FastRelax(self.scorefxn)
        if self.constrain_relax:
            self.relax.constrain_relax_to_start_coords(True)
            self.relax.coord_constrain_sidechains(True)
        if self.task_factory is not None:
            self.relax.set_task_factory(self.task_factory)
    def run(self, pose, output_path=None, nstruct=1):
        if output_path is None:
            raise ValueError("output_path must be provided")
        all_poses = []
        for i in range(nstruct):
            pose_i = pose.clone()
            self.configure_relax()
            self.relax.apply(pose_i)
            pose_i.dump_pdb(output_path[:-4] + f"_{i}.pdb")
            all_poses.append(pose_i)
        best_pose = all_poses[
            np.argmin([self.scorefxn.score(pose) for pose in
all_poses])
        ]
        best_pose.dump_pdb((output_path[:-4] + f"_best.pdb"))
        return best_pose

def relax_a_structure(pdb_file, out_folder):
    pyrosetta_init()
    relaxer = FastRelaxer(constrain_relax=True)
    relaxer.configure_relax()
    pose = pose_from_md_structure(pdb_file)
    base_pdb = pdb_file.split("/")[-1].split(".")[0]
    relaxed_pdb = f"{out_folder}/{base_pdb}_relaxed.pdb"
    relaxer.run(pose, relaxed_pdb, nstruct=5)

def runUnsatScript(pose: pyrosetta.Pose):
    xml: pyrosetta.rosetta.protocols.rosetta_scripts.XmlObjects = (

pyrosetta.rosetta.protocols.rosetta_scripts.XmlObjects.create_from_stri
ng(f"""
    <ROSETTASCRIPTS>
    <SCOREFXNS>
    <ScoreFunction name="r15" weights="ref2015"/>
    </SCOREFXNS>
    <TASKOPERATIONS>
```

```
    <RestrictToRepacking name="RestrictToRepacking" />
    </TASKOPERATIONS>
    <RESIDUE_SELECTORS>
    <Chain name="chAll" chains="A,B,X"/>
    </RESIDUE_SELECTORS>
    <FILTERS>
    </FILTERS>
    <MOVERS>
    <ExplicitWaterMover name="solvate" mode="replace" gen_fixed="1"
scorefxn="r15" task_operations="RestrictToRepacking"/>
    <PackRotamersMover name="pack" scorefxn="r15"
task_operations="RestrictToRepacking"/>
    </MOVERS>
    <PROTOCOLS>
    <Add mover_name="solvate"/>
    <Add mover_name="pack"/>
    </PROTOCOLS>
    <OUTPUT/>
    </ROSETTASCRIPTS>
    """)
    )
    xml.get_mover("solvate").apply(pose)
    xml.get_mover("pack").apply(pose)

def runScript(files: list):
    # pyrosetta.init("-mute all")
    pyrosetta.init()
    for pdb in files:
        # params_list =
pyrosetta.Vector1(["/ifs/scratch/home/master/TPB/2p9h_test/test1/test2/
IPT.params"])
        pose = pyrosetta.Pose()
        # pyrosetta.generate_nonstandard_residue_set(pose, params_list)
        pyrosetta.pose_from_file(pose, pdb)
        runUnsatScript(pose)
        pose.dump_pdb(f"{pdb[:-4]}_solvated.pdb")

if __name__ == "__main__":
    WDIR = "LacI"
    RELAX_DIR = f"{WDIR}/01_relax"
    relax_a_structure("1EFA_DNA.pdb", RELAX_DIR)
    runScript([f"{RELAX_DIR}/1EFA_DNA_relaxed_best.pdb"])
```

##### PIGEON error calculations

For all HX/MS datasets, using the ‘Keep’ option, we compared the mean score, the percentage of the protein sequence that was not covered by the data (score 0), and the root-mean-square (RMS) ppm error of the *m*/*z* values after correcting for calibration error at each stage of analysis.

Each of these values is listed for each dataset in Table S1, with RMS ppm errors for the following stages of PIGEON analysis: before removing any peptides (*input*) (Extended Data Fig. 1e, i, 1f, Fig. S1a); in the high scoring dataset used for the mass error fit (*high score*) (Extended Data Fig. 1e, ii, Fig. S1b); after re-thresholding by ppm error (*mz threshold*) (Extended Data Fig. 1e, ii, Fig. S1c); after removing redundant peaks for each peptide (*single match*) (Extended Data Fig. 1e, ii, Fig. S1d), the final cleaned dataset (*clean*) (Extended Data Fig. 1e, v, h), and discarded degenerate peptides (Extended Data Fig. 1e, iii, iv, g).

##### FEATHER error calculations

The uncertainty of the log(PF) values was calculated by analyzing their posterior distributions. These distributions were derived by combining results from multiple posterior analyses, each of which used bootstrapping techniques where 90% of the peptides were randomly resampled from the entire dataset. Additionally, the average fitting error including both centroid and isotopic mass envelope for all peptides covering a residue was calculated. Our analysis indicated that the uncertainty of the posterior distributions, when combined with multiple bootstrapping runs, yields the highest correlation. This approach was therefore adopted as the primary method of error estimation for our analysis. These residue-level bootstrap errors are included as error bars in PIGEON-FEATHER-generated -log($k_{ex}$) vs. residue plots.

##### DHFR activity assay

The DHFR activity assay was adapted from previous publications.[27,28] We performed the assay at room temperature using a Thermo Scientific NanoDrop 2000 spectrophotometer. The reaction was conducted in a buffer containing 50 mM HEPES, 150 mM NaCl, 1 mM TCEP, and 7.5% glycerol, pH 7.0. Freshly purified DHFR (10 µL from a 1 µM stock) was mixed with dihydrofolic acid (DHF, MilliporeSigma) (40 µL from an 80 µM stock) and incubated for 5 minutes. The reaction was initiated by adding NADPH (20 µL from a 500 µM stock), and the rate of NADPH consumption during the conversion of dihydrofolate to tetrahydrofolic acid (THF) was monitored by measuring the absorbance at 340 nm every 2 seconds over the initial 30 seconds. In the final reaction mixture, the concentrations of each compound were: 100 nM DHFR, 100 µM NADPH, 32 µM DHF, and a specific amount of the inhibitor (37.5-1200 nM).

### 6. References

1. Bai, Y., Milne, J. S., Mayne, L. & Englander, S. W. Primary structure effects on peptide group hydrogen exchange. *Proteins Struct. Funct. Bioinforma.* **17**, 75–86 (1993).
2. Pace, C. N. [14]Determination and analysis of urea and guanidine hydrochloride denaturation curves. in *Methods in Enzymology* vol. 131 266–280 (Academic Press, 1986).
3. Shirley, B. A. Urea and Guanidine Hydrochloride Denaturation Curves. in *Protein Stability and Folding: Theory and Practice* (ed. Shirley, B. A.) 177–190 (Humana Press, Totowa, NJ, 1995). doi:10.1385/0-89603-301-5:177.
4. Bai, Y., Sosnick, T. R., Mayne, L. & Englander, S. W. Protein Folding Intermediates: Native-State Hydrogen Exchange. *Science* **269**, 192–197 (1995).
5. Chamberlain, A. K., Handel, T. M. & Marqusee, S. Detection of rare partially folded molecules in equilibrium with the native conformation of RNaseH. *Nat. Struct. Biol.* **3**, 782–787 (1996).
6. Hollien, J. & Marqusee, S. Structural distribution of stability in a thermophilic enzyme. *Proc. Natl. Acad. Sci.* **96**, 13674–13678 (1999).
7. Manna, M. S. *et al.* A trimethoprim derivative impedes antibiotic resistance evolution. *Nat. Commun.* **12**, 2949 (2021).
8. Vendruscolo, M., Paci, E., Dobson, C. M. & Karplus, M. Rare Fluctuations of Native Proteins Sampled by Equilibrium Hydrogen Exchange. *J. Am. Chem. Soc.* **125**, 15686–15687 (2003).
9. Alford, R. F. *et al.* The Rosetta All-Atom Energy Function for Macromolecular Modeling and Design. *J. Chem. Theory Comput.* **13**, 3031–3048 (2017).
10. Bystroff, C. & Kraut, J. Crystal structure of unliganded Escherichia coli dihydrofolate reductase. Ligand-induced conformational changes and cooperativity in binding. *Biochemistry* **30**, 2227–2239 (1991).
11. Sawaya, M. R. & Kraut, J. Loop and Subdomain Movements in the Mechanism of Escherichia coli Dihydrofolate Reductase:  Crystallographic Evidence,. *Biochemistry* **36**, 586–603 (1997).
12. Bhabha, G. *et al.* A Dynamic Knockout Reveals That Conformational Fluctuations Influence the Chemical Step of Enzyme Catalysis. *Science* **332**, 234–238 (2011).
13. Abramson, J. *et al.* Accurate structure prediction of biomolecular interactions with AlphaFold 3. *Nature* 1–3 (2024) doi:10.1038/s41586-024-07487-w.
14. Rodriguez, D. C. P., Weber, K. C., Sundberg, B. & Glasgow, A. MAGPIE: An interactive tool for visualizing and analyzing protein–ligand interactions. *Protein Sci.* **33**, e5027 (2024).
15. RCSB PDB - 2W3A: HUMAN DIHYDROFOLATE REDUCTASE COMPLEXED WITH NADPH AND TRIMETHOPRIM. https://www.rcsb.org/structure/2W3A.
16. Cody, V., Luft, J. R. & Pangborn, W. Understanding the role of Leu22 variants in methotrexate resistance: comparison of wild-type and Leu22Arg variant mouse and human dihydrofolate reductase ternary crystal complexes with methotrexate and NADPH. *Acta Crystallogr. D Biol. Crystallogr.* **61**, 147–155 (2005).
17. Gangjee, A. *et al.* Structure-Based Design and Synthesis of Lipophilic 2,4-Diamino-6-

Substituted Quinazolines and Their Evaluation as Inhibitors of Dihydrofolate Reductases and Potential Antitumor Agents. *J. Med. Chem.* **41**, 3426–3434 (1998).
18. McTigue, M. A., Davies, J. F. I., Kaufman, B. T. & Kraut, J. Crystal structure of chicken liver dihydrofolate reductase complexed with NADP+ and biopterin. *Biochemistry* **31**, 7264–7273 (1992).
19. Bhabha, G. *et al.* Divergent evolution of protein conformational dynamics in dihydrofolate reductase. *Nat. Struct. Mol. Biol.* **20**, 1243–1249 (2013).
20. Cody, V., Luft, J. R. & Pangborn, W. Understanding the role of Leu22 variants in methotrexate resistance: comparison of wild-type and Leu22Arg variant mouse and human dihydrofolate reductase ternary crystal complexes with methotrexate and NADPH. *Acta Crystallogr. D Biol. Crystallogr.* **61**, 147–155 (2005).
21. Bell, C. E. & Lewis, M. A closer view of the conformation of the Lac repressor bound to operator. *Nat. Struct. Biol.* **7**, 209–214 (2000).
22. Glasgow, A. *et al.* Ligand-specific changes in conformational flexibility mediate long-range allostery in the lac repressor. *Nat. Commun.* **14**, 1179 (2023).
23. Daber, R., Stayrook, S., Rosenberg, A. & Lewis, M. Structural analysis of lac repressor bound to allosteric effectors. *J. Mol. Biol.* **370**, 609–619 (2007).
24. Saltzberg, D. J. *et al.* A Residue-Resolved Bayesian Approach to Quantitative Interpretation of Hydrogen–Deuterium Exchange from Mass Spectrometry: Application to Characterizing Protein–Ligand Interactions. *J. Phys. Chem. B* **121**, 3493–3501 (2017).
25. Masson, G. R. *et al.* Recommendations for performing, interpreting and reporting hydrogen deuterium exchange mass spectrometry (HDX-MS) experiments. *Nat. Methods* **16**, 595–602 (2019).
26. Lewis, M. *et al.* Crystal Structure of the Lactose Operon Repressor and Its Complexes with DNA and Inducer. *Science* **271**, 1247–1254 (1996).
27. Krucinska, J. *et al.* Structure-guided functional studies of plasmid-encoded dihydrofolate reductases reveal a common mechanism of trimethoprim resistance in Gram-negative pathogens. *Commun. Biol.* **5**, 1–14 (2022).
28. Srinivasan, B., Rodrigues, J. V., Tonddast-Navaei, S., Shakhnovich, E. & Skolnick, J. Rational Design of Novel Allosteric Dihydrofolate Reductase Inhibitors Showing Antibacterial Effects on Drug-Resistant Escherichia coli Escape Variants. *ACS Chem. Biol.* **12**, 1848–1857 (2017).
